# sf-pediatric: A robust and age-adaptable end-to-end pipeline for pediatric diffusion MRI

**DOI:** 10.64898/2026.01.19.700454

**Authors:** Anthony Gagnon, Arnaud Boré, Alex Valcourt Caron, Manon Edde, Stanislas Thoumyre, Jean-François Lepage, Ardesheer Talati, Jonathan Posner, Annie Ouellet, Marie A. Brunet, Larissa Takser, François Rheault, Maxime Descoteaux

**Author notes:** **Corresponding author:** Anthony Gagnon, MSc, Département de Pédiatrie, Faculté de Médecine et des Sciences de la Santé, Université de Sherbrooke, 3001, 12^ième^ Avenue Nord, Sherbrooke, QC, Canada, J1H 5N4. **Competing Interests:** Author MD is a co-founder and shareholder at Imeka Solutions Inc (www.imeka.ca). All other authors declare no financial or non-financial competing interests.

## Abstract

Diffusion MRI (dMRI) provides a powerful, non-invasive window into white matter (WM) development. Yet, most existing processing pipelines are not well-suited to the rapidly evolving neurophysiology of the pediatric brain. Here, we introduce sf-pediatric, a scalable, end-to-end, age-adaptable dMRI pipeline that integrates normative models of brain diffusivities to enable optimal subject-specific analysis from birth through 18 years old. Leveraging normative trajectories derived from nearly 2,000 participants from six cohorts, sf-pediatric dynamically calibrates diffusion priors, template selection, segmentation, and WM atlases based on the subject’s age. By incorporating automatic quality control into a portable, tested, containerized, open-access, and press-button framework across computing environments, sf-pediatric provides a robust pipeline for large-scale pediatric dMRI studies. We validated this approach by showing improved local modeling and cortical fanning while preserving reproducibility and the ability to derive brain-behavior relationships. Additionally, we demonstrated robust recovery of known developmental trajectories of WM microstructure and connectome-derived network organization.

## Introduction

Infant brain development has been increasingly studied in recent years, stemming partly from the drastic increase in prevalence of neurodevelopmental disorders^1,2^, which are thought to originate from atypical early developmental processes^3^. One method to investigate and assess developmental changes is the use of non-invasive methods, such as diffusion MRI (dMRI), which allows the investigation of white matter (WM) macro- and microstructural changes across the developmental period. DMRI relies on the diffusion properties of water molecules and their restriction within the brain^4^. Using either simple^5^ or more complex models^6,7^, researchers can reconstruct the organization of WM across the lifespan, offering powerful insights into the structural connectivity of the human brain. While conceptually sound, faithfully reconstructing the WM organization during development remains an active research problem in the dMRI field^8^.

Current techniques for processing adult diffusion MRI acquisitions cannot be directly applied to infant data due to the rapid neurophysiological changes that occur in the first few years of life^3,9–11^. While those changes are mainly apparent in the first year^11^, they continue to affect diffusion properties and signal intensities/contrasts throughout development^12^. For example, diffusion properties will be significantly affected by the lack of myelin at younger ages, but will rapidly evolve as myelination proceeds^12,13^. This impacts simple processing steps such as brain extraction, modality registration, tissue segmentation, and tractography, which become unstable and require specialized settings or approaches to perform well on pediatric data^3,9^. This also impacts how brain templates can be integrated into image processing. Since the brain is rapidly changing, we cannot use a single brain template for the complete developmental age range. In practice, this means a processing pipeline should be able to assign the optimal brain template for a single subject. This is also true for WM atlases, and even more so for advanced diffusion models such as the Neurite Orientation Dispersion and Density Imaging (NODDI)^14^, which rely on water diffusivity priors (mm^2^/s) across different brain tissues. Other MRI modalities have already proposed pediatric-tailored solutions to handle the varying signal intensities/contrasts^15,16^. Similarly, to enable large-scale studies of WM brain development, a pediatric-tailored dMRI pipeline that accounts for those rapidly changing neurophysiological properties is needed.

Existing pipelines offer specific configurations for infant data but are either suboptimal, limited to a particular age range, poorly scalable to large numbers of participants, or do not support all possible analysis types (e.g., connectomics analysis)^17–20^. Growing pediatric initiatives are generating tens of thousands of dMRI acquisitions yearly^21,22^, reinforcing the need for a scalable processing solution. Additionally, there are limited end-to-end solutions that require no manual input, enabling users to go from raw data acquisition to statistics-ready datasets while providing integrated QC reports. While manual input might not be an issue for small datasets, the risk of human error scales exponentially with the number of participants, especially when working with remote computing resources. Indeed, cloud-based computing resources or high-performance computing (HPC) clusters are more accessible to researchers worldwide and are the backbone of large-scale neuroimaging studies. Concurrently, containerization technologies have made software versions more manageable and represent a step toward reproducible processing^23,24^. Those valuable resources force new neuroimaging pipelines to integrate containerization and support a wide range of computing architectures beyond local computers, increasing their technological complexity.

Normative models of brain development, including WM development, are increasingly accessible, offering significant translational opportunities for clinical practice^25,26^. Those normative curves of expected diffusivities in the brain throughout development provide valuable insights that could be leveraged in dMRI image processing but are currently excluded from the existing processing solutions. As such, we introduce sf-pediatric, a next-generation, scalable, end-to-end, age-adaptable diffusion MRI pipeline for pediatric data that builds on knowledge gained from normative modeling to provide age-adaptable priors of the brain’s diffusivities while dynamically interacting with brain templates and WM atlases. To validate our age-adaptable priors, we evaluated their effects on local modeling, cortical fanning, and the ability to derive brain-behavior relationships when compared to reference young-adult priors. To demonstrate the pipeline’s usability across multiple contexts, we derived normative curves for WM microstructure and graph theory network metrics across the developmental age range. To our knowledge, sf-pediatric is the first portable, containerized, open-access, press-button, and age-adaptable pipeline to actively leverage normative data derived from nearly 2,000 participants to improve dMRI processing. Combined with a modular architecture based on nf-neuro^27^, ensuring thoroughly tested modules, and the portable Nextflow workflow manager^28^, users can select four main workflows (e.g., profiles) based on their research aims and obtain results from advanced diffusion models ready for analysis, making sf-pediatric a viable solution for large-scale pediatric studies.

## Results

### Leveraging normative models for improved signal reconstruction

The scope of normative trajectories of brain development has primarily been focused on clinical applications. However, they also provide valuable insights for image processing by indicating normative expectations for the brain’s neurophysiology, specifically during the first year of life^3^. To leverage this information, we derived normative models of the brain’s diffusivities from dMRI acquisitions across six pediatric cohorts spanning the 0-18-year-old range (Figure 1a and Supplementary Tables 1-2). From the diffusion tensors, we extracted fractional anisotropy (FA), axial diffusivity (AD), and radial diffusivity (RD) from single fiber populations as an indicator of anisotropy, while mean diffusivity (MD) was computed in voxels representing cerebrospinal fluid (CSF) as an indicator of isotropic diffusivity (see Methods). For each metric, we modeled the normative curves using a generalized additive model for location, shape, and size (GAMLSS)^29^, enabling estimation of age-related changes while accounting for study effects (see Methods).

**Figure 1.**
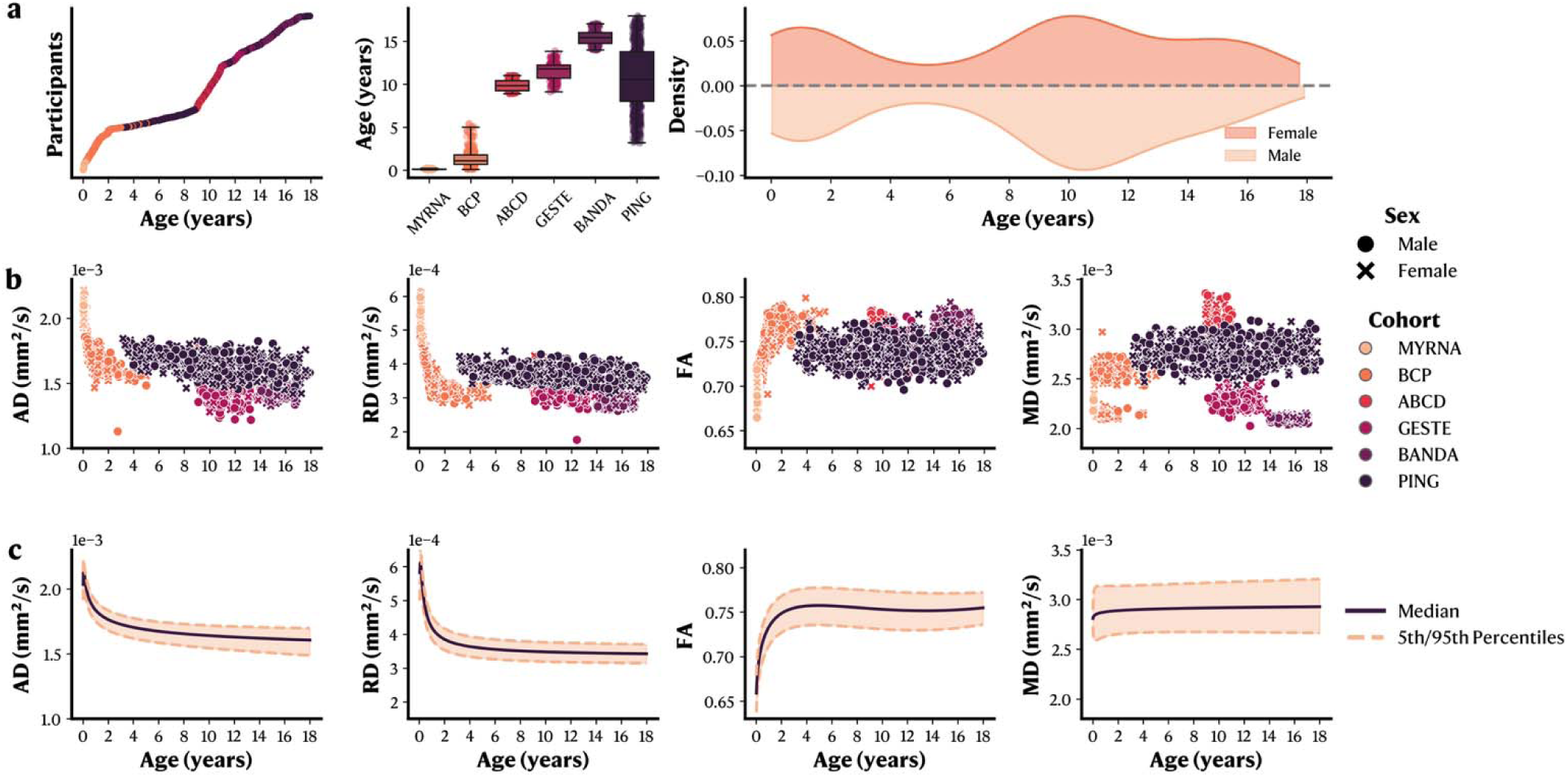
Normative curves of the diffusivities across the developmental age range for custom age-adaptable diffusivity priors. **a.** Cohorts’ demographics and age range covered by the selected cohorts. **b.** Scatter plots representing the raw data distribution across all cohorts. AD, RD, and FA were extracted from single fiber populations, whereas MD was extracted from voxels within the ventricles. **c.** Normative curve for each diffusion metric with the 5^th^ and 95^th^ percentiles. AD: Axial Diffusivity. RD: Radial Diffusivity. FA: Fractional Anisotropy. MD: Mean Diffusivity. MYRNA: Maternal and Youth Research on Neurodevelopment and behAvior. BCP: Baby Connectome Project. ABCD: Adolescent Brain Cognitive Development. GESTE: GESTation and Environment. BANDA: Boston Adolescent Neuroimaging of Depression and Anxiety. PING: Pediatric Imaging, Neurocognition, and Genetics.

Normative curves showed expected trajectories throughout development (Figure 1b-c), with both RD and AD showing a steep decrease in the first two years of life, while FA showed an inverse pattern, followed by stabilization for the rest of the developmental period. MD values in CSF showed a relatively stable trajectory from birth to adulthood, suggesting that isotropic diffusivities do not change significantly throughout development. To leverage those normative curves within sf-pediatric, we approximated the median data from the GAMLSS models. Those equations enable direct use of the acquired knowledge from normative trajectories for both large samples and smaller datasets, while retaining most of the information (R^2^ = 0.88–1.00, see Methods, and Supplementary Figures 1-4).

### Age-adaptable priors yield better local modeling and cortical fanning

To evaluate the impact of the age-adaptable priors on results derivatives relative to the baseline young-adult priors, we computed an average diffusivity from the three directions forming the fiber response function (see Methods). A higher average diffusivity makes a sharper fiber-oriented distribution function (fODF)^30,31^. As expected, we observed a higher average diffusivity in younger participants, which decreases exponentially through development toward the reference young-adult prior (Figure 2a). Qualitative evaluation of the resulting fODF showed larger, sharper lobes when using the age-adaptable priors than with the young-adult reference, particularly in younger participants, while older participants showed more subtle differences (Figure 2b). One challenge in tractography with young participants is properly reconstructing the cortical fanning of WM bundles, partly due to how crossing fibers are modeled^9^. To evaluate the impact of the improved fODF on the streamline reconstruction, we compared the cortical surface area of the corticospinal tract (CST) using either the young-adult priors or the custom age-adaptable priors using a paired one-tailed t-test (see Methods). For both the left and right CST, introducing the age-adaptable priors returned a higher cortical surface area, suggesting a better cortical fanning and an improved tractography coverage, compared to the young-adult priors (left CST; *t* = -16.09, *p* < 0.001 and right CST; *t* = -14.10, *p* < 0.001) (Figure 2c-d). Using test-retest data from an external, unseen cohort (n=83 participants aged 5 to 8 years), we further assessed whether the addition of age-adaptable priors would significantly impact the reproducibility of the extracted bundles. Comparing the weighted dice score on the CST across test-retest acquisitions (see Methods), we found no significant differences before and after the inclusion of age-adaptable priors (Supplementary Figure 5), suggesting that these priors improve cortical fanning without compromising reproducibility.

**Figure 2.**
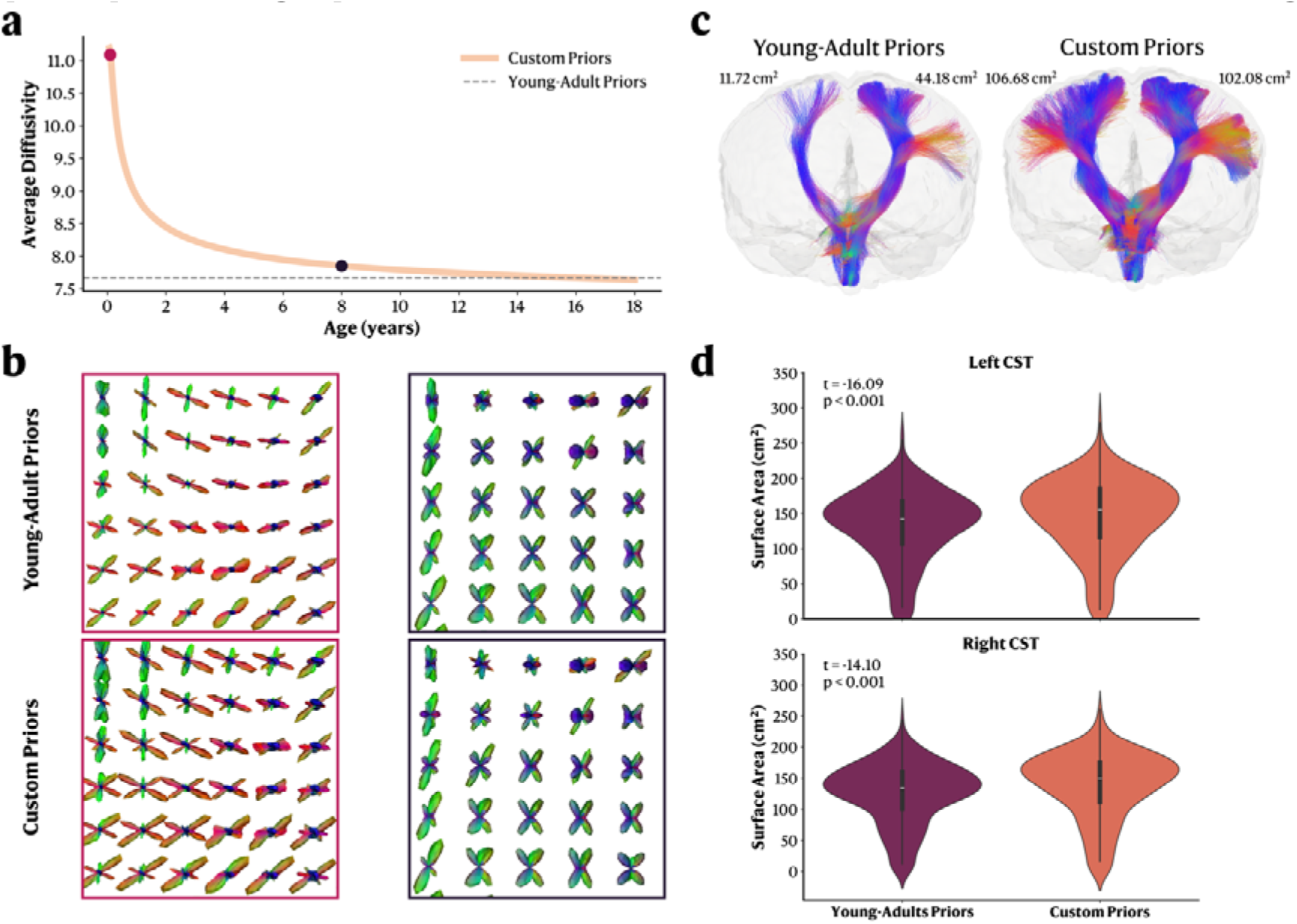
Age-adaptable intelligent pipelines allow for better diffusivity priors, and thus better fiber-oriented distribution function (fODF) calibration, which leads to sharper fODFs and improved tractography coverage of WM anatomy. **a.** Average diffusivity across development compared to the fixed average diffusivity of the young-adult priors. **b.** Visual representation of the fODF using the young-adult priors (top row) and custom age-adaptable priors (bottom row). Red squares (left column) represent a participant aged 1 month, and black squares (second column) represent a participant aged 8 years old. **c.** Visual representation of the corticospinal track in a 2-month-old subject using young-adult priors and custom priors. **d.** Cortical surface area of the corticospinal track using either young-adult priors or custom priors in the Baby Connectome Project. t: t-statistic from a one-tailed paired sample t-test. p: p-value.

### Age-adaptable priors do not affect brain-behavior associations

To further understand the impact of introducing age-adaptable priors on the ability to derive brain-behavior relationships, we conducted multivariate regression analyses to explore the association between the fiber density of the arcuate fasciculus (AF), previously linked to language capabilities^32^, and both receptive and expressive language scores in participants from a single cohort. By conducting regression analyses on the same participants using the young-adult priors and the age-adaptable custom priors, we aim to assess how these priors affect the ability to detect effect sizes in a known brain-behavior relationship. As a result, when controlling for age and sex, we observed no meaningful changes in effect sizes following the introduction of age-adaptable priors, suggesting they do not affect the ability to derive a brain-behavior relationship (Figure 3, Supplementary Table 3). To further ensure that our age-adaptable priors would not introduce spurious associations, we repeated the analysis using behavioral scores (internalization and externalization) and the fixel-based AFD on the superior longitudinal fasciculus (SLF). As expected, results were nearly identical when using the young-adult priors compared to the age-adaptable priors, suggesting they do not introduce artificial brain-behavior relationships (Supplementary Table 4 and Supplementary Figure 6).

**Figure 3.**
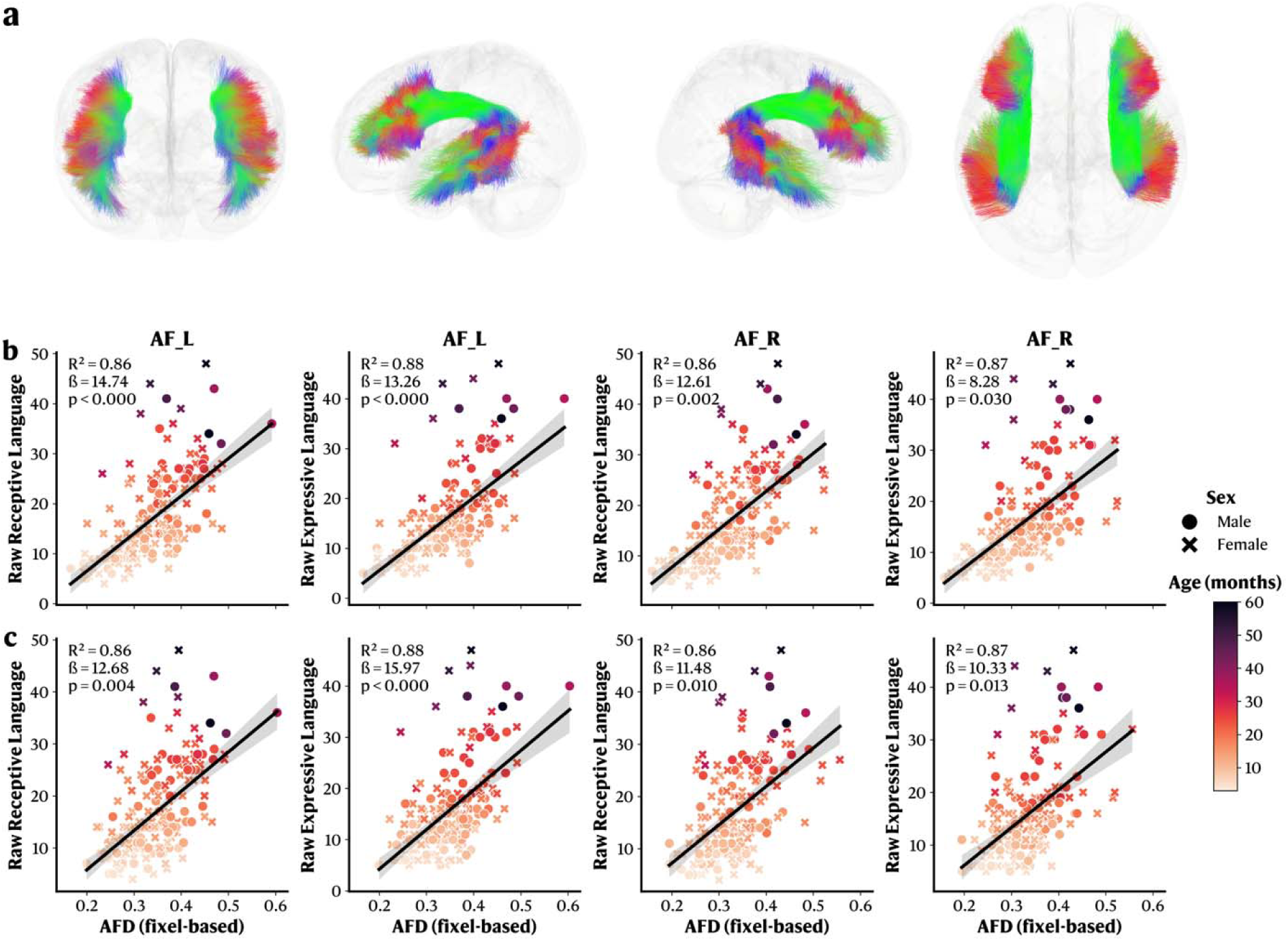
Age-adaptable priors do not break the ability to derive brain-behavior relationships. **a.** Visual representation of the left and right AF in a glass brain. **b-c.** Multivariate regression results between fixel-based average fiber density (AFD Fixel) on the arcuate fasciculus (AF) and raw language score in the BCP cohort before and after the inclusion of the custom age-adaptable priors. **b.** Scatter plots representing the association between raw language scores and AFD Fixel on both the left and right AF using the **young-adult priors**. **c.** Scatter plots representing the association between raw language scores and AFD Fixel on both the left and right AF using the **age-age-adaptable priors.** ß: Beta coefficients for the AFD Fixel variable. *p*: *p*-value for the AFD fixel variable.

### Based on a community-driven module library for robust, reproducible, and containerized workflows

The sf-pediatric pipeline is built on nf-neuro^27^, a community-hosted open-source collection of thoroughly tested, containerized, and reproducible Nextflow modules. Based on the success of nf-core^33^, a community-developed set of guidelines for standardized Nextflow modules widely adopted in bioinformatics, nf-neuro includes robust processing steps, or building blocks, from multiple state-of-the-art neuroimaging software, such as FreeSurfer^34^, ANTs^35^, scilpy^36^, FSL^37^, and MRtrix^38^ that are importable within external Nextflow pipelines. Those building blocks have designated Docker^23^ containers, making them self-contained, and are rigorously tested to ensure reproducible results. Additionally, nf-neuro provides sets of subworkflows, e.g., a chain operation consisting of multiple modules, representing standard, community-accepted preprocessing workflows. This separation between the module library and the pipeline offers a unique advantage by enabling easy continuous development and integration on one side while ensuring production-quality on the other. All code is openly available on GitHub (https://github.com/scilus/sf-pediatric.git), enabling users to propose new features or raise issues, along with a thorough documentation website to facilitate widespread adoption.

### Leveraging BIDS conventions for an age-adaptable and standardized workflow

To further the adoption of conventions in the neuroimaging field, sf-pediatric adheres to the BIDS conventions for both input and output datasets^39^. This enforced BIDS structure allows the pipeline to handle multiple acquisition schemes (e.g., complete AP-PA acquisition or single PA acquisition with single reverse-encoded b0) while also gaining access to each subject’s metadata. This access to metadata is the core principle behind sf-pediatric age-adaptable features. Using the BIDS structure, the pipeline extracts each subject’s age to adapt the processing steps to the data, select the tissue segmentation template and WM atlas, and derive the appropriate diffusivity priors (Figure 4). To preserve consistency between the enforced input structure and output files, sf-pediatric also organizes its output in a BIDS-compliant derivative folder, facilitating the integration with additional BIDS apps. sf-pediatric organizes itself around four main workflows: 1) anatomical and DWI processing, including signal reconstruction using age-adaptable priors, tissue segmentation using age-tailored brain templates, inter-modality registration, and tractography, 2) cortical/subcortical segmentation using age-optimal methods, 3) whole-brain connectome analysis leveraging the age-adaptable priors for optimal streamline filtering, and 4) automatic virtual WM bundle dissection using age-tailored WM atlases combined with tractometry. Given the wide range of possible profiles (Figure 4), sf-pediatric can be applied to various studies assessing WM development across pediatric samples, regardless of the research question.

**Figure 4.**
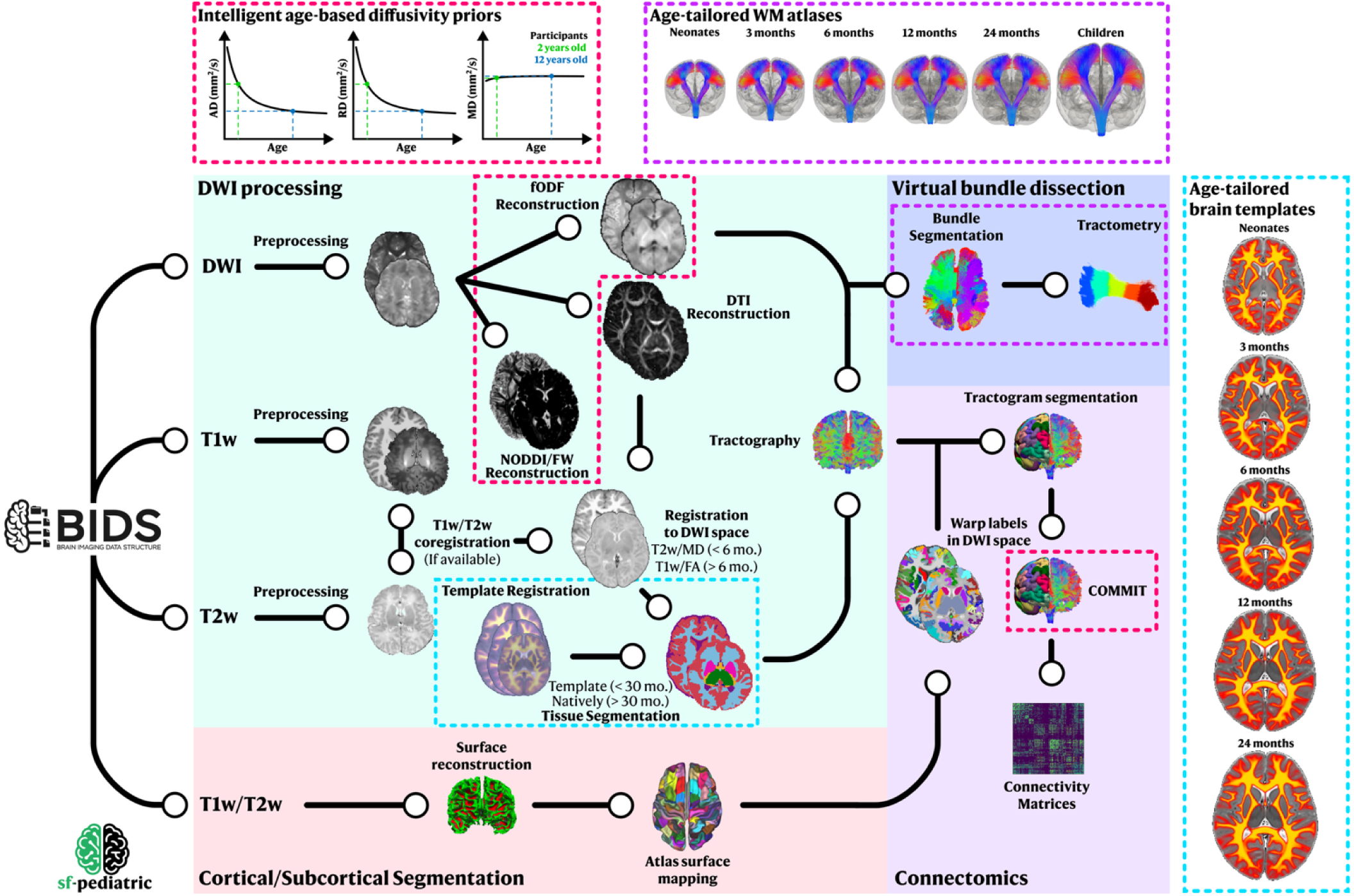
sf-pediatric pipeline schema. DWI: Diffusion weighted imaging. fODF: Fiber-oriented distribution function. DTI: Diffusion tensor imaging. PFT: Particle filter tracking. BET: Brain extraction. FW: Freewater Elimination. NODDI: Neurite Orientation Dispersion and Density Imaging. MD: Mean diffusivity. FA: Fractional anisotropy. DK: Desikan-Killiany atlas. COMMIT: Convex Optimization for Microstructure-informed Tractography.

### Subject-specific and population-wide reports ensure easy quality control

Quality control (QC) is an essential step in neuroimaging studies, as processing errors or acquisition artifacts can significantly affect downstream analyses and introduce bias. To facilitate this step, sf-pediatric leverages MultiQC^40^ to generate a global report for population-wide QC and subject-specific QC reports. The population-wide report includes distributional data over all participants, showing violin plots of maximal movement between two acquired directions, number of streamlines in the tractogram, the amount of WM covered by the tractogram, the number of extracted bundles, and cortical/subcortical region volumes. To easily pinpoint potential outliers, sf-pediatric automatically assigns a pass, warning, or fail label to each subject for each step described. Flagged subjects can then be further investigated in their own subject-specific report, which includes visual QC of key processing steps such as TOPUP/eddy, framewise displacement, metric maps, anatomical registration in diffusion space, tissue segmentation, tracking coverage, and cortical/subcortical segmentation. In all reports, the pipeline includes a methods boilerplate, along with software versions used and pipeline parameters, that is dynamically adapted to the pipeline execution parameters, enabling an easy, detailed description of the processing steps applied to the data. Example boilerplate, population-wide, and subject-specific reports are available in Supplementary Files 1, 2, and 3, respectively.

### sf-pediatric reproduces the microstructural evolution of WM bundles across the developmental age range

As shown in Figure 4, multiple processing profiles can be selected, enabling different types of analysis on diffusion data (all profiles are described in depth in the Methods section). One of which consists of extracting major WM bundles and deriving per-bundle metrics of WM microstructure. To validate that our pipeline yielded consistent results across the entire developmental age range, we performed bundle extraction and tractometry on participants from all included cohorts. We then modeled AD, RD, MD, and FA as functions of age using GAMLSS models^29^, while controlling for sex and cohort effects. For all bundles, AD, RD, and MD showed an exponential-like decrease with age, while FA showed an inverse pattern, consistent with multiple prior studies^11,25,41–43^. An example of derived trajectories is shown in Figure 5 for the posterior frontal part of the corpus callosum; the remaining bundles are shown in Supplementary Figures 7-37.

**Figure 5.**
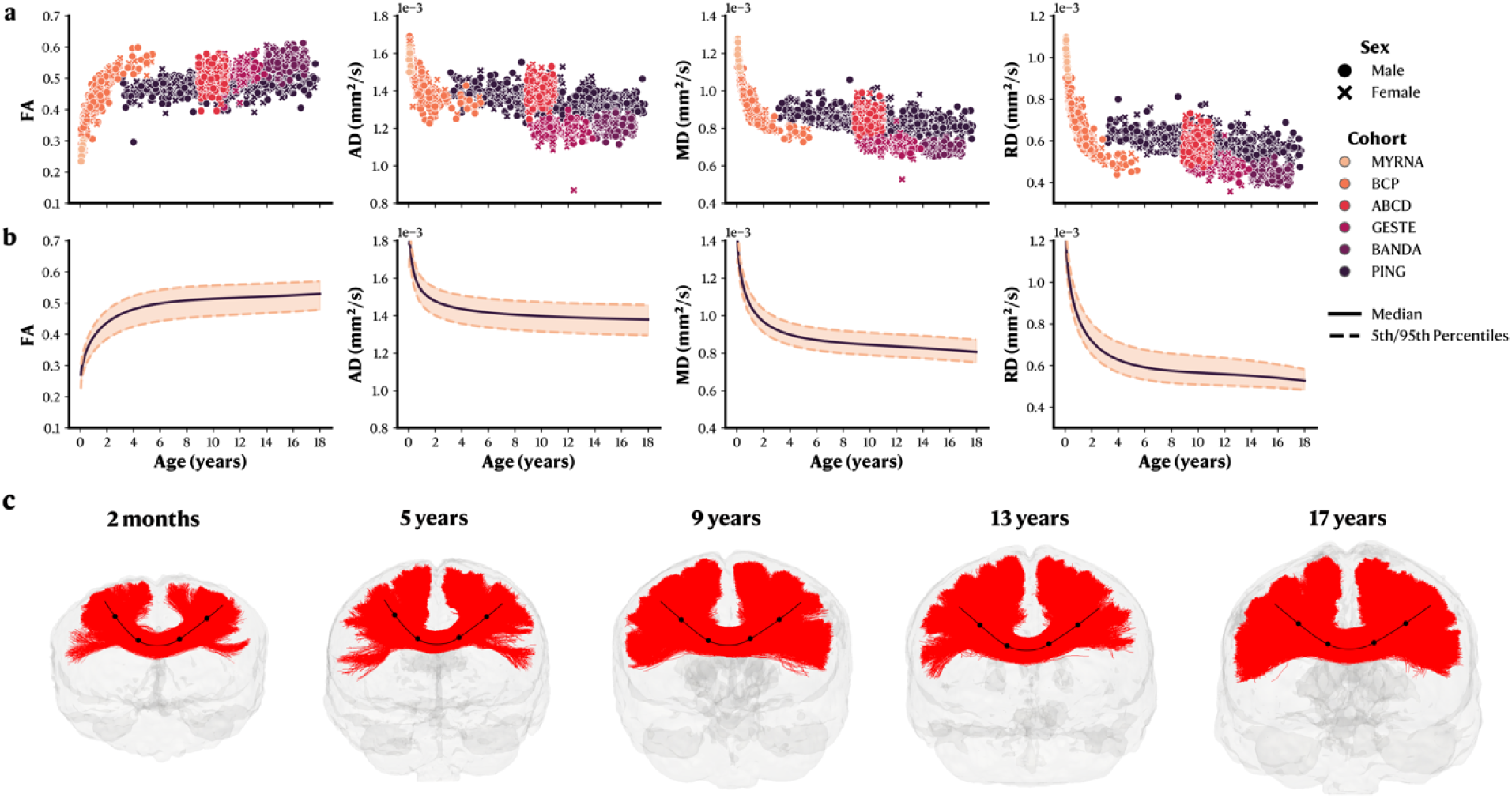
sf-pediatric can derive known WM microstructural changes across the developmental age range on the posterior frontal part of the corpus callosum. **a.** Raw data for each metric in function of age, stratified by sex and cohort. **b.** GAMLSS normative models with median and 5^th^/95^th^ percentiles. **c.** Visual representation of the mid-frontal part of the corpus callosum for five different subjects spanning the developmental age range. Centroid line showing the bundles’ sub-sections from which metrics can be derived. FA: Fractional Anisotropy. AD: Axial Diffusivity. RD: Radial Diffusivity. MD: Mean Diffusivity.

### sf-pediatric reproduces structural connectome network analysis across the developmental age range

In addition to extracting WM bundles, sf-pediatric can also derive structural connectome matrices, enabling evaluation of the structural network across the developmental age range. To showcase a possible application, we performed connectomics analyses using the Desikan-Killiany atlas^44^ for participants younger than 3 months and the brainnetome atlas for preadolescents^45^ (see Methods) across all included participants. Using COMMIT2-filtered matrices to remove false-positive streamlines (e.g., keeping streamlines with non-zero weights)^46,47^, we computed the connectome density, global efficiency, local efficiency, modularity, and average betweenness centrality for each participant. This enabled a good coverage of the three main types of network metrics: integration, segregation, and centrality. All metrics were then modeled as functions of age using GAMLSS models^29^ controlled for sex and cohort effects. Connectome density and global efficiency decreased until approximately 10 years of age, then increased until 18 years of age. Local efficiency and modularity remained stable across the developmental age range. Betweenness centrality decreased throughout the developmental age range (Figure 6a-b). Those results were consistent with prior findings^48^. To assess the stability of the extracted connections, we calculated a bootstrapped frequency percentage to quantify how frequently a single connection was found within a single age bin (see Methods). By constraining connections to be found at least 90% of the time, stable patterns were observed throughout the developmental age range, although variability increased in higher age bins (Figure 6c). These results align with our analysis and previous findings regarding connectome density^48^ and possibly reflect the pruning mechanism occurring during this age period^13^.

**Figure 6.**
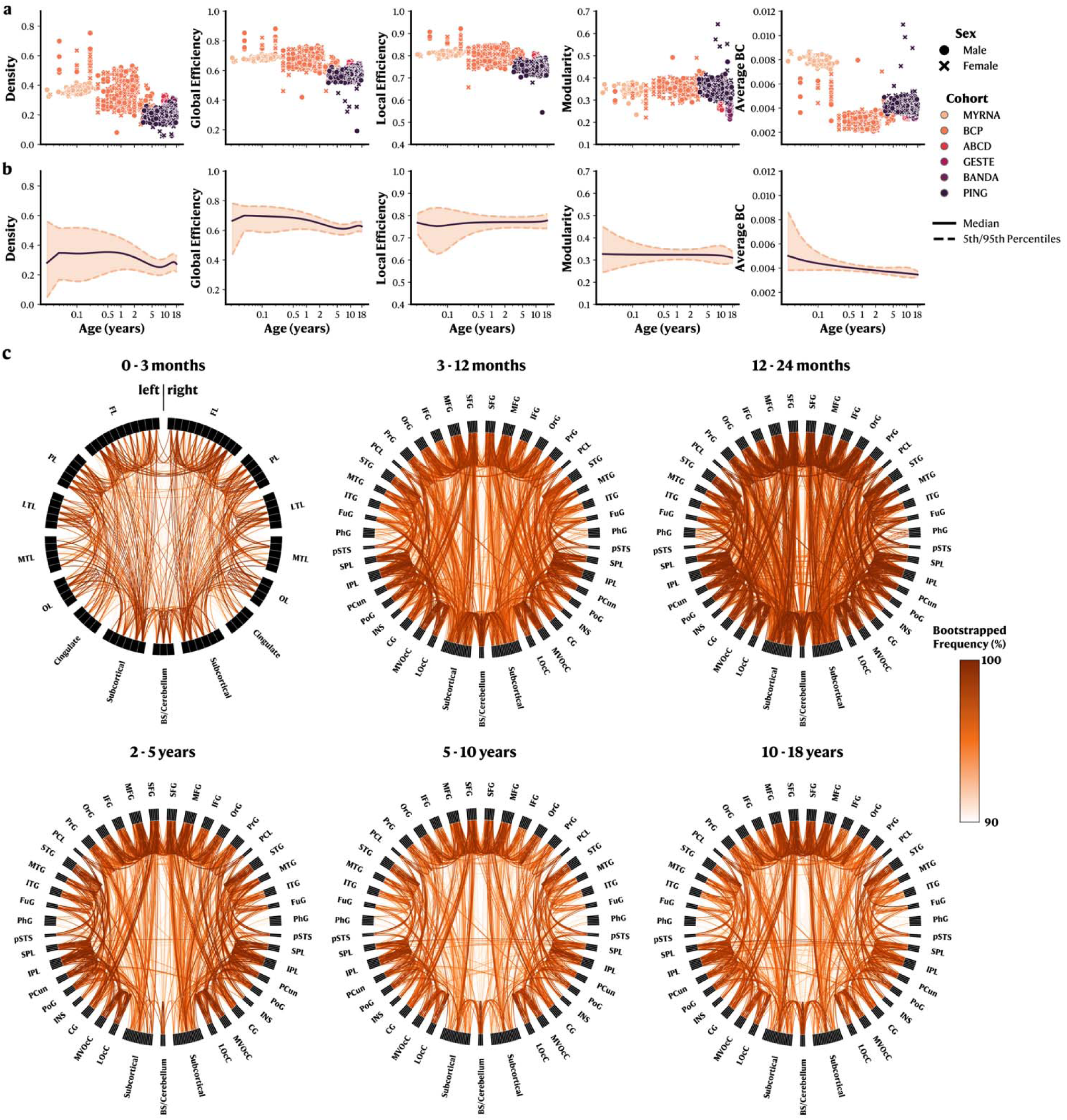
sf-pediatric enables the network analysis of the structural connectivity across the developmental age range, reproducing the known evolution of network properties during development. **a.** Raw data for each network metric as a function of age, stratified by sex and cohort. **b.** GAMLSS normative models with median and 5^th^/95^th^ percentiles. A log scale was used for age visualization. **c.** Chord charts of the most frequent connections (seen in over 90% over participants per age bin) for six age bins spanning the developmental age range. 0-3 months: Connectivity based on the Desikan-Killiany atlas. FL: Frontal lobe. PL: Parietal lobe. LTL: Lateral temporal lobe. MTL: Medial temporal lobe. OL: Occipital lobe. BS: Brainstem. 3 months – 18 years bins: Connectivity based on the brainnetome atlas for preadolescents. SFG: Superior frontal gyrus. MFG: Middle frontal gyrus. IFG: Inferior frontal gyrus. OrG: Orbital gyrus. PrG: Precentral gyrus. PCL: Paracentral gyrus. STG: Superior temporal gyrus. MTG: Middle temporal gyrus. ITG: Inferior temporal gyrus. FuG: Fusiform gyrus. PhG: Parahippocampal gyrus. pSTS: Posterior superior temporal sulcus. SPL: Superior parietal lobule. IPL: Inferior parietal lobule. PCun: Precuneus. PoG: Postcentral gyrus. INS: Insula. CG: Cingulate gyrus. MVOcC: Medioventral occipital cortex. LOcC: Lateral occipital cortex. BS: Brainstem.

## Discussion

Our proposed pipeline significantly differs from existing pediatric alternatives^18,20^. To our knowledge, this is the first “intelligent” pipeline that leverages information from normative models to adapt processing priors to individual subjects. By comparing results obtained with and without age-adaptable priors, we showed that tailoring priors to each subject yields better local modeling and cortical fanning, without affecting reproducibility or subsequent brain-behavior analyses, which have been previously identified as current challenges in pediatric dMRI^3,9^. Additionally, we propose an end-to-end solution for both the analysis of major WM bundles and the structural connectome. We showcased its end-to-end capabilities by deriving normative curves for diffusion metrics for each bundle, showing high consistency with previous studies^10,25,41–43^. Similarly, we leveraged graph theory concepts to build normative models of network organization across the developmental age range and showed consistency with existing findings^48^. Those results are strong evidence of the quality, reproducibility, and robustness of sf-pediatric across the developmental age range, as well as of the sub-optimal results that uniform priors can yield, making it suitable for a range of studies. Our reliance on the state-of-the-art Nextflow modules library nf-neuro^27^ ensures thoroughly tested, reproducible processing steps while facilitating maintenance and improvements, and making it easy to upgrade to newly developed tools. Crucially, we integrate a complete, interactive quality control procedure that leverages both population-wide quantitative data, which other alternatives do not provide, and single-subject reports for visual assessment of data quality.

A crucial advantage of sf-pediatric is that processing is performed directly in the native subject space. Tractography is performed directly from the subject’s diffusion signal, thereby avoiding any inherent biases and limitations arising from the use of atlases. While bundle extraction is performed using age-matched WM atlases, these atlases are not used to generate streamlines; instead, they are used to match existing subject-generated streamlines with the same geometry and location. This approach enables a more faithful assessment of each subject underlying WM organization. Structural connectomes also benefit from the same native space processing. However, concerns have been raised in the literature regarding false-positive connections in structural connectomes^49^. While sf-pediatric is not exempt from this limitation, we incorporated COMMIT2 filtering, combined with our age-adaptable diffusivity priors, to limit the impact of false-positive connections on our resulting connectomes^46,47^. This innovative approach enables the derivation of structural connectomes across the developmental age range, which, to our knowledge, is not supported by other pediatric-tailored pipelines. Similarly, sf-pediatric includes the option of fitting advanced diffusion models, such as NODDI^14^ and freewater-corrected DTI^50^, to derive additional WM metric maps. Those models rely on our age-adaptable priors for calibration, making sf-pediatric a unique opportunity to leverage these advanced WM metric maps in subsequent analyses.

The present work has some limitations. First of all, while we currently support connectomics analyses across the full pediatric age range, we only support two brain atlases: the Desikan-Killiany (DK)^44^ atlas for participants under 3 months and the brainnetome atlas for preadolescents^45^. However, this is on the roadmap for sf-pediatric, as we aim to support multiple brain atlases that would cover the whole age range. While we provide statistics-ready data in tabular format or connectivity matrices, we currently do not implement harmonization techniques required for neuroimaging studies, as they rely on specific assumptions that are difficult to incorporate into a neuroimaging pipeline^51^. Fortunately, sf-pediatric finds itself in a unique position due to the inclusion of normative models, which potentially offer a unique target for post-hoc harmonization, with Clinical-ComBAT^52^ already available in the nf-neuro library. Those normative models themselves have limitations, as they were derived primarily from participants in North America. While the curves showed expected trajectories compared to studies using participants with diverse backgrounds^11,25,41–43^, future curves designed for harmonization should represent participants from different cultural backgrounds.

Built on the nf-neuro library^27^ and following the nf-core guidelines^33^, sf-pediatric is integrated into an ecosystem surrounding the Nextflow^28^ workflow manager, recently proposed as the optimal solution for neuroimaging pipelines^53^, that enables easy, rigorously tested continuous development and integration. However, it also allows users to deploy pipelines across most available platforms, ranging from local computers to HPC servers and cloud services. While it may not be needed for small datasets, the increasingly larger size of developmental cohorts makes those remote computing resources of utmost importance. While complex, sf-pediatric supports the processing of large-scale datasets. Its complexity, stemming from its versatility, results in many configurable parameters. However, to ensure users have maximum control over how their data is processed, we expose every parameter as a configurable command-line argument. This ensures transparency and, combined with the quality control reports and the boilerplate methods section, successfully promotes sf-pediatric as a “glass-box” solution^53^ comparable to other neuroimaging pipelines and frameworks^17,54^. Additionally, to further improve the transparency of sf-pediatric, we provide documentation, and all code is openly available on GitHub (https://github.com/scilus/sf-pediatric.git), allowing the community to open issues for desired features and provide feedback. Due to the reliance on the nf-neuro library (https://github.com/nf-neuro/modules.git), which contains the building blocks for the processing steps underlying sf-pediatric, every improvement made in a block can be easily integrated into the pipeline, ensuring maintainability as new state-of-the-art tools are developed and making it easy to create “à-la-carte” pipelines.

Overall, we propose a new age-adaptable end-to-end processing pipeline for pediatric diffusion MRI that, for the first time, incorporates data from normative models directly into the processing stream. The quality and robustness of the processed results make sf-pediatric suitable for a wide range of developmental studies with different aims. In contrast, its scalability, portability, and quality control reports will facilitate its adoption across a wide range of users.

## Methods

### Participants’ data

Data used to validate and assess normative trajectories were obtained from 1,703 participants across six cohorts spanning the pediatric age range (Supplementary Table 1). Participants from the Maternal and Youth Research on Neurodevelopment and behAvior (MYRNA; 83 participants; aged 0-0.2 years old), Baby Connectome Project (BCP; 406 participants; 0.1-5.4 years old)^55^, Adolescent Brain Cognitive Development (ABCD; 300 participants; 8.9-11.0 years old)^21,56^, GESTation and Environment (GESTE; 199 participants, 9.1-13.8 years old)^57^, Boston Adolescent Neuroimaging of Depression and Anxiety (BANDA; 203 participants, 14-17 years old)^58^, and Pediatric Imaging, Neurocognition, and Genetics (PING; 512 participants, 3.2-17.9 years old)^59^ were downloaded, converted in BIDS datasets, and controlled for incomplete or bad quality acquisitions. For all cohorts, study protocols were approved by their respective Institutional Ethics Board. Across all cohorts, Siemens scanners were the most widely used, accounting for 59.95% of participants, while Philips and GE accounted for 23.49% and 15.80%, respectively; the remaining 0.76% of participants had missing scanner information. Acquisition schemes varied across cohorts, ranging from 30-direction acquisitions with a single low b-value to multi-shell, high-angular-resolution diffusion imaging (HARDI). This variability in acquisition protocols ensured the pipeline was tested across a broader range of protocols, thereby demonstrating its generality to other clinical or non-clinical datasets. Details of the acquisition protocols are provided in Supplementary Table 2. While an overview of the processing profiles is presented in Figure 4, the following sections will provide in-depth details on the different processing steps and the libraries used.

### Computing normative models of the brain diffusivities

To derive normative models of the brain diffusivities, we processed all participants using the *tracking* profile to generate diffusion tensor imaging (DTI) metric maps, specifically fractional anisotropy (FA), axial diffusivity (AD), radial diffusivity (RD), and mean diffusivity (MD) (the full dMRI and DTI processing pipeline is described below). To reliably extract single fiber population and cerebrospinal fluid (CSF) voxels, we designed a set of heuristics and thresholds, using both FA and MD maps. To limit the search space within each volume, we created a cube centered in the volume, extending 20 voxels in each direction. This restricted search space ensures the selection of voxels that represent either the corpus callosum/corticospinal tract or the lateral ventricles. Single fiber voxels were extracted by selecting voxels with an FA > 0.65 but < 0.95 (to avoid degenerate voxels) within the restricted search space. CSF voxels were extracted by selecting voxels with MD > 0.002 mm^2^/s but with an FA < 0.1. Voxels meeting those criteria were converted to binary masks and used to compute mean values for FA, AD, RD, and MD. This procedure was repeated for each participant from each cohort. A visual representation of an example single fiber and CSF masks is presented in Supplementary Figure 38.

For each metric, we fitted a Generalized Additive Model for Location, Scale, and Shape (GAMLSS)^29^, widely used in brain chart studies^25,26^. In its general form, a GAMLSS model assumes that each observation *Y_i_* follows a probability distribution *D*(*μ_i_,σ_i_, v_i_, τ_i_*) where the parameters *μ*_*i*_, σ_*i*_, *v*_*i*_, and *τ*_*i*_ are linked to additive predictors through link functions *g_k_* (·). Each link function may include fixed effects *β* with design matrix *X* (e.g., for covariates such as sex), random effects *γ* with design matrix *Z* (e.g., to model cohort-specific variability), and smooth functions *f* applied to continuous covariate *x*_*i*_ (here, the age variable). In compact form, the additive predictor for each parameter *θ_k_* can be expressed as:

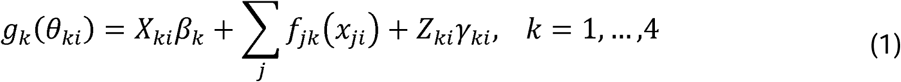

This framework allows each distributional parameter (*μ*_*i*_, *σ*_*i*_, *v*_*i*_, *τ*_*i*_) to vary as a function of covariates and random effects, while flexibly capturing nonlinear age-related changes in brain diffusivities across development. Aligning our approach with that of Bethlehem et al^26^ and Kim et al.^25^, we modeled the association between age and brain diffusivities using a Generalized Gamma distribution. Both the mean (*μ*) and scale (*σ*) parameters were allowed to vary as fractional polynomial (*f*) functions of age and sex, with random intercepts for cohort, while the shape (*v*) parameter was held constant using an identity matrix (*m*). The fitted model can thus be expressed as:

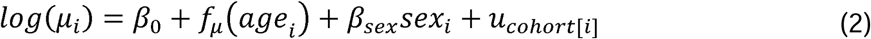

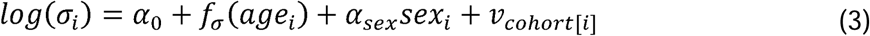

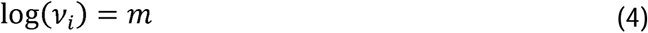

For each of the four fitted models, we ran polynomial-degree combinations from 1 to 3 and selected the best-fitting model using the Bayesian Information Criterion (BIC). To use the normative models as quierable curves in sf-pediatric for unseen data from unknown sites, we sampled each model to generate median data. Then, we approximated each of the four final fitted models using either logistic, Gompertz, Weibull, exponential, power, linear, quadratic, cubic, or log equations. The equation that best approximated each median data was selected based on the R^2^ value (Supplementary Figures 1-4), resulting in the following equations:

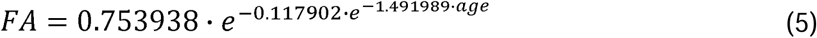

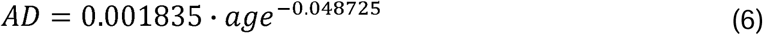

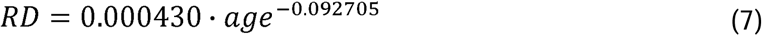

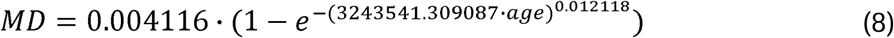

To quantify how different the derived priors are compared to the reference young-adults priors, we computed the average diffusivity of the fiber response function, defined as the average of the three eigenvectors forming the tensor as defined in equation (9). For experimental differentiation purposes, we named this metric the average diffusivity of the fiber response function, although the equation is identical to the MD metric.

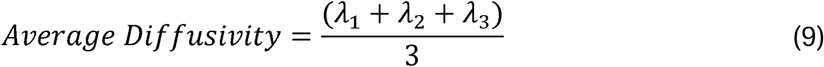

### Evaluating the impact of including age-adaptable priors

To assess the effects of the age-adaptable priors on processed data, we selected a single cohort, the BCP cohort^55^, as our study population because: 1) its age range spans the range where the age-adaptable priors differ most from the young adult priors, and 2) to limit the need for data harmonization across cohorts/sites. First, we processed the BCP cohort using the young-adult prior (e.g., a fiber response function of 15 x 10^-3^ mm^2^/s, 4 x 10^-3^ mm^2^/s, 4 x 10^-3^ mm^2^/s) and performed bundle extraction and tractometry. Then, using the exact same parameters, we reprocessed the BCP cohort with the age-adaptable priors derived using equations (6) and (7). To investigate the impact on cortical fanning, we selected the corticospinal track (CST) as our bundle of interest, as it is known to be present early in neurodevelopment^3^. To measure cortical fanning, we computed the surface area of the bundle at the cortical endpoint. Surface voxels were defined as a voxel with at least one zero-valued neighboring voxel. Then, the surface area was calculated as the voxel resolution squared (in millimeters) multiplied by the number of voxels that meet the surface requirements. This metric is directly available in the *bundling* profile (see the *Bundling processing profile* section below). To compare between young-adult priors and age-adaptable priors, we used a paired sample t-test with a one-sided p-value.

Additionally, we tested whether including age-adaptable priors would affect the bundle’s reproducibility in unseen test-retest data from the open-access MASiVar dataset (n=83 participants aged 5 to 8 years, OpenNeuro accession number: ds003416)^60^. We processed the MASiVar dataset twice using the *tracking* and *bundling* profiles, once with the reference young-adult priors and once with the age-adaptable priors. Using a built-in option in sf-pediatric, we registered each bundle in the MNIPediatricAsym template space^61^. Using the scilpy toolbox^36^, we computed the dice score (weighted by streamline density) between each test-retest acquisition on both the left and right CST extracted using either the young-adult priors or the age-adaptable priors. We then compared the weighted dice scores from the young-adult and age-adaptable priors using a paired sample t-test with a two-sided p-value.

To investigate whether the introduction of age-adaptable priors would affect the ability to derive meaningful relationships with external variables, we pulled the raw expressive and receptive language scores from the Mullen Scales of Early Learning (MSEL) in the BCP cohort^55,62^. This test is specifically tailored to measure language abilities in children up to 68 months old. We selected the arcuate fasciculus (AF) as our bundle of interest due to its known role in language abilities^32^. Using the fixel-based AFD to capture the variability introduced by the age-adaptable priors, we fitted multivariate regression models with the raw language scores as the dependent variable and fixel-based AFD as the independent variable, using data derived from either the young-adult or age-adaptable priors. A single model was fitted for both scores, both left and right AF, and for the young adult and age-adaptable priors, resulting in eight models. For all models, we included sex and age as control variables. Reported statistics included the R^2^ for the model, the beta coefficient for the fixel-based AFD, and the p-value for the fixel-based AFD. To ensure the age-adaptable priors did not introduce spurious associations, we repeated the analysis described above using two behavioral scores from the Preschool-Aged Child Behavioral Checklist (CBCL) in the BCP cohort: internalization and externalization. Similarly to the language scores, we fitted a single model for both scores, both left and right superior longitudinal fasciculus (SLF), and for the young adult and age-adaptable priors, resulting in eight models. We selected the SLF as our bundle of interest since there is no clear evidence of implication in either externalization or internalization behaviors and should, therefore, return non-significant results. All models were controlled for sex and age.

### Building normative models of WM bundles and structural networks

To showcase the results produced by sf-pediatric throughout the developmental age range, we processed all participants using the *tracking* and *bundling* profiles (see sections below). Using the mean values for AD, RD, MD, and FA directly exported by the pipeline, we built normative models for 31 bundles using GAMLSS models as defined in equations (2), (3), and (4). We fitted models with fractional polynomial degrees ranging from 1 to 3 and selected the best model using BIC. Additionally, we further processed all participants using the *segmentation* and *connectomics* profiles (see sections below). Participants aged under 3 months were segmented using the Desikan-Killiany atlas^44^, while older participants were segmented using the brainnetome atlas for preadolescents^45^. Using the structural connectome matrices, we computed connectome density, global efficiency, local efficiency, modularity, and average betweenness centrality using the NetworkX Python package^63^. For each metric, we fitted a GAMLSS model using equations (2), (3), and (4) to derive normative curves of network characteristics throughout the developmental age range. Similar to the bundle models, we fitted a range of models with different fractional polynomial degrees and selected the best model using BIC. To quantify the stability of the extracted connections throughout development, we computed the extraction frequency for each connection in six age bins: 0 – 3 months, 3 – 12 months, 12 – 24 months, 2 – 5 years, 5 – 10 years, and 10 – 18 years. Since the number of participants varied across age bins, we used a bootstrapping approach based on the age bin with the fewest participants. For each age bin, we computed the frequency of each connection using a random subsample of participants, matching the number of participants in the smaller age bin. This frequency was then converted into percentages and averaged across 1,000 iterations. This yielded a bootstrapped frequency value for each of the six age bins.

### Tracking processing profile

#### Preprocessing of diffusion and anatomical volumes

To perform tractography in subject-space, sf-pediatric proposes a “tracking” profile. This is the main entry point of the pipeline, as it includes all preprocessing steps applied to the raw diffusion and anatomical data. When selected, raw diffusion volumes will be denoised using the MP-PCA algorithm^64^ implemented in the Mrtrix software^38^, corrected for Gibbs ringing artefacts using Mrtrix^65^, corrected for susceptibility distortions using *TOPUP* followed by eddy from the FSL library^37^, brain extracted using *synthstrip*^66^ (using alternative infant model weights for participants < 2 years) from the FreeSurfer package^34^, bias field corrected using the N4 algorithm^67^ from the ANTs library^35^, intensity-normalized using the *dwinormalise* command from Mrtrix^38^, and resampled to a resolution of 1 mm isotropic (configurable). Concurrently, anatomical images will undergo similar processing steps, including denoising (using the *nlmeans* algorithm from DIPY^68^), bias-field correction, resampling to an isotropic resolution of 1 mm (configurable), and brain extraction. Since it is a common practice to acquire a T2w image before a T1w image in infants due to the flipped contrast, the pipeline will look for both in the BIDS structure. If both are found, the pipeline will compute an affine transformation using the ANTs library^35^ to co-register the two modalities.

#### Diffusion tensor imaging and fiber-oriented distribution functions

Following preprocessing, diffusion tensor imaging (DTI)^5^, a widely used diffusion model, will be used to estimate water diffusion in each voxel using the DIPY^68^ library. If not specified at runtime, shells below 1200 s/mm^2^ will be used to fit the DTI model. Fractional anisotropy (FA), mean diffusivity (MD), radial diffusivity (RD), and axial diffusivity (AD) maps, amongst others, will be computed from the estimated tensors. Those white-matter microstructure metric maps will be kept and reused to derive per-bundle and per-connection metrics in the *bundling* and *connectomics* profiles, respectively.

Leveraging diffusivities from the proposed normative models across the developmental age range, fiber orientation distribution functions (fODF) will be estimated using constrained spherical deconvolution from the DIPY toolbox^68^ using a single-shell, single-tissue method. Unless specified, shells higher than 700 s/mm^2^ will be used. To set the fiber response function (FRF), the underlying tensor’s eigenvalues *λ*_1_, *λ*_2_, and *λ*_3_ will be modeled using equations (6) and (7), where equation (7) will be used for both *λ*_2_ and *λ*_3_, resulting in an age-adaptable FRF. To our knowledge, this is the first pipeline to leverage normative data “on-the-fly” to adapt processing priors based on subject age. Using the scilpy toolbox^36^, fODF-derived metrics, such as apparent fiber density (AFD) and the number of fiber orientations per voxel, will be computed and stored as metric maps.

#### Registration and tissue segmentation

To ensure optimal multimodal registration between the T1w/T2w anatomical space and the diffusion space, sf-pediatric selects the best pair of moving and target images based on the participant’s age and performs registration using the ANTs library^35^. For subjects under 6 months, the T2w image will be registered using the b=0 and MD maps as targets, whereas for older participants or subjects with only a T1w image, the b=0 and FA maps will be used as targets. This adaptability ensures that the varying contrast in the first years of life is taken into account^3,9,11^, which, otherwise, can lead to poor tissue alignment.

Following registration in diffusion space, the anatomical image will be segmented to obtain tissue probability maps. Since this can be challenging in neonates, sf-pediatric leverages a 4D brain atlas derived from the Baby Connectome Project that includes tissue probability maps^69^. Using a neonate, 3 months, 6 months, 12 months, and 24 months templates, each participant under 2.5 years old will be matched to their closest template. The selected template will be registered in the subject space using non-linear transformations from the ANTs library^35^. Computed transformations will then be used to warp tissue probability maps into subject space. Probability maps can then be thresholded to obtain binary masks when required. For older participants, tissue segmentation will be performed using the FAST algorithm from the FSL library^37,70^. Similarly, resulting tissue probability maps will be thresholded to obtain binary masks when required.

#### Tractography

To perform streamline reconstruction, sf-pediatric proposes, by default, two tractography methods: standard tracking using a binary stopping criterion and particle filter tracking (PFT)^71^. The standard tracking algorithm uses a mask to determine the end of a streamline (e.g., by stopping when it is outside the tracking mask). The PFT method leverages anatomical information to assess the end of a streamline using a continuous mapping criterion (CMC)^71^. The CMC leverages partial-volume tissue estimation maps to determine when a streamline should stop (similar to the anatomically constrained tractography method^72^). When a streamline incorrectly stops (e.g., in a voxel of either WM or CSF), the PFT method will try to find an alternate segment to reach the correct stop signal (e.g., the gray matter (GM)). This method is particularly useful in connectomics analyses, as it ensures all streamlines start and end in GM tissue. Both methods support customizable tracking parameters via the command line (step size, maximum/minimum length, maximum angle, etc.) and can be toggled on/off, enabling users to fine-tune their execution. For standard tracking, users can select either a white matter mask or a thresholded FA map as the seeding mask, while PFT tracking also supports seeding from the WM/GM interface. For both methods, deterministic and probabilistic algorithms are available, with the latter as the default. If both tracking methods are used, the pipeline will combine the resulting tractograms into a single tractogram, which will be reused in downstream processing.

#### NODDI and freewater-corrected DTI

If selected by the user, NODDI and freewater-corrected DTI models can be fitted on the processed DWI volume using the AMICO implementation^14,50,73^. Similarly to the fODF fitting, the diffusivity priors (parallel, perpendicular, and isotropic) will be derived using equations (6), (7), and (8) based on the subject’s age. Metrics such as the isotropic volume fraction, intra-cellular volume fraction, extra-cellular volume fraction, and orientation dispersion index will be computed from the NODDI model and stored. For the freewater elimination model, all DTI metrics will be recomputed using the freewater-corrected diffusion volume to yield freewater-corrected DTI metric maps. Those maps will be stored as metric maps combined with the fiber volume and freewater maps from the freewater model.

### Bundling processing profile

#### Bundle extraction

Following the tracking processing profile, users can select two different analysis trajectories. One of which consists of extracting known WM bundles and computing diffusion-related metrics along each bundle. When this profile is selected, the BundleSeg algorithm^74^ combined with age-matched WM atlases will be used to extract 51 WM bundles from the final processed tractogram (Supplementary Table 4). WM atlases are selected by taking the one closest to the participant’s age. Each WM atlas was built by incrementally registering the adult version^75^ to younger anatomical templates. An affine transformation is then computed to map the atlas template into subject space. Then, the BundleSeg algorithm will extract the WM bundles based on the streamline’s geometry and distance from the warped reference bundles. Extracted bundles are then filtered for spurious streamlines to remove possible false positives.

#### Tractometry

Using the scilpy toolbox^36^, the fixel-based AFD is computed for each bundle^76^. Then, each bundle is segmented based on a user-defined number of points, generating sub-sections for each bundle. Using all metric maps computed during previous processing steps, mean values for each sub-section and for the complete bundle are calculated. Those metrics include diffusion metrics (e.g., FA, MD, RD, AD, AFD, NuFO, fixel-based AFD, etc.), as well as macroscale metrics such as bundle volume, number of streamlines, curvature, and surface area of bundle endpoints. To facilitate the inclusion of these metrics in subsequent statistical analyses, all bundle metrics (per section or for the complete bundle) are concatenated across all subjects into a clean tab-separated values file in long format.

### Segmentation processing profile

In addition to extracting WM bundles, users can also select a trajectory for connectome analysis. However, cortical and subcortical segmentations are required to derive a structural connectome. To facilitate this step, we include a segmentation profile that uses either the M-CRIB-S tool for neonatal segmentation (for participants under 3 months)^77^, the FastSurfer deep-learning tool^78^, or the FreeSurfer library (recon-all^34^ or recon-all-clinical^79^). For participants older than 3 months, the brainnetome atlas for preadolescents is then mapped in subject space using surface-based registration methods^45^. Opting for registration based on surfaces rather than volumes yields better alignment with individual cortical folding patterns than traditional methods^80^. For younger participants, the original M-CRIB-S(DK) atlas from the M-CRIB-S tool is used directly for connectome analysis^44,77^. Following segmentation, volume, surface area, and cortical thickness are extracted for each parcel, concatenated across all subjects, and outputted in a clean tab-separated values file in wide format.

### Connectomics processing profile

Structural connectivity matrices are generated using the scilpy toolbox^36^. Briefly, labels in anatomical space are transformed into diffusion space using already computed transformations. The final processed tractogram is then decomposed to extract all streamlines connecting each pair of parcels. To limit the number of false-positive connections, the Convex Optimization Modeling for Microstructure-Informed Tractography (COMMIT2) filtering method is applied to the decomposed tractogram, and only connections with non-zero weights are retained^46,47^. Priors for COMMIT executions are fetched “on-the-fly” based on the participant’s age using equations (6), (7), and (8). Fixel-based AFD is then computed for all connections^76^, and structural connectivity matrices are generated by calculating the number of streamlines, mean streamline length, FA, MD, AD, RD, AFD, fixel-based AFD, and NuFO, amongst other metrics.

### Automatic Quality Control

Each pipeline execution yields two types of QC reports rendered in an HTML format using MultiQC^40^: one assessing the distribution of quantitative QC indicators across the population (population-wide report) and the other showcasing a visual assessment of key processing steps for a single subject (single subject report). To facilitate QC, we provide an initial automatic labeling using the “pass”, “warn”, or “fail” system in the population-wide report. For the framewise displacement, we automatically flag subjects with a maximum framewise displacement value between 0.8mm and 2mm as “warn”, whereas participants exceeding 2mm are labeled as “fail”. To evaluate the quality of the tractogram WM coverage, we compute the tract density image from the tractogram and evaluate the overlap with the WM mask using the DICE score. Participants with a DICE score <0.8 are labeled as “fail”, whereas participants between 0.8 and 0.9 are labeled as “warn”. Additionally, we compute the median and inter-quartile range (IQR) of the number of streamlines within the tractogram across the population. Then, we flag subjects that exceed 1.5*IQR as outliers and report them as “fail”. To provide a quality metric for the bundle extraction process, we report the percentage of recognized bundles. Participants with <80% recognized bundles are labeled as “fail”, while participants with 80% to 90% recognized bundles are labeled as “warn”. Finally, for both cortical and subcortical volumes, we use the IQR detection method to count parcels considered as outliers per subject. Then, using the total number of regions, we compute the percentage of parcels considered outliers (defined as outside the 1.5*IQR). Participants with 10-20% of parcels considered as outliers are labeled as “warn”, whereas participants with more than 20% are labeled as “fail”. Combined, those QC steps ensure easy detection of potential processing issues that can then be further investigated using the subject-specific report.

When a participant gets flagged for further investigation, users can open the subject-specific report. In this report, we present visual assessments of key processing steps, including: visualization of the b-values direction on the sphere, pre-post TOPUP GIF, line plot of movement between acquired directions, pre-post eddy current correction GIF, screenshot of FA, MD, RGB FA, and NuFO metric maps, anatomical to diffusion space registration GIF, tissue segmentation screenshot, tracking coverage, and cortical/subcortical segmentation screenshot. Each section contains a small description of the presented figure and what to look for in an efficient QC. This enables inexperienced users to assess the quality of their processed data autonomously. The final decision to exclude a participant from downstream analyses rests solely with the user.

## Acknowledgements

Data used in the preparation of this article were obtained from the Adolescent Brain Cognitive Development^SM^ (ABCD) Study (https://abcdstudy.org), held in the NIMH Data Archive (NDA). This is a multisite, longitudinal study designed to recruit more than 10,000 children age 9-10 and follow them over 10 years into early adulthood. The ABCD Study® is supported by the National Institutes of Health and additional federal partners under award numbers U01DA041048, U01DA050989, U01DA051016, U01DA041022, U01DA051018, U01DA051037, U01DA050987, U01DA041174, U01DA041106, U01DA041117, U01DA041028, U01DA041134, U01DA050988, U01DA051039, U01DA041156, U01DA041025, U01DA041120, U01DA051038, U01DA041148, U01DA041093, U01DA041089, U24DA041123, U24DA041147. A full list of supporters is available at https://abcdstudy.org/federal-partners.html. A listing of participating sites and a complete listing of the study investigators can be found at https://abcdstudy.org/consortium_members/. ABCD consortium investigators designed and implemented the study and/or provided data but did not necessarily participate in the analysis or writing of this report. This manuscript reflects the views of the authors and may not reflect the opinions or views of the NIH or ABCD consortium investigators. The ABCD data repository grows and changes over time. The ABCD data used in this report came from <u>10.15154/z563-zd24</u>. DOIs can be found at https://nda.nih.gov/abcd/abcd-annual-releases. Data were provided by the Boston Adolescent Neuroimaging of Anxiety and Depression (BANDA) Consortium’s Human Connectome Project, supported by 1U01MH108168 (PIs: Susan Whitfield-Gabrieli, John Gabrieli). Data were also provided by the GESTE study, supported by the National Institute of Environmental Health (grant # R01ES027845), acquired by Dr Andrea Baccarelli and Dr Larissa Takser. Data were also provided by the UNC/UMN Baby Connectome Project, supported by the National Institute of Mental Health (NIMH) awards R01MH104324 and U01MH110274 (PIs: Jed T. Elison, Weili Lin) and accessed through the NIH-sponsored NIMH Data Archive (NDA) platform. Data were provided in part by the Pediatric Imaging, Neurocognition, and Genetics (PING) project, supported by awards RC2DA029475 and R01HD061414 from the National Institute on Drug Abuse and the Eunice Kennedy Shriver National Institute of Child Health and Human Development (PI: Terry Jernigan), and accessed through the NDA platform. Data were also provided by the Maternal Youth Research on Neurodevelopment and behAvior (MYRNA) supported by NIH grant R01MH119510 (PIs: Ardesheer Talati, Jonathan Posner, Larissa Takser). The authors thank all research staff and participants from all studies for their contributions and dedication to research. AG was supported by a Canadian Institute of Health Research Doctoral Award (#493956). MAB is supported by a Junior 1 career award from the Fonds de Recherche du Québec – Santé (FRQS).

## Data availability

Data from the ABCD cohort are available after obtaining a Data Use Certificate (DUC) on the NIH Brain Development Cohort platform (https://nbdc-datashare.lassoinformatics.com/). Data from the BANDA, PING, and BCP cohorts can be accessed through the NIMH Data Archive platform (https://nda.nih.gov/) after obtaining a DUC. For both platforms, researchers can apply for a DUC online, and study administrators will grant data access. Data from the MYRNA and GESTE cohorts can be obtained by contacting the corresponding principal investigators. Data from the MASiVar dataset is freely available on OpenNeuro (Accession Number: ds003416, https://openneuro.org/datasets/ds003416/versions/2.0.2). Code to reproduce analyses can be found here: https://github.com/gagnonanthony/sf-pediatric-paper.git.

