## Supplementary for "sf-pediatric: A robust and age-adaptable end-to-end pipeline for pediatric diffusion MRI": Supplement.docx

**Gagnon A, *et al.***

**Table of contents**

**Supplementary Tables**

**Supplementary Figures**

[Supplementary Figure 6. Age-adaptable priors do not introduce significant brain-behavior relationships. a. Visual representation of the left and right SLF in a glass brain. b-c. Multivariate regression results between fixel-based average fiber density (AFD Fixel) on the superior longitudinal fasciculus (SLF) and raw externalization and internalization scores in the BCP cohort before and after the inclusion of the custom age-adaptable priors. b. Scatter plots representing the association between raw externalization and raw internalization scores and AFD Fixel on both left and right SLF using the young-adult priors. c. Scatter plots representing the association between raw externalization and internalization scores and AFD Fixel on both left and right SLF using the age-adaptable priors. ß: Beta coefficients for the AFD Fixel variable. p: p-value for the AFD fixel variable. CBCL: Child Behavioral Checklist.	15](#_Toc219724977)

| **Cohorts** | **Sex** | **N** | **Age (median (range)) in years** |
| --- | --- | --- | --- |
| MYRNA | M | 35 | 0.08 (0.0-0.2) |
|  | F | 48 | 0.09 (0.0-0.3) |
| BCP | M | 191 | 1.00 (0.1-5.3) |
|  | F | 215 | 1.08 (0.1-5.4) |
| ABCD | M | 146 | 9.88 (8.9-11.0) |
|  | F | 154 | 9.83 (8.9-11.0) |
| GESTE | M | 120 | 11.81 (9.1-13.3) |
|  | F | 79 | 11.64 (9.5-13.8) |
| BANDA | M | 72 | 15.00 (14.0-17.0) |
|  | F | 131 | 15.58 (14.0-17.0) |
| PING | M | 268 | 11.08 (3.4-17.9) |
|  | F | 244 | 10.04 (3.2-17.8) |
| Total |  | 1703 | 9.75 (0.0-17.9) |

**Supplementary Table 1.** Participant information from all included cohorts. M: Male, F: Female. MYRNA: Maternal and Youth Research on Neurodevelopment and behAvior. BCP: Baby Connectome Project. ABCD: Adolescent Brain Cognitive Development. GESTE: GESTation and Environment. BANDA: Boston Adolescent Neuroimaging of Depression and Anxiety. PING: Pediatric Imaging, Neurocognition, and Genetics.

| **Cohort** | **Scanner** | **Modality** | **Slices** | **FOV** | **Resolution (mm)** | **TR (ms)** | **TE (ms)** | **Diffusion Directions** | **b-values** |
| --- | --- | --- | --- | --- | --- | --- | --- | --- | --- |
| MYRNA | Phillips | T2 | 180 | 240 x 240 | 1.0 x 1.0 x 1.0 | 2500 | 252 | - | - |
|  |  | Diffusion | 66 | 224 x 224 | 2.0 x 2.0 x 2.0 | 5650 | 96 | 64 | 1500 |
| BCP | Siemens | T1 | 208 | 256 x 256 | 0.8 x 0.8 x 0.8 | 2400/1060 | 2.24 | - | - |
|  |  | T2 | 208 | 256 x 256 | 0.8 x 0.8 x 0.8 | 3200 | 564 | - | - |
|  |  | Diffusion | 95 | 210 x 210 | 1.5 x 1.5 x 1.5 | 2640 | 88.6 | 144 | 500, 1000, 1500, 2000, 2500, 3000 |
| ABCD | Siemens | T1 | 176 | 256 x 256 | 1.0 x 1.0 x 1.0 | 2500 | 2.88 | - | - |
|  |  | Diffusion | 81 | 240 x 240 | 1.7 x 1.7 x 1.7 | 4100 | 88 | 96 | 500, 1000, 2000, 3000 |
|  | GE | T1 | 208 | 256 x 256 | 1.0 x 1.0 x 1.0 | 2500 | 2 | - | - |
|  |  | Diffusion | 81 | 240 x 240 | 1.7 x 1.7 x 1.7 | 4100 | 81.9 | 96 | 500, 1000, 2000, 3000 |
| GESTE | Phillips | T1 | 180 | 240 x 240 | 1.0 x 1.0 x 1.0 | 8.1 | 3.7 | - | - |
|  |  | Diffusion | 80 | 230 x 230 | 1.8 x 1.8 x 1.8 | 8200 | 97.5 | 64 | 1500 |
| BANDA | Siemens | T1 | 208 | 256 x 256 | 0.8 x 0.8 x 0.8 | 2400 | 2.18 | - | - |
|  |  | Diffusion | 92 | 210 x 210 | 1.5 x 1.5 x 1.5 | 3230 | 89.2 | 183 | 1500, 3000 |
| PING | GE | T1 | 170 | 256 x 192 | 0.94 x 1.2 x 1.2 | 8.1 | 3.5 | - | - |
|  |  | Diffusion | 53 | 240 x 240 | 1.85 x 1.85 x 2.5 | 8000 | 81 | 30 | 1000 |
|  | Siemens | T1 | 160 | 256 x 256 | 1 x 1 x 1.2 | 2170 | 4.33 | - | - |
|  |  | Diffusion | 68 | 240 x 240 | 2.5 x 2.5 x 2.5 | 9000 | 91 | 30 | 1000 |
|  | Phillips | T1 | 170 | 240 x 240 | 1 x 1 x 1.2 | 6.8 | 3.1 | - | - |
|  |  | Diffusion | 60 | 240 x 240 | 2.5 x 2.5 x 2.5 | 9000 | 91 | 32 | 1000 |

**Supplementary Table 2.** Details of acquisition protocols across all cohorts. MYRNA: Maternal and Youth Research on Neurodevelopment and behAvior. BCP: Baby Connectome Project. ABCD: Adolescent Brain Cognitive Development. GESTE: GESTation and Environment. BANDA: Boston Adolescent Neuroimaging of Depression and Anxiety. PING: Pediatric Imaging, Neurocognition, and Genetics.

|  | **Left AF** | | | **Right AF** | | |
| --- | --- | --- | --- | --- | --- | --- |
| Variables | ß | Std. Err. | *p*-value | ß | Std. Err. | *p*-value |
| **Young-Adult Priors** | | | | | | |
| **Raw expressive language** | | | | | | |
| **R^2^ = 0.88, *p* < 0.001** | | | | **R^2^ = 0.87, *p* < 0.001** | | |
| Intercept | -0.863 | 1.318 | 0.513 | 0.702 | 1.271 | 0.581 |
| AFD Fixel | 13.259 | 3.681 | < 0.001 | 8.283 | 3.782 | 0.030 |
| Age | 0.749 | 0.028 | < 0.001 | 0.773 | 0.028 | < 0.001 |
| Sex | 0.249 | 0.435 | 0.569 | 0.187 | 0.440 | 0.672 |
| **Raw receptive language** | | | | | | |
| **R^2^ = 0.86, *p* < 0.001** | | | | **R^2^ = 0.86, *p* < 0.001** | | |
| Intercept | -1.842 | 1.439 | 0.202 | -0.940 | 1.369 | 0.493 |
| AFD Fixel | 14.735 | 4.020 | < 0.001 | 12.606 | 4.073 | 0.002 |
| Age | 0.762 | 0.030 | < 0.001 | 0.773 | 0.030 | < 0.001 |
| Sex | 1.234 | 0.475 | 0.010 | 1.161 | 0.474 | 0.015 |
| **Custom Priors** | | | | | | |
| **Raw expressive language** | | | | | | |
| **R^2^ = 0.88, *p* < 0.001** | | | | **R^2^ = 0.87, *p* < 0.001** | | |
| Intercept | -2.403 | 1.491 | 0.109 | -0.344 | 1.449 | 0.812 |
| AFD Fixel | 15.968 | 3.986 | < 0.001 | 10.333 | 4.100 | 0.013 |
| Age | 0.761 | 0.026 | < 0.001 | 0.782 | 0.026 | < 0.001 |
| Sex | 0.473 | 0.433 | 0.276 | 0.293 | 0.447 | 0.512 |
| **Raw receptive language** | | | | | | |
| **R^2^ = 0.86, *p* < 0.001** | | | | **R^2^ = 0.86, *p* < 0.001** | | |
| Intercept | -1.794 | 1.633 | 0.273 | -1.076 | 1.554 | 0.489 |
| AFD Fixel | 12.675 | 4.368 | 0.004 | 11.483 | 4.398 | 0.010 |
| Age | 0.790 | 0.028 | < 0.001 | 0.796 | 0.028 | < 0.001 |
| Sex | 1.358 | 0.474 | 0.005 | 1.217 | 0.479 | 0.012 |

**Supplementary Table 3.** Multivariate regression results between fixel-based average fiber density (AFD Fixel) and raw language scores from the Mullen Scales of Early Learning (MSEL) before and after custom age-adaptable priors in the BCP cohort. AF: Arcuate Fasciculus.

|  | **Left SLF** | | | **Right SLF** | | |
| --- | --- | --- | --- | --- | --- | --- |
| Variables | ß | Std. Err. | *p*-value | ß | Std. Err. | *p*-value |
| **Young-Adult Priors** | | | | | | |
| **Raw CBCL internalization** | | | | | | |
| **R^2^ = 0.03, *p* = 0.062** | | | | **R^2^ = 0.03, *p* = 0.065** | | |
| Intercept | -0.799 | 2.174 | 0.714 | -0.608 | 2.220 | 0.785 |
| AFD Fixel | 2.332 | 4.672 | 0.618 | 1.722 | 4.468 | 0.701 |
| Age | 0.087 | 0.034 | 0.012 | 0.088 | 0.034 | 0.011 |
| Sex | 0.159 | 0.460 | 0.730 | 0.150 | 0.467 | 0.748 |
| **Raw CBCL externalization** | | | | | | |
| **R^2^ = 0.02, *p =* 0.112** | | | | **R^2^ = 0.05, *p* = 0.017** | | |
| Intercept | 14.191 | 4.768 | 0.003 | 18.448 | 4.790 | < 0.001 |
| AFD Fixel | -16.936 | 10.248 | 0.101 | -25.701 | 9.641 | 0.009 |
| Age | 0.087 | 0.074 | 0.246 | 0.096 | 0.073 | 0.191 |
| Sex | -2.006 | 1.010 | 0.049 | -2.417 | 1.007 | 0.018 |
| **Custom Priors** | | | | | | |
| **Raw CBCL internalization** | | | | | | |
| **R^2^ = 0.04, *p* = 0.050** | | | | **R^2^ = 0.03, *p* = 0.059** | | |
| Intercept | -0.946 | 2.039 | 0.644 | -0.182 | 2.001 | 0.928 |
| AFD Fixel | 2.572 | 4.135 | 0.535 | 0.705 | 3.977 | 0.860 |
| Age | 0.089 | 0.034 | 0.009 | 0.091 | 0.034 | 0.008 |
| Sex | 0.141 | 0.461 | 0.760 | 0.078 | 0.460 | 0.866 |
| **Raw CBCL externalization** | | | | | | |
| **R^2^ = 0.02, *p* = 0.168** | | | | **R^2^ = 0.04, *p* = 0.033** | | |
| Intercept | 11.820 | 4.483 | 0.009 | 15.556 | 4.332 | < 0.001 |
| AFD Fixel | -10.606 | 9.091 | 0.245 | -19.507 | 8.614 | 0.025 |
| Age | 0.079 | 0.074 | 0.288 | 0.098 | 0.073 | 0.186 |
| Sex | -1.944 | 1.013 | 0.057 | -2.254 | 0.996 | 0.025 |

**Supplementary Table 4.** Multivariate regression results between fixel-based average fiber density (AFD Fixel) and raw internalization and externalization scores from the Child Behavioral Checklist (CBCL) before and after custom age-adaptable priors in the BCP cohort. SLF: Superior longitudinal fasciculus.

| **Abbreviation** | **Full name** |
| --- | --- |
| AC | Anterior commisure |
| AF_(L/R) | Arcuate Fasciculus |
| CC_Fr_1 | Corpus callosum, Frontal lobe (anterior part) |
| CC_Fr_2 | Corpus callosum, Frontal lobe (posterior part) |
| CC_Oc | Corpus callosum, Occipital lobe |
| CC_Pa | Corpus callosum, Parietal lobe |
| CC_Pr_Po | Corpus callosum, Pre/Post central gyri |
| CC_Te | Corpus callosum, Temporal lobe |
| CG_(L/R)_An | Cingulum, anterior part |
| CG_(L/R)_curve | Cingulum, curved part |
| CG_(L/R)_Po | Cingulum, posterior part |
| CG_(L/R) | Cingulum |
| FAT_(L/R) | Frontal aslant tract |
| FPT_(L/R) | Fronto-pontine tract |
| FX_(L/R) | Fornix |
| ICP_(L/R) | Inferior cerebellar peduncle |
| IFOF_(L/R) | Inferior fronto-occipital fasciculus |
| ILF_(L/R) | Inferior longitudinal fasciculus |
| MCP_(L/R) | Middle cerebellar peduncle |
| MdLF_(L/R) | Middle longitudinal fascicle |
| OR_ML_(L/R) | Optic radiation and Meyer’s loop |
| PC | Posterior commisure |
| POPT_(L/R) | Parieto-occipito pontine tract |
| PYT_(L/R) | Pyramidal tract |
| SCP_(L/R) | Superior cerebellar peduncle |
| SLF_(L/R) | Superior longitudinal fasciculus |
| UF_(L/R) | Uncinate fasciculus |

**Supplementary Table 5.** WM bundles definition automatically extracted using the bundling profile. The bundles were defined using the BundleSeg atlas (<https://zenodo.org/records/10103446>). (L/R): left and right bundles.

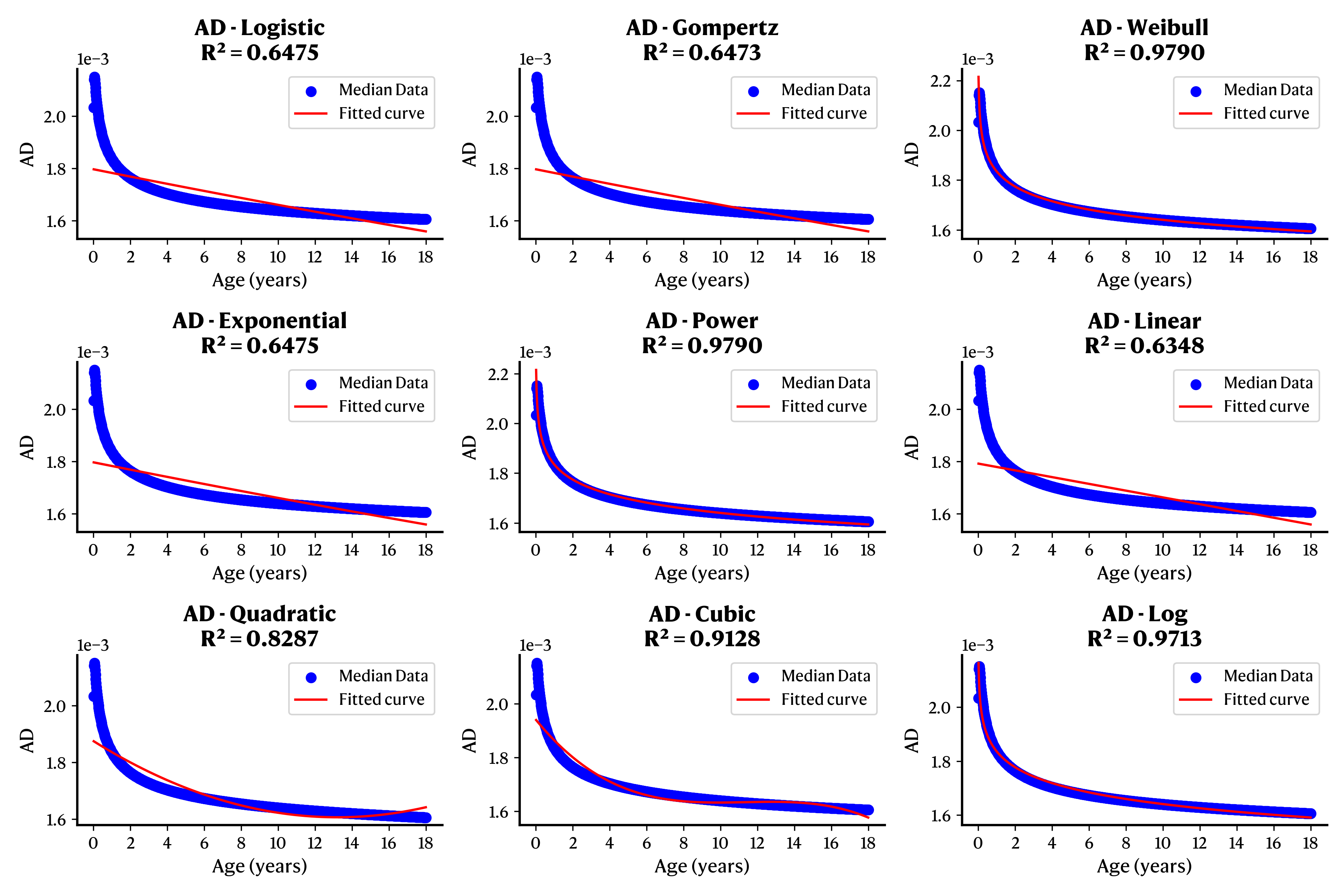

**Supplementary Figure 1.** AD median data approximation using a simpler equation as a function of age. Each scatter plot shows one of the tested equations and its associated R^2^ value.

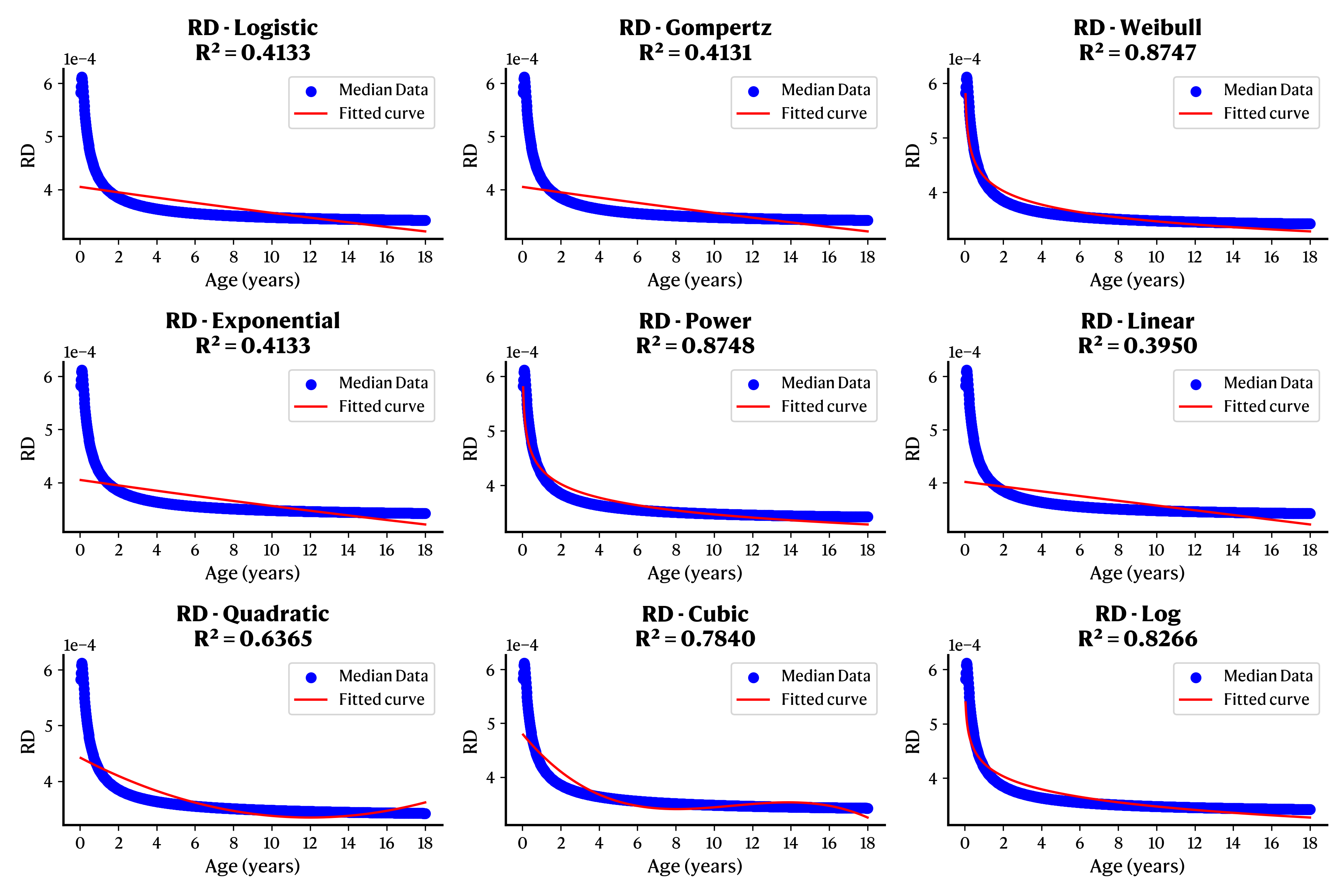

**Supplementary Figure 2.** RD median data approximation using simpler equations as a function of age. Each scatter plot shows one of the tested equations and its associated R^2^ value.

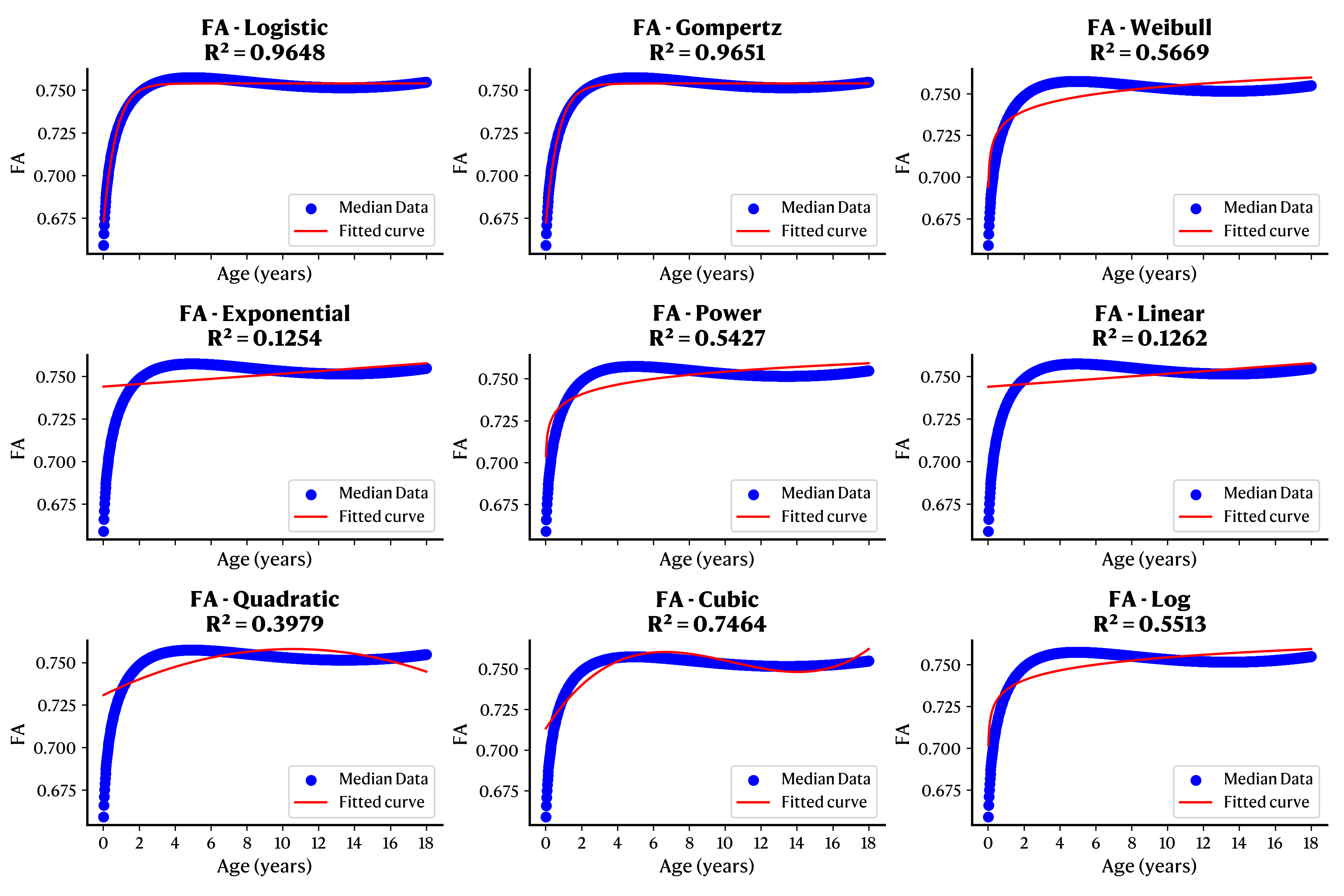

**Supplementary Figure 3.** FA median data approximation using simpler equations as a function of age. Each scatter plot shows one of the tested equations and its associated R^2^ value.

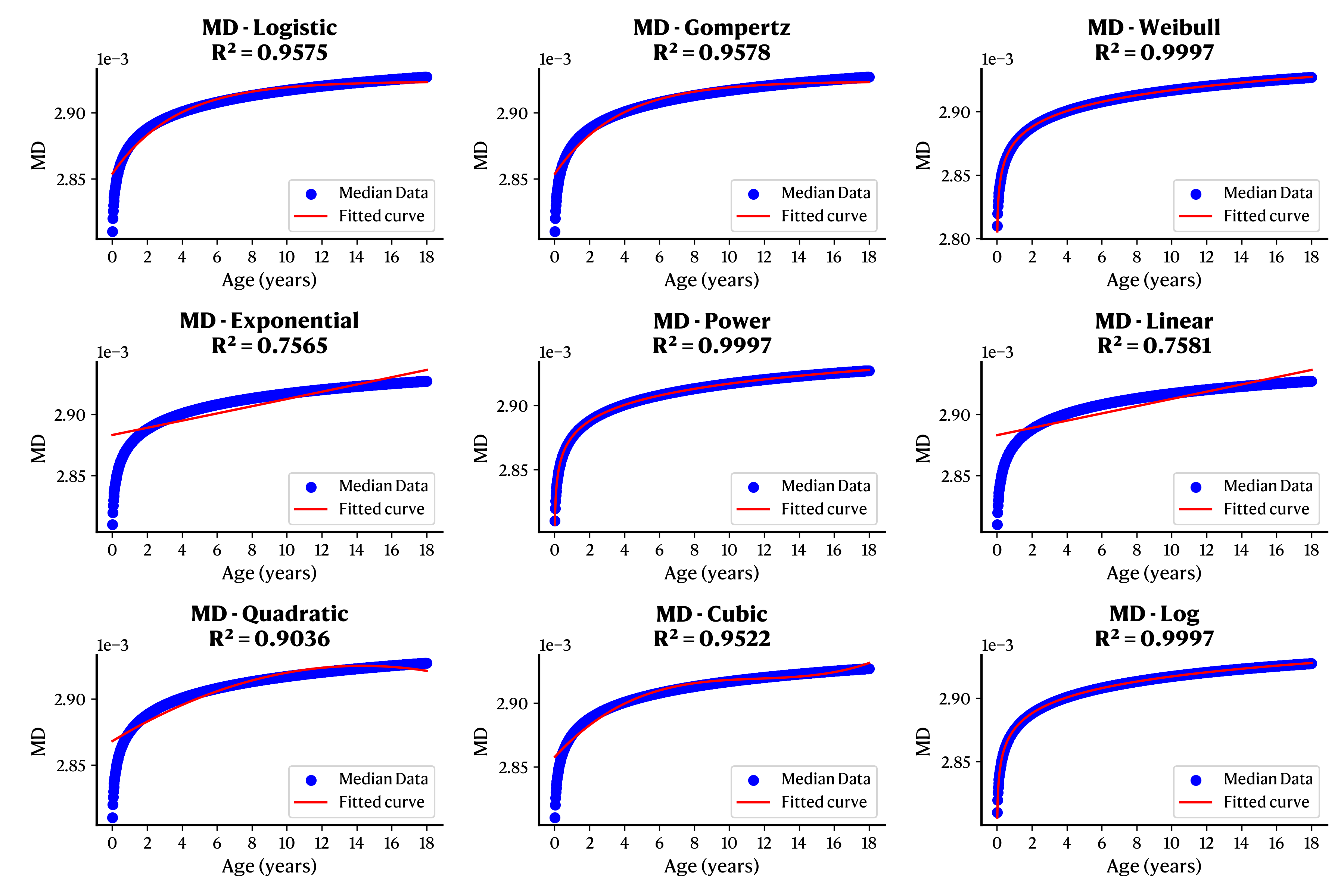

**Supplementary Figure 4.** MD median data approximation using simpler equations as a function of age. Each scatter plot shows one of the tested equations and its associated R^2^ value.

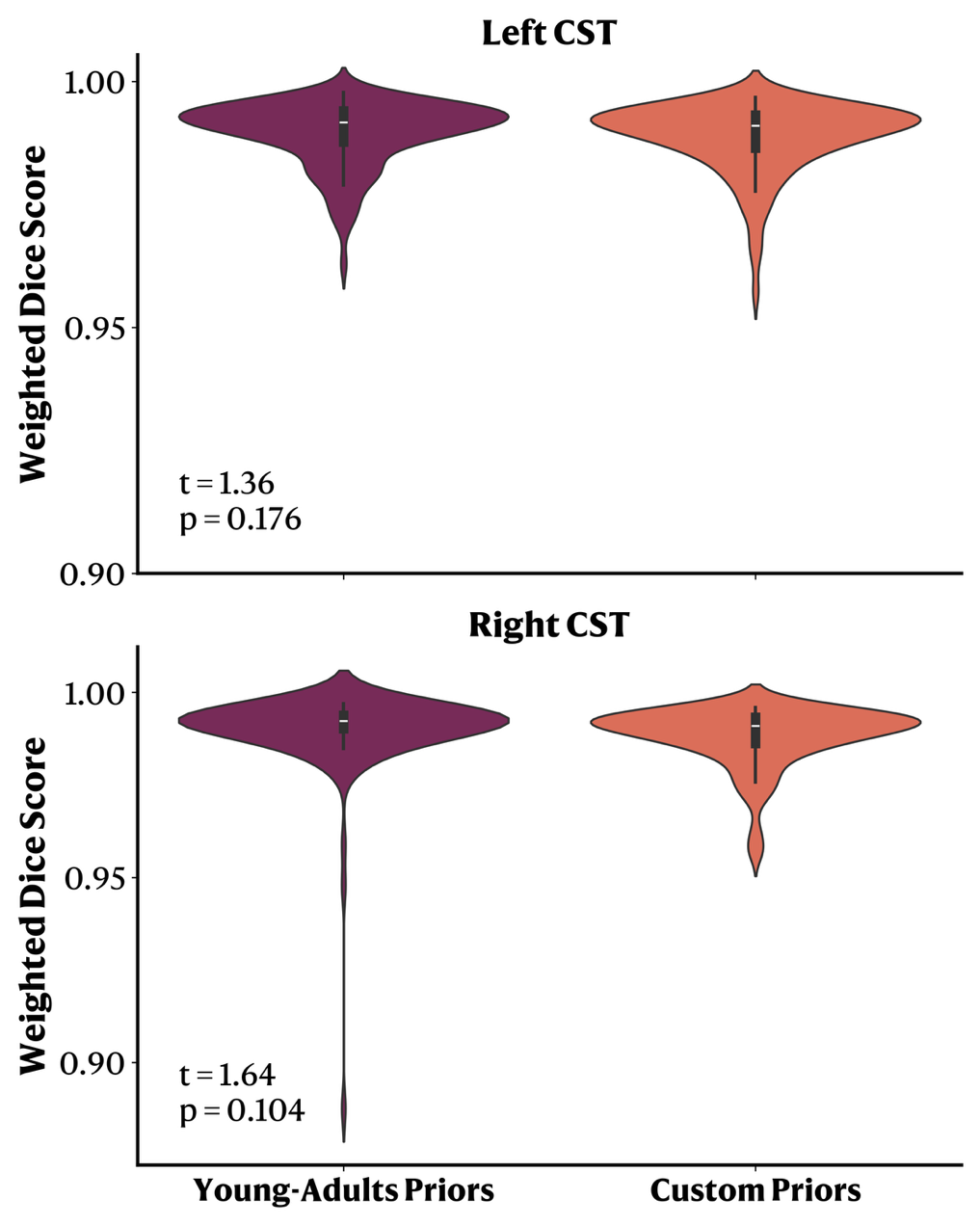

**Supplementary Figure 5.** Weighted dice score computed in test-retest acquisitions in the MASiVar dataset before and after the inclusion of the age-adaptable priors. t: t-statistic from a two-tailed t-test. p: p-value.

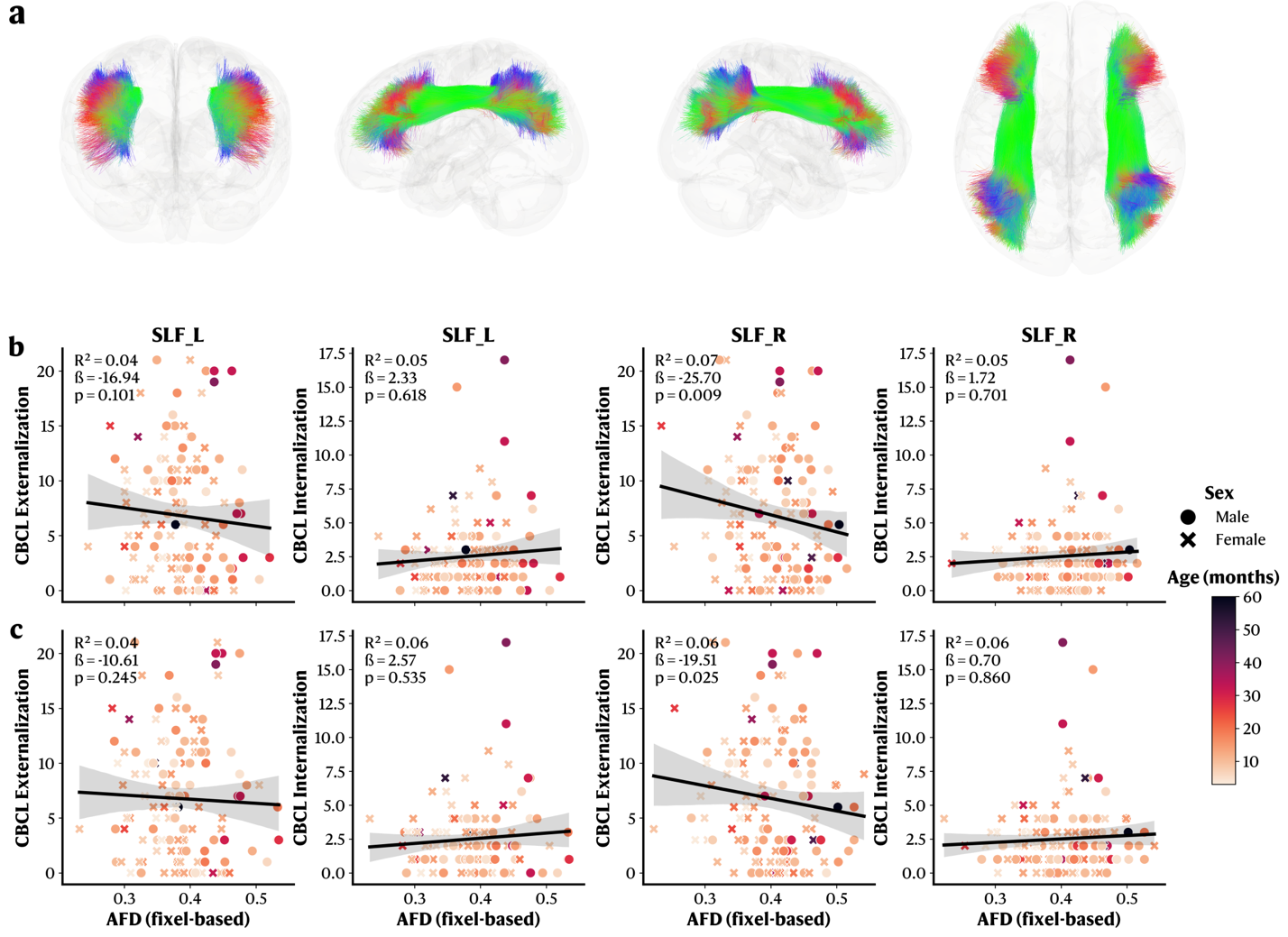

**Supplementary Figure 6.** Age-adaptable priors do not introduce significant brain-behavior relationships. **a.** Visual representation of the left and right SLF in a glass brain. **b-c.** Multivariate regression results between fixel-based average fiber density (AFD Fixel) on the superior longitudinal fasciculus (SLF) and raw externalization and internalization scores in the BCP cohort before and after the inclusion of the custom age-adaptable priors. **b.** Scatter plots representing the association between raw externalization and raw internalization scores and AFD Fixel on both left and right SLF using the **young-adult priors.** **c.** Scatter plots representing the association between raw externalization and internalization scores and AFD Fixel on both left and right SLF using the **age-adaptable priors**. ß: Beta coefficients for the AFD Fixel variable. p: p-value for the AFD fixel variable. CBCL: Child Behavioral Checklist.

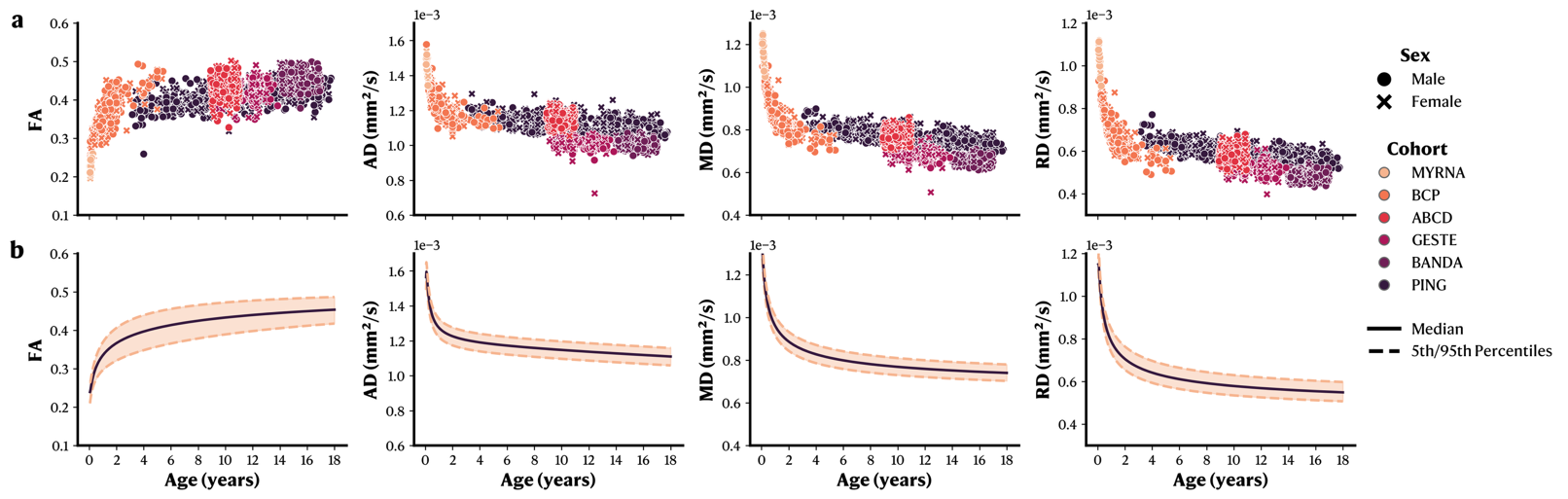

**Supplementary Figure 7.** WM microstructural changes across the developmental age range on the left arcuate fasciculus (AF_L). **a.** Raw data for each metric in function of age, stratified by sex and cohort. **b.** GAMLSS normative models with median and 5^th^/95^th^ percentiles. FA: Fractional Anisotropy. AD: Axial Diffusivity. RD: Radial Diffusivity. MD: Mean Diffusivity.

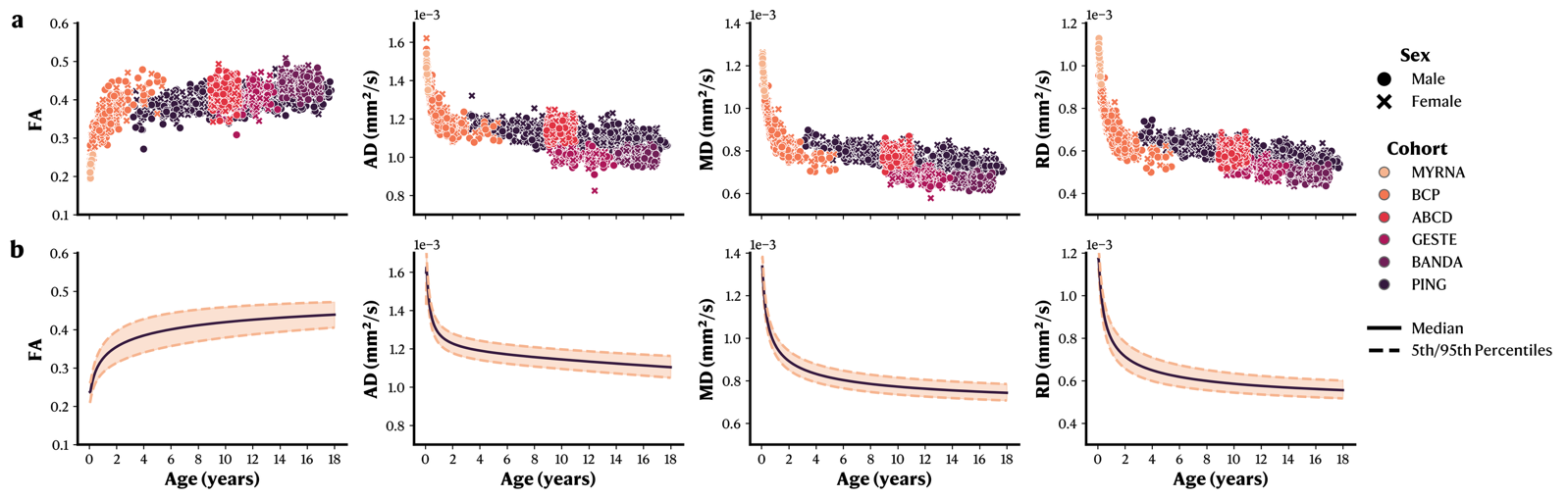

**Supplementary Figure 8.** WM microstructural changes across the developmental age range on the right arcuate fasciculus (AF_R). **a.** Raw data for each metric in function of age, stratified by sex and cohort. **b.** GAMLSS normative models with median and 5^th^/95^th^ percentiles. FA: Fractional Anisotropy. AD: Axial Diffusivity. RD: Radial Diffusivity. MD: Mean Diffusivity.

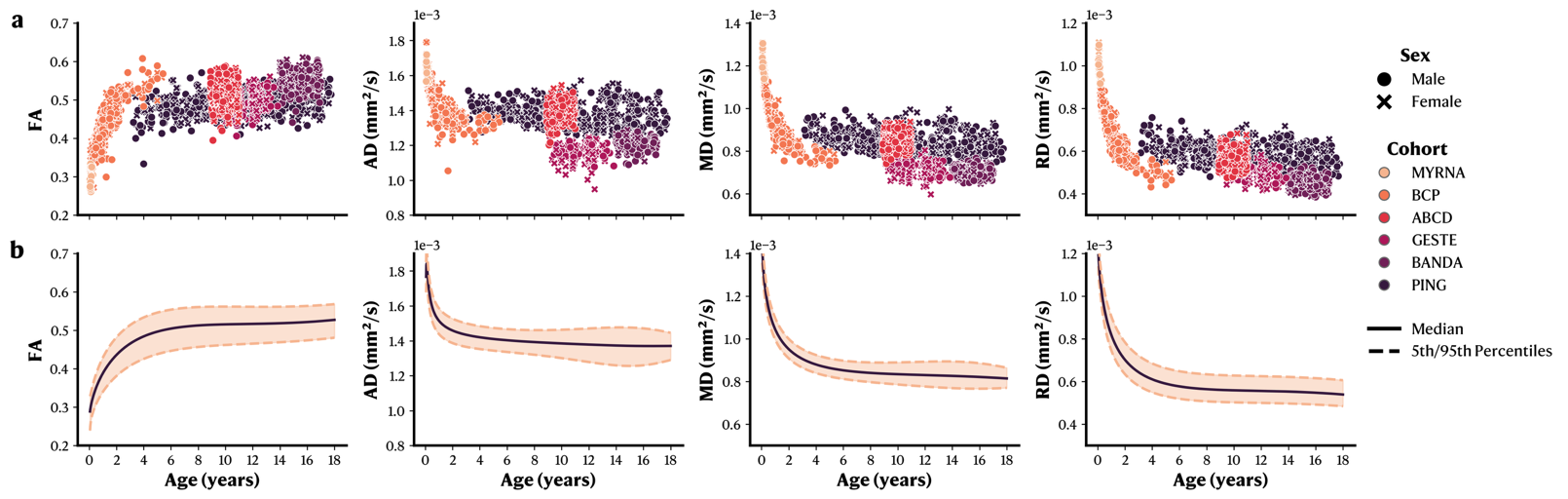

**Supplementary Figure 9.** WM microstructural changes across the developmental age range on the anterior frontal part of the corpus callosum (CC_Fr_1). **a.** Raw data for each metric in function of age, stratified by sex and cohort. **b.** GAMLSS normative models with median and 5^th^/95^th^ percentiles. FA: Fractional Anisotropy. AD: Axial Diffusivity. RD: Radial Diffusivity. MD: Mean Diffusivity.

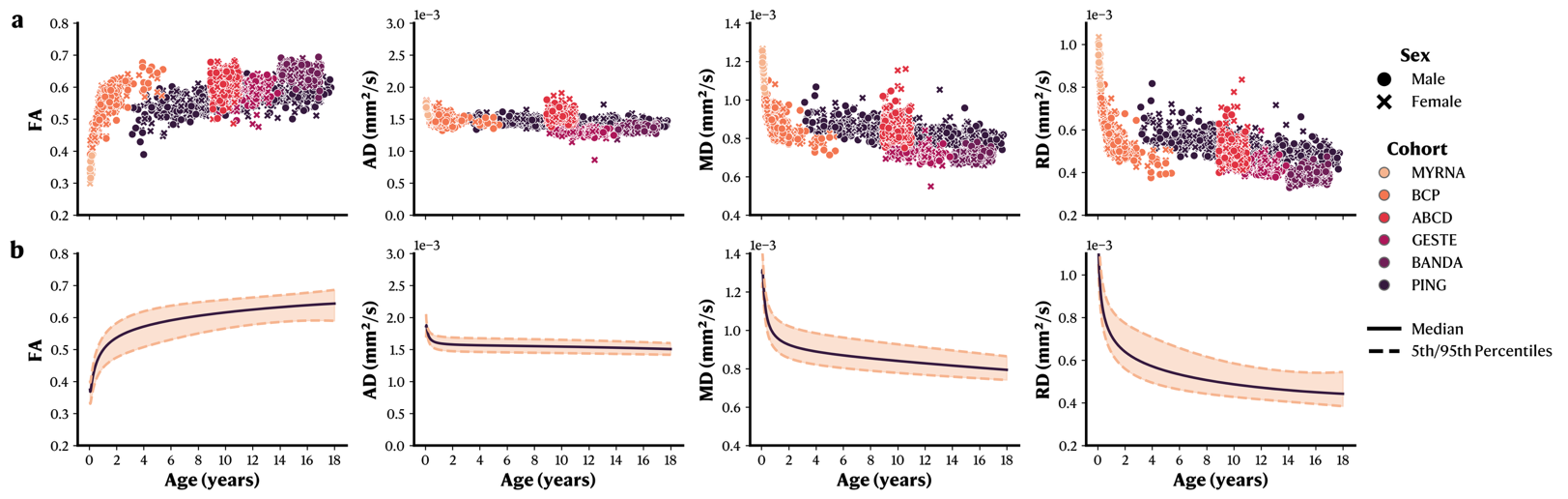

**Supplementary Figure 10.** WM microstructural changes across the developmental age range on the occipital part of the corpus callosum (CC_Oc). **a.** Raw data for each metric in function of age, stratified by sex and cohort. **b.** GAMLSS normative models with median and 5^th^/95^th^ percentiles. FA: Fractional Anisotropy. AD: Axial Diffusivity. RD: Radial Diffusivity. MD: Mean Diffusivity.

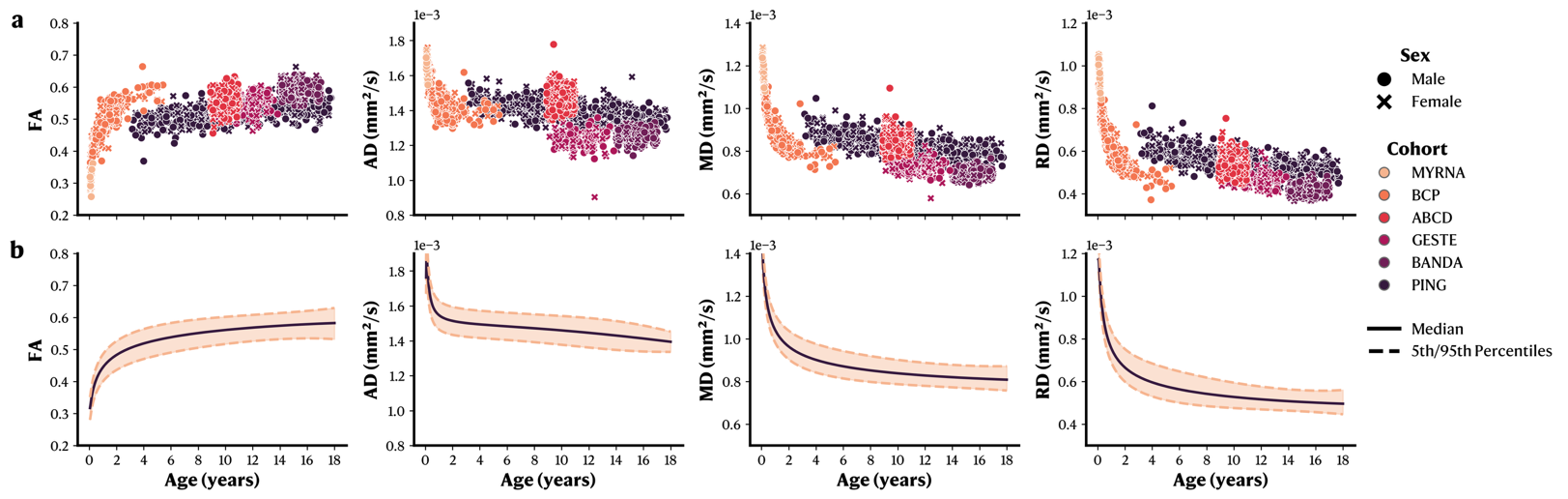

**Supplementary Figure 11.** WM microstructural changes across the developmental age range on the parietal part of the corpus callosum (CC_Pa). **a.** Raw data for each metric in function of age, stratified by sex and cohort. **b.** GAMLSS normative models with median and 5^th^/95^th^ percentiles. FA: Fractional Anisotropy. AD: Axial Diffusivity. RD: Radial Diffusivity. MD: Mean Diffusivity.

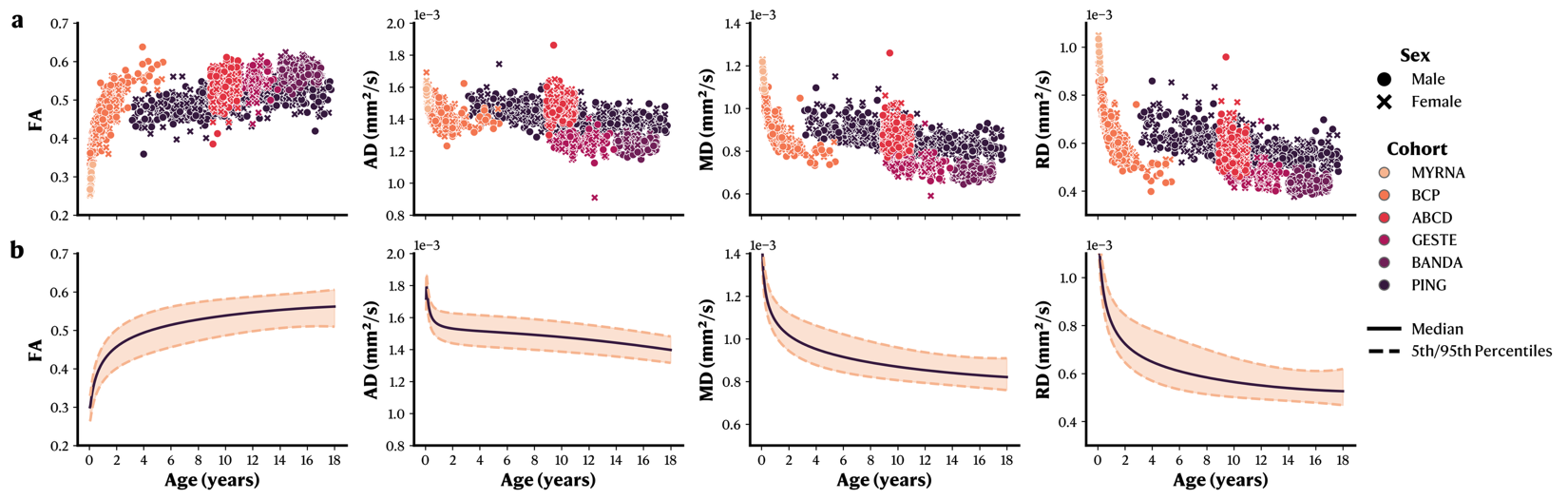

**Supplementary Figure 12.** WM microstructural changes across the developmental age range on the pre/post central gyri part of the corpus callosum (CC_Pr_Po). **a.** Raw data for each metric in function of age, stratified by sex and cohort. **b.** GAMLSS normative models with median and 5^th^/95^th^ percentiles. FA: Fractional Anisotropy. AD: Axial Diffusivity. RD: Radial Diffusivity. MD: Mean Diffusivity.

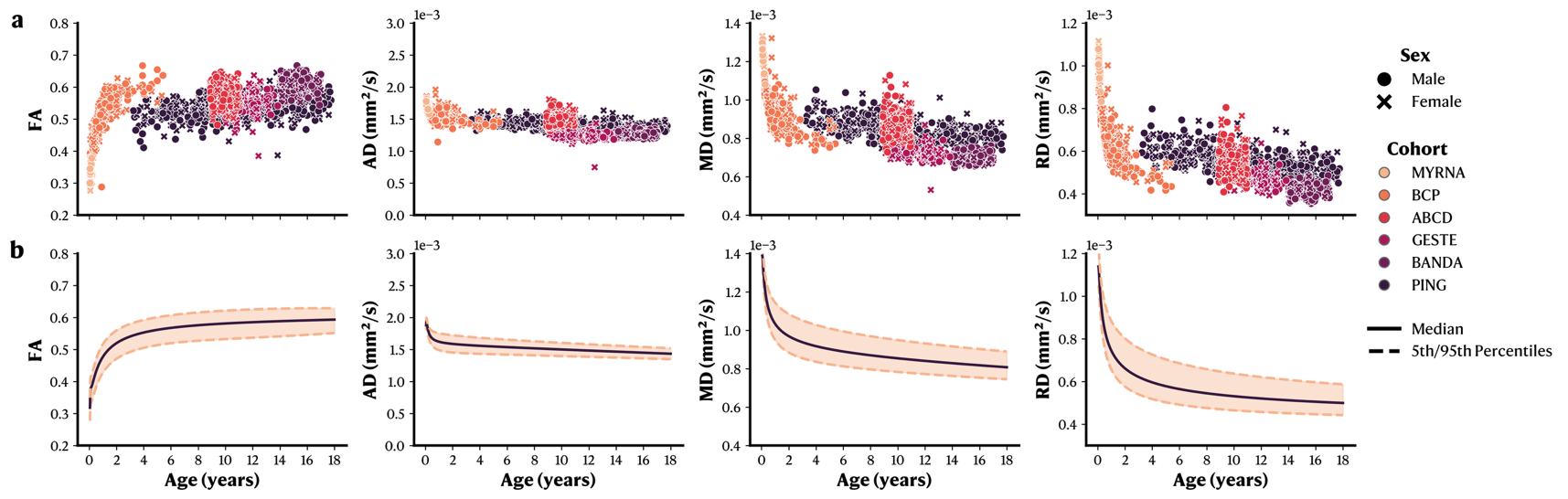

**Supplementary Figure 13.** WM microstructural changes across the developmental age range on the temporal part of the corpus callosum (CC_Te). **a.** Raw data for each metric in function of age, stratified by sex and cohort. **b.** GAMLSS normative models with median and 5^th^/95^th^ percentiles. FA: Fractional Anisotropy. AD: Axial Diffusivity. RD: Radial Diffusivity. MD: Mean Diffusivity.

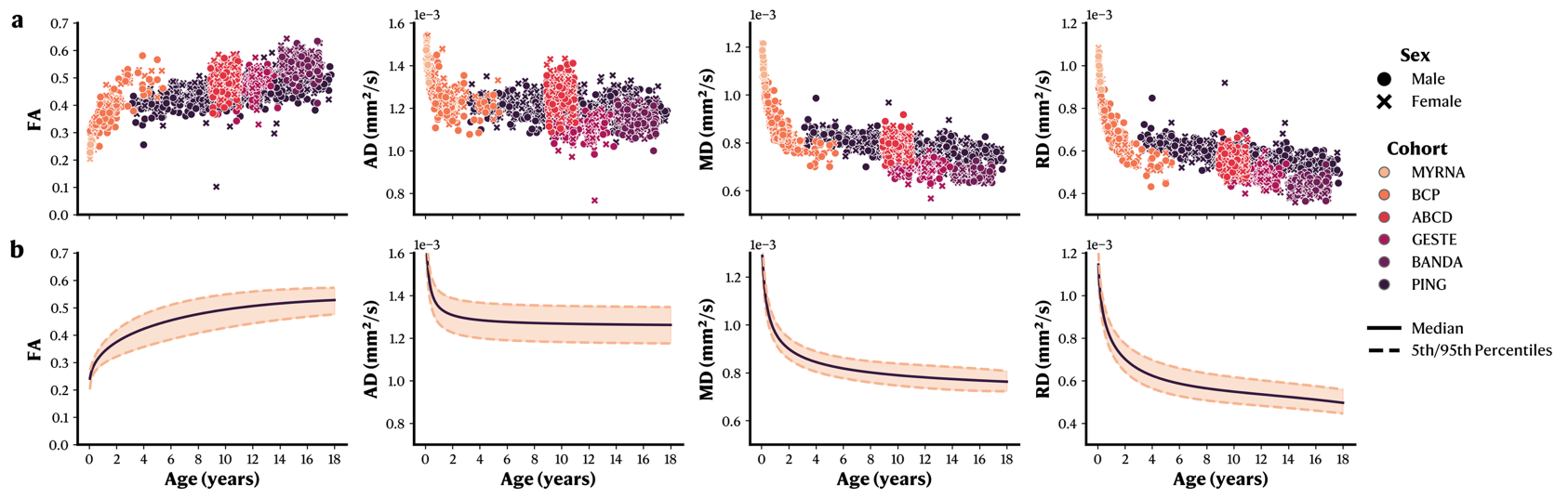

**Supplementary Figure 14.** WM microstructural changes across the developmental age range on the left cingulum (CG_L). **a.** Raw data for each metric in function of age, stratified by sex and cohort. **b.** GAMLSS normative models with median and 5^th^/95^th^ percentiles. FA: Fractional Anisotropy. AD: Axial Diffusivity. RD: Radial Diffusivity. MD: Mean Diffusivity.

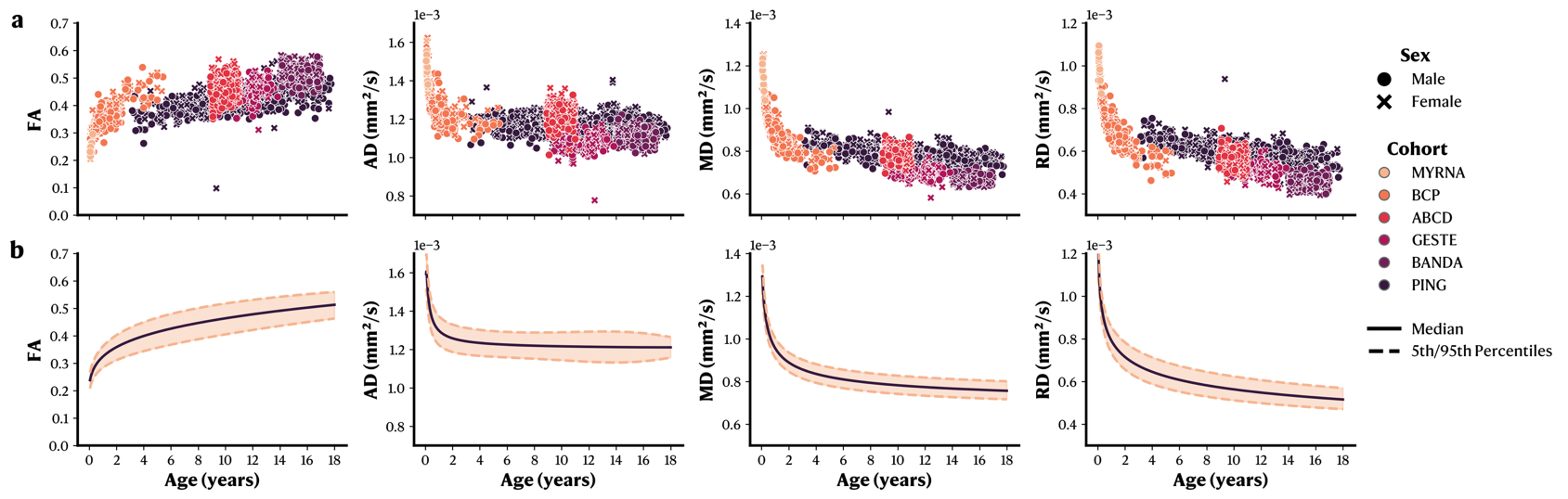

**Supplementary Figure 15.** WM microstructural changes across the developmental age range on the right cingulum (CG_R). **a.** Raw data for each metric in function of age, stratified by sex and cohort. **b.** GAMLSS normative models with median and 5^th^/95^th^ percentiles. FA: Fractional Anisotropy. AD: Axial Diffusivity. RD: Radial Diffusivity. MD: Mean Diffusivity.

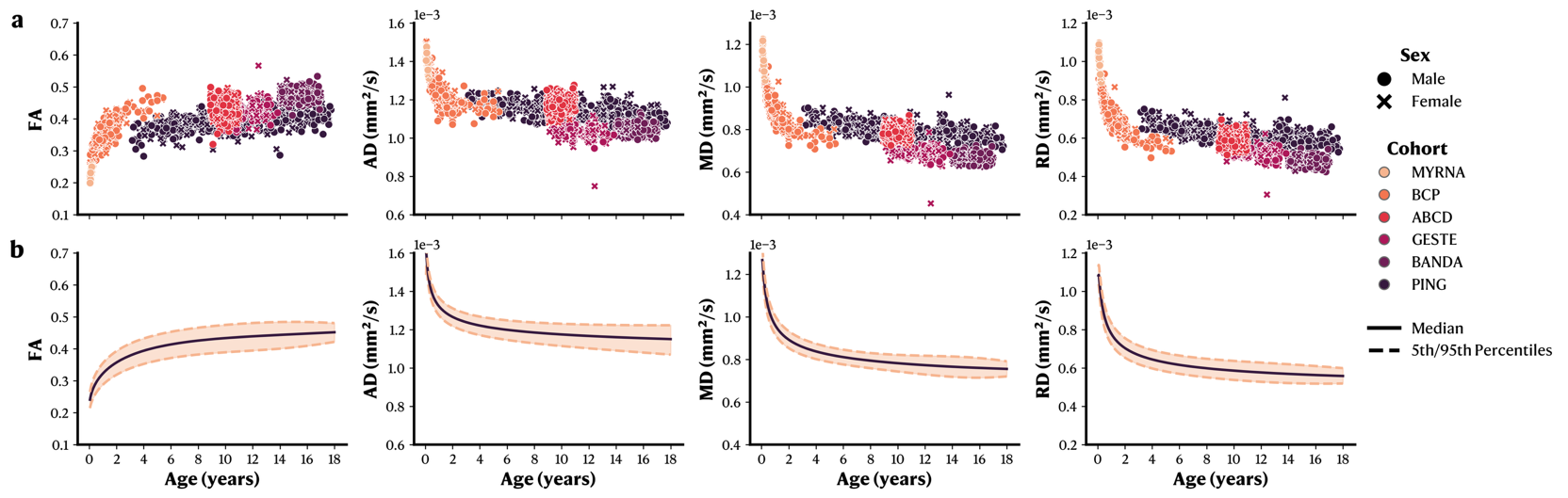

**Supplementary Figure 16.** WM microstructural changes across the developmental age range on the left frontal aslant tract (FAT_L). **a.** Raw data for each metric in function of age, stratified by sex and cohort. **b.** GAMLSS normative models with median and 5^th^/95^th^ percentiles. FA: Fractional Anisotropy. AD: Axial Diffusivity. RD: Radial Diffusivity. MD: Mean Diffusivity.

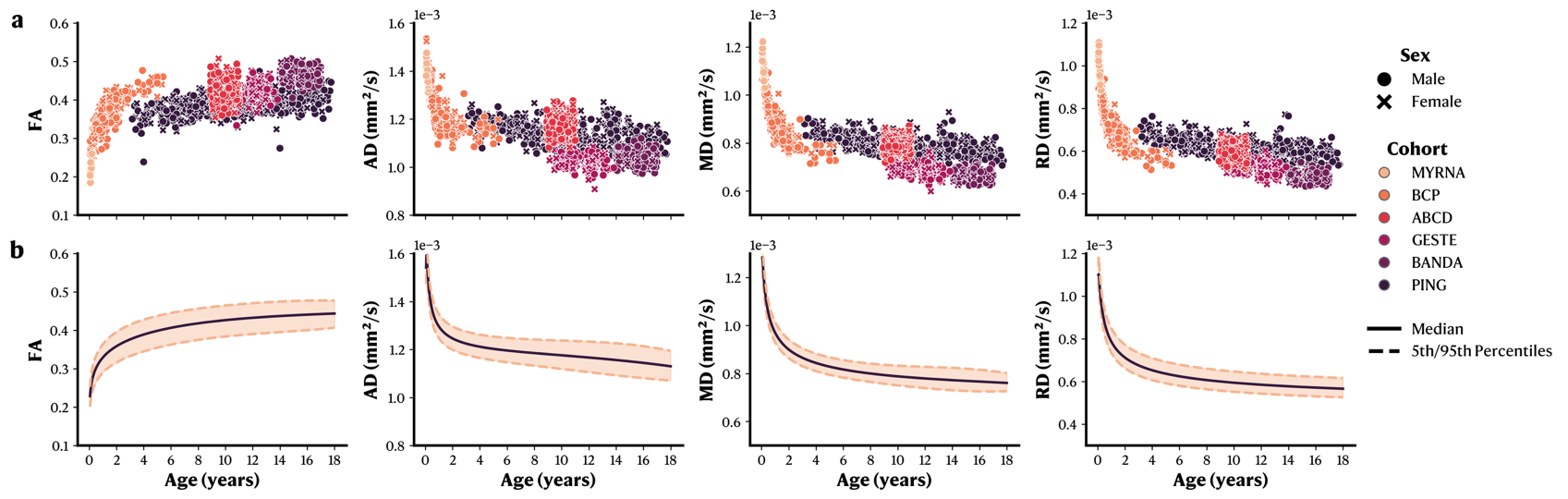

**Supplementary Figure 17.** WM microstructural changes across the developmental age range on the right frontal aslant tract (FAT_R). **a.** Raw data for each metric in function of age, stratified by sex and cohort. **b.** GAMLSS normative models with median and 5^th^/95^th^ percentiles. FA: Fractional Anisotropy. AD: Axial Diffusivity. RD: Radial Diffusivity. MD: Mean Diffusivity.

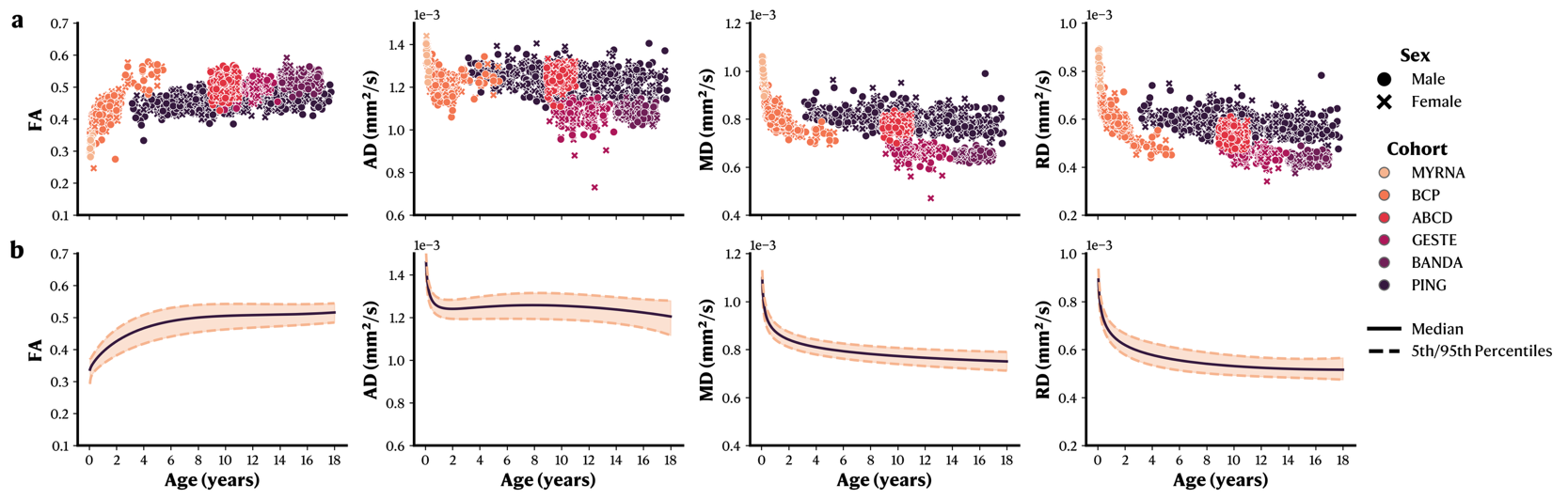

**Supplementary Figure 18.** WM microstructural changes across the developmental age range on the left fronto-pontine tract (FPT_L). **a.** Raw data for each metric in function of age, stratified by sex and cohort. **b.** GAMLSS normative models with median and 5^th^/95^th^ percentiles. FA: Fractional Anisotropy. AD: Axial Diffusivity. RD: Radial Diffusivity. MD: Mean Diffusivity.

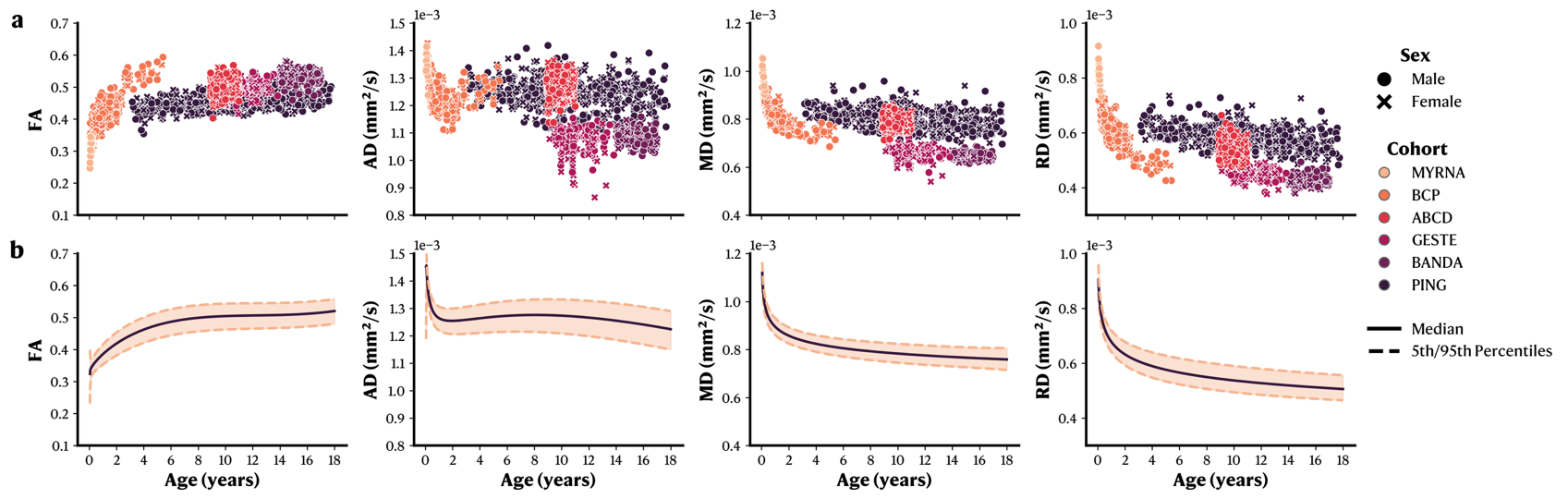

**Supplementary Figure 19.** WM microstructural changes across the developmental age range on the right fronto-pontine tract (FPT_R). **a.** Raw data for each metric in function of age, stratified by sex and cohort. **b.** GAMLSS normative models with median and 5^th^/95^th^ percentiles. FA: Fractional Anisotropy. AD: Axial Diffusivity. RD: Radial Diffusivity. MD: Mean Diffusivity.

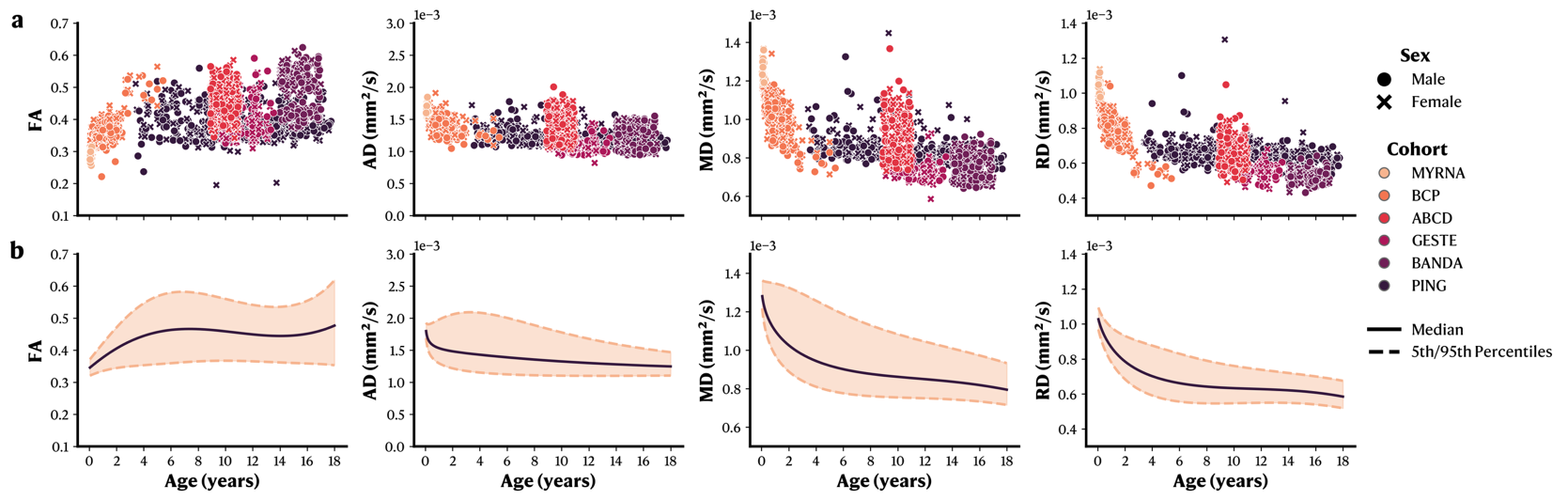

**Supplementary Figure 20.** WM microstructural changes across the developmental age range on the left fornix (FX_L). **a.** Raw data for each metric in function of age, stratified by sex and cohort. **b.** GAMLSS normative models with median and 5^th^/95^th^ percentiles. FA: Fractional Anisotropy. AD: Axial Diffusivity. RD: Radial Diffusivity. MD: Mean Diffusivity.

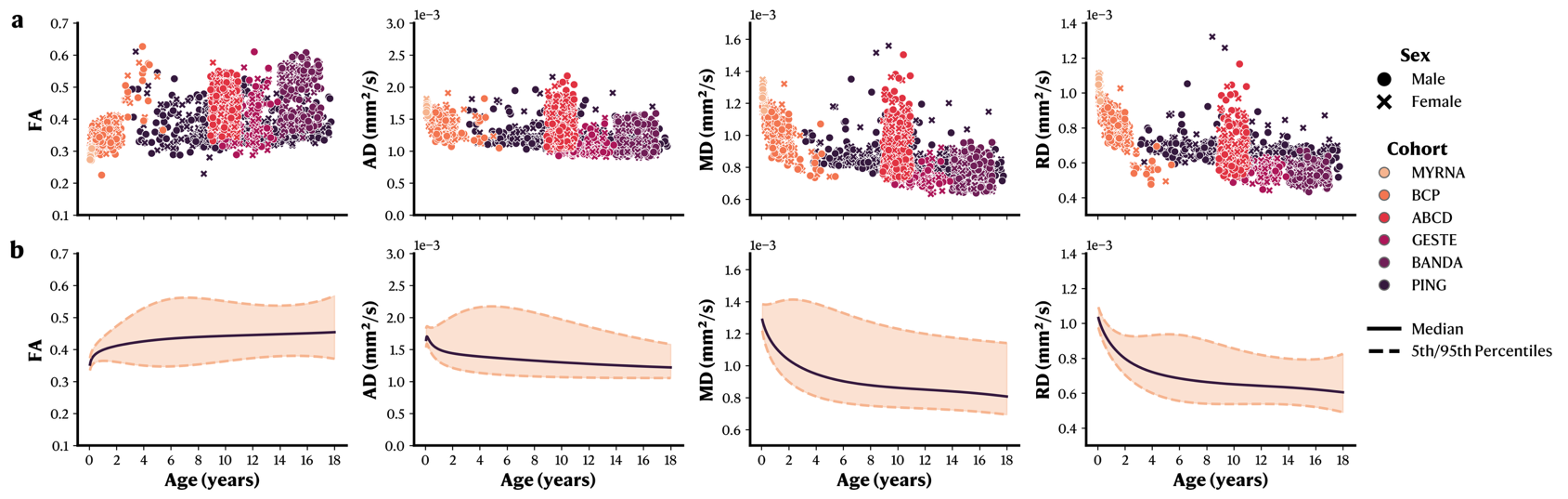

**Supplementary Figure 21.** WM microstructural changes across the developmental age range on the right fornix (FX_R). **a.** Raw data for each metric in function of age, stratified by sex and cohort. **b.** GAMLSS normative models with median and 5^th^/95^th^ percentiles. FA: Fractional Anisotropy. AD: Axial Diffusivity. RD: Radial Diffusivity. MD: Mean Diffusivity.

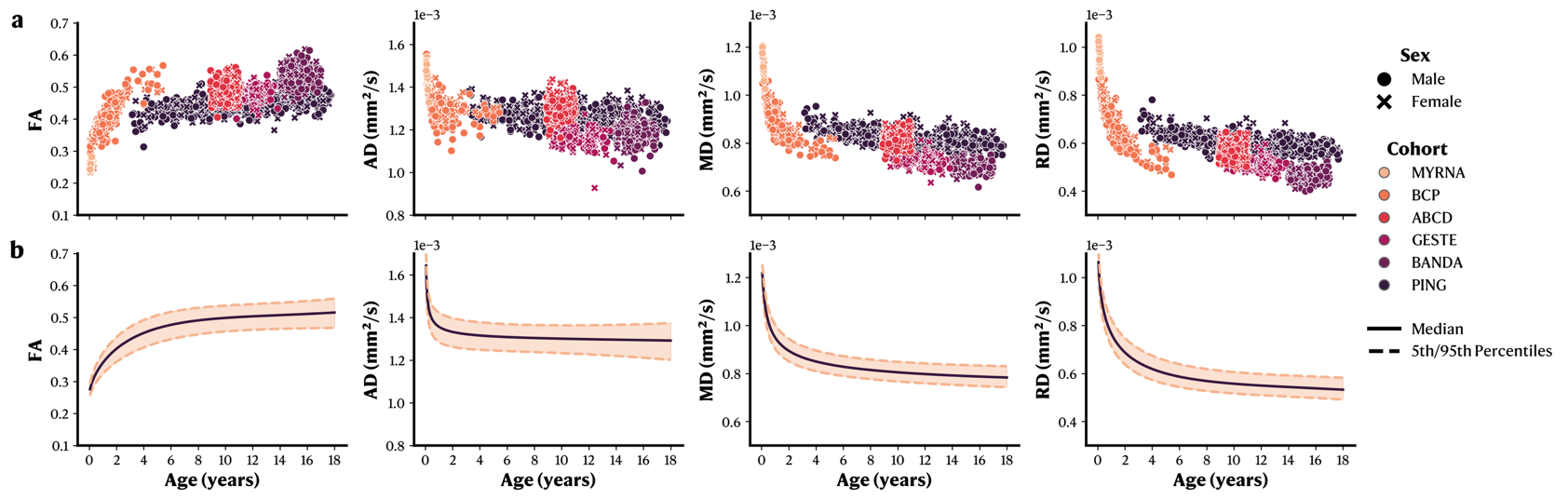

**Supplementary Figure 22.** WM microstructural changes across the developmental age range on the left inferior fronto-occipital fasciculus (IFOF_L). **a.** Raw data for each metric in function of age, stratified by sex and cohort. **b.** GAMLSS normative models with median and 5^th^/95^th^ percentiles. FA: Fractional Anisotropy. AD: Axial Diffusivity. RD: Radial Diffusivity. MD: Mean Diffusivity.

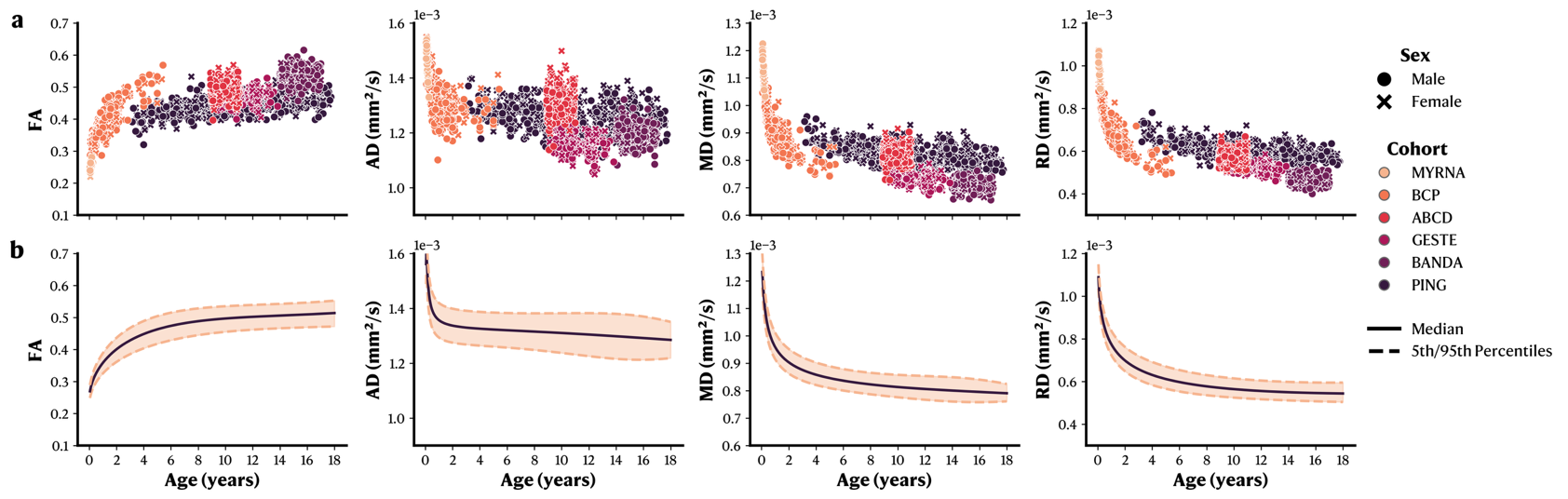

**Supplementary Figure 23.** WM microstructural changes across the developmental age range on the right inferior fronto-occipital fasciculus (IFOF_R). **a.** Raw data for each metric in function of age, stratified by sex and cohort. **b.** GAMLSS normative models with median and 5^th^/95^th^ percentiles. FA: Fractional Anisotropy. AD: Axial Diffusivity. RD: Radial Diffusivity. MD: Mean Diffusivity.

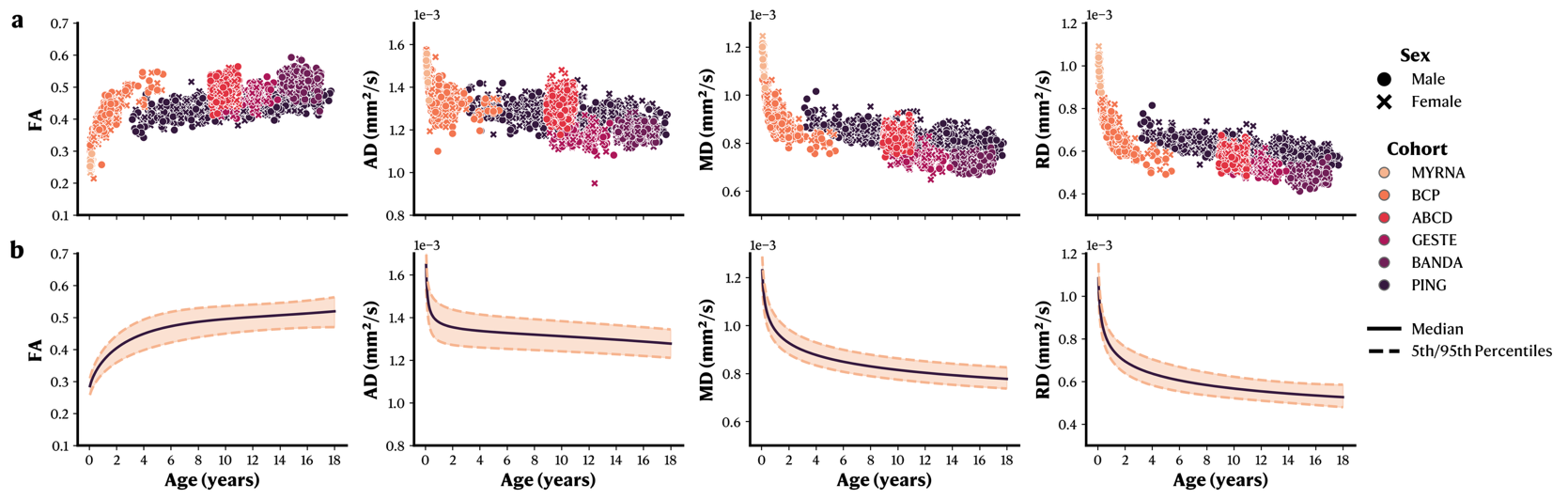

**Supplementary Figure 24.** WM microstructural changes across the developmental age range on the left inferior longitudinal fasciculus (ILF_L). **a.** Raw data for each metric in function of age, stratified by sex and cohort. **b.** GAMLSS normative models with median and 5^th^/95^th^ percentiles. FA: Fractional Anisotropy. AD: Axial Diffusivity. RD: Radial Diffusivity. MD: Mean Diffusivity.

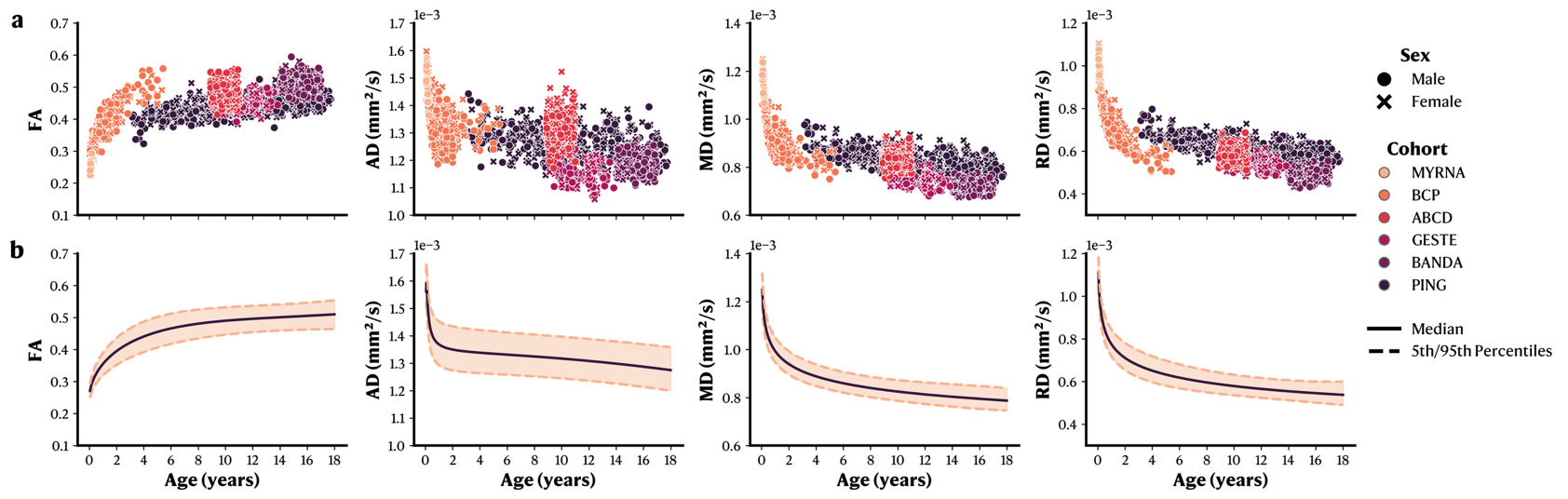

**Supplementary Figure 25.** WM microstructural changes across the developmental age range on the right inferior longitudinal fasciculus (ILF_R). **a.** Raw data for each metric in function of age, stratified by sex and cohort. **b.** GAMLSS normative models with median and 5^th^/95^th^ percentiles. FA: Fractional Anisotropy. AD: Axial Diffusivity. RD: Radial Diffusivity. MD: Mean Diffusivity.

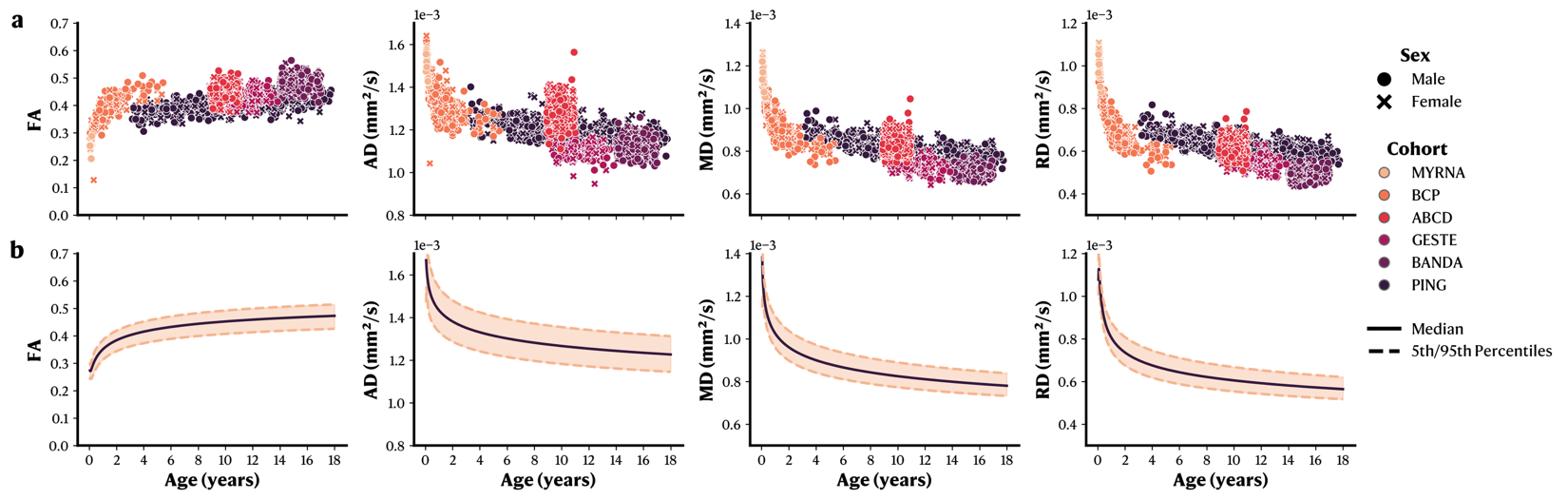

**Supplementary Figure 26.** WM microstructural changes across the developmental age range on the left middle longitudinal fascicle (MdLF_L). **a.** Raw data for each metric in function of age, stratified by sex and cohort. **b.** GAMLSS normative models with median and 5^th^/95^th^ percentiles. FA: Fractional Anisotropy. AD: Axial Diffusivity. RD: Radial Diffusivity. MD: Mean Diffusivity.

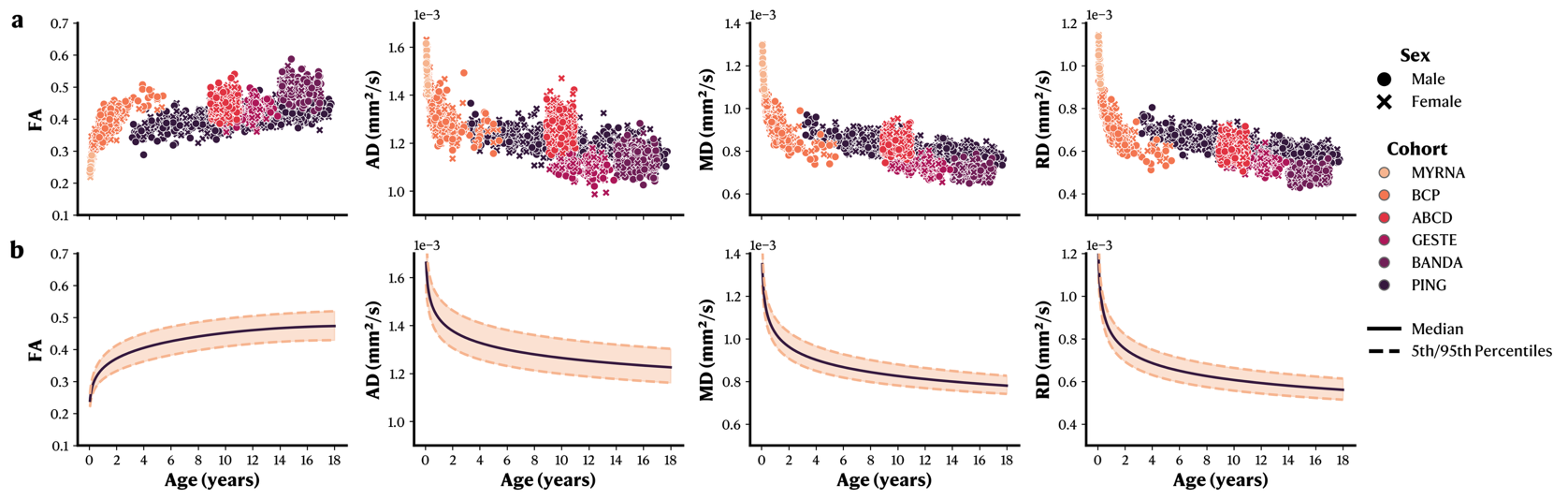

**Supplementary Figure 27.** WM microstructural changes across the developmental age range on the right middle longitudinal fascicle (MdLF_R). **a.** Raw data for each metric in function of age, stratified by sex and cohort. **b.** GAMLSS normative models with median and 5^th^/95^th^ percentiles. FA: Fractional Anisotropy. AD: Axial Diffusivity. RD: Radial Diffusivity. MD: Mean Diffusivity.

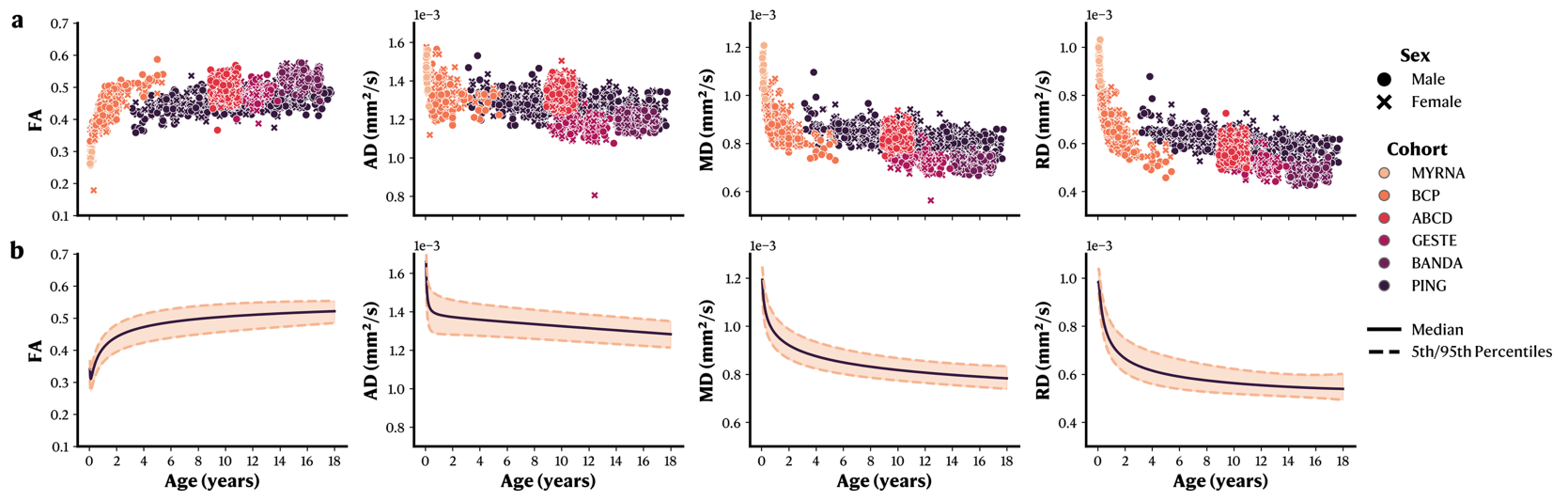

**Supplementary Figure 28.** WM microstructural changes across the developmental age range on the left optic radiation and Meyer’s loop (OR_ML_L). **a.** Raw data for each metric in function of age, stratified by sex and cohort. **b.** GAMLSS normative models with median and 5^th^/95^th^ percentiles. FA: Fractional Anisotropy. AD: Axial Diffusivity. RD: Radial Diffusivity. MD: Mean Diffusivity.

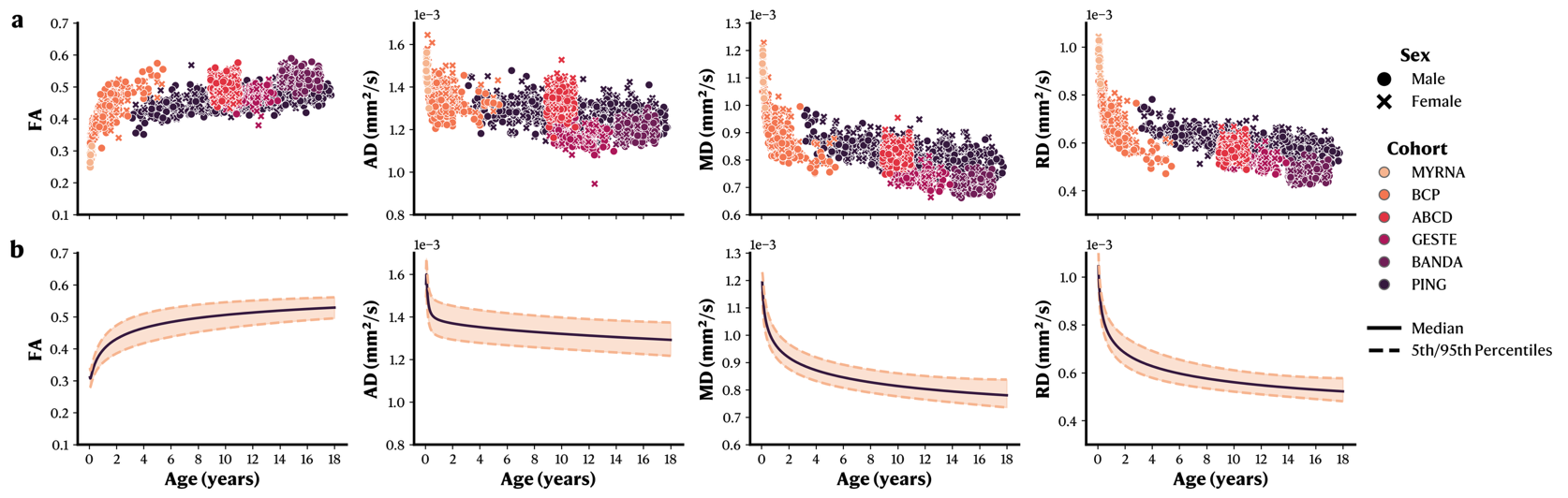

**Supplementary Figure 29.** WM microstructural changes across the developmental age range on the right optic radiation and Meyer’s loop (OR_ML_R). **a.** Raw data for each metric in function of age, stratified by sex and cohort. **b.** GAMLSS normative models with median and 5^th^/95^th^ percentiles. FA: Fractional Anisotropy. AD: Axial Diffusivity. RD: Radial Diffusivity. MD: Mean Diffusivity.

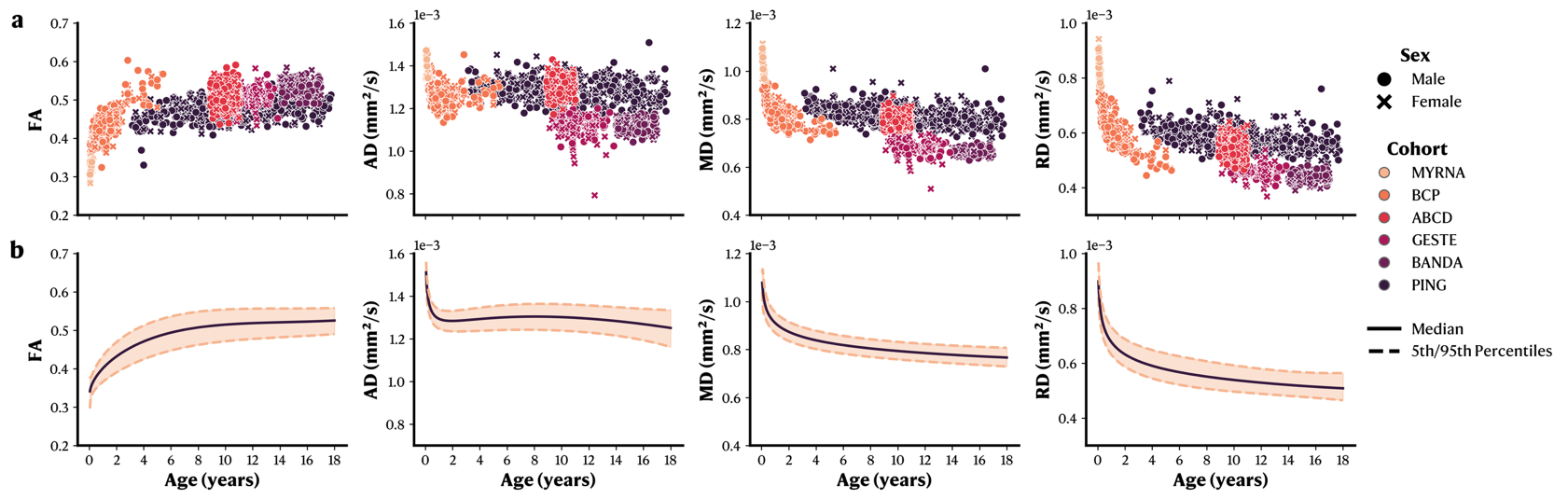

**Supplementary Figure 30.** WM microstructural changes across the developmental age range on the left parieto-occipito pontine tract (POPT_L). **a.** Raw data for each metric in function of age, stratified by sex and cohort. **b.** GAMLSS normative models with median and 5^th^/95^th^ percentiles. FA: Fractional Anisotropy. AD: Axial Diffusivity. RD: Radial Diffusivity. MD: Mean Diffusivity.

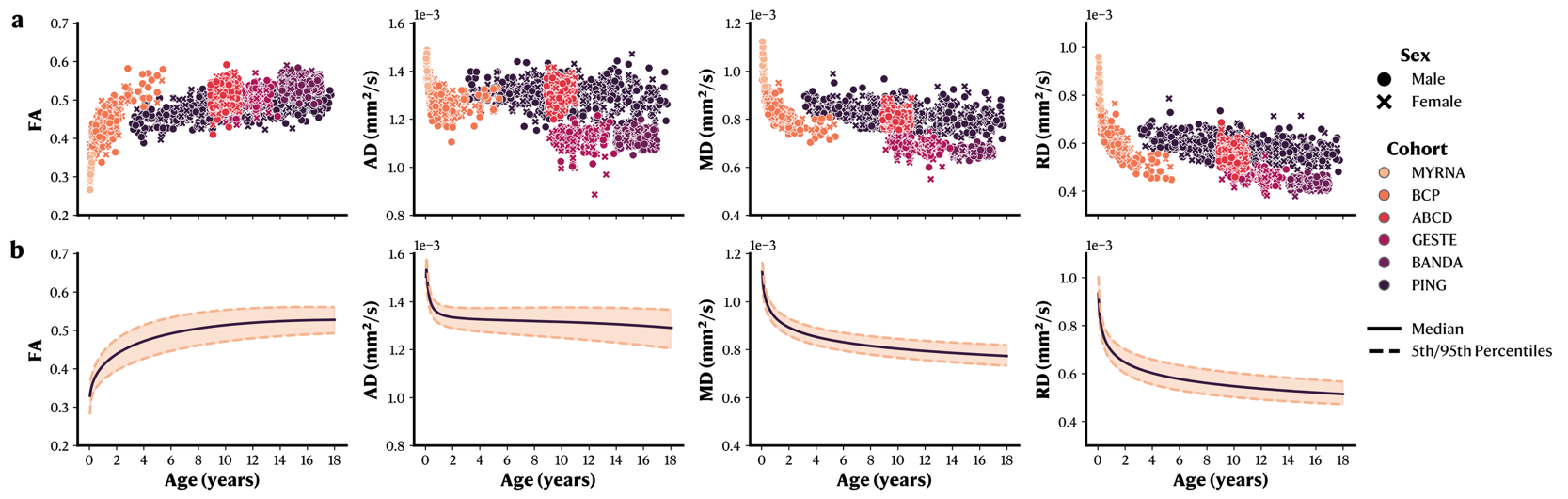

**Supplementary Figure 31.** WM microstructural changes across the developmental age range on the right parieto-occipito pontine tract (POPT_R). **a.** Raw data for each metric in function of age, stratified by sex and cohort. **b.** GAMLSS normative models with median and 5^th^/95^th^ percentiles. FA: Fractional Anisotropy. AD: Axial Diffusivity. RD: Radial Diffusivity. MD: Mean Diffusivity.

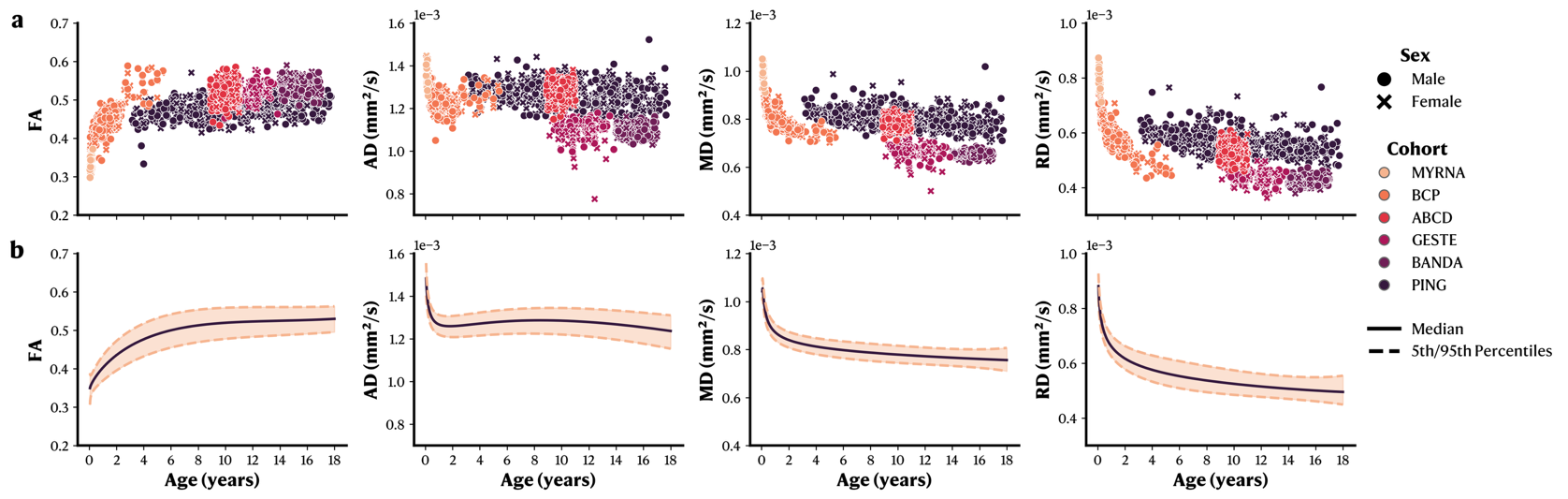

**Supplementary Figure 32.** WM microstructural changes across the developmental age range on the left pyramidal tract (PYT_L). **a.** Raw data for each metric in function of age, stratified by sex and cohort. **b.** GAMLSS normative models with median and 5^th^/95^th^ percentiles. FA: Fractional Anisotropy. AD: Axial Diffusivity. RD: Radial Diffusivity. MD: Mean Diffusivity.

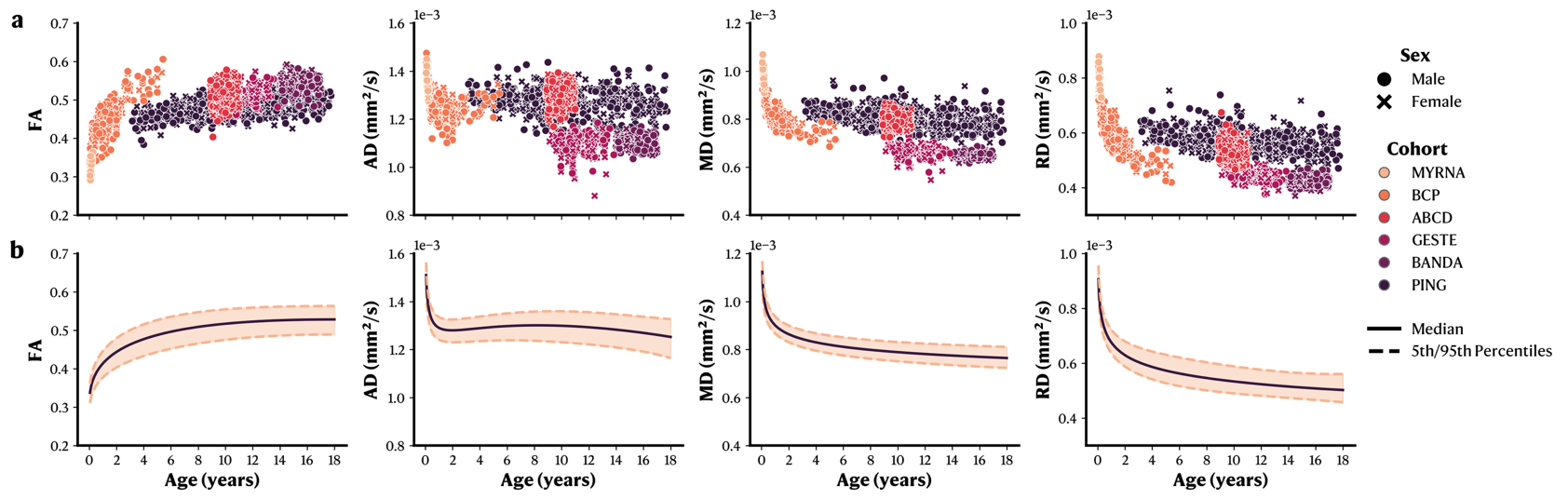

**Supplementary Figure 33.** WM microstructural changes across the developmental age range on the right pyramidal tract (PYT_R). **a.** Raw data for each metric in function of age, stratified by sex and cohort. **b.** GAMLSS normative models with median and 5^th^/95^th^ percentiles. FA: Fractional Anisotropy. AD: Axial Diffusivity. RD: Radial Diffusivity. MD: Mean Diffusivity.

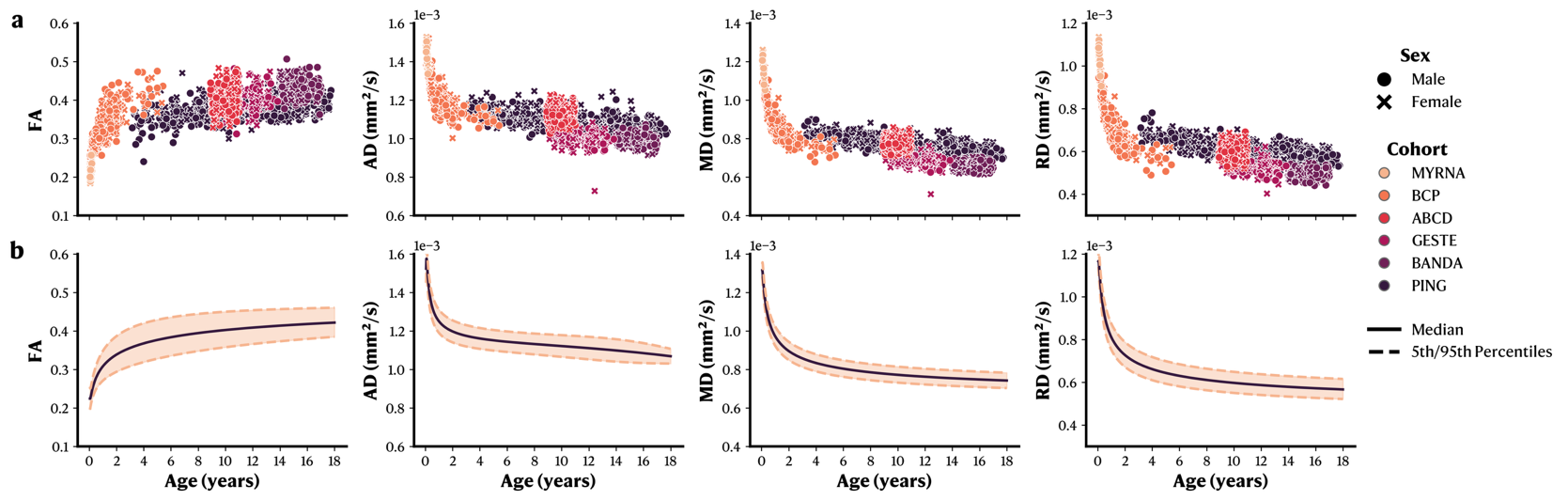

**Supplementary Figure 34.** WM microstructural changes across the developmental age range on the left superior longitudinal fasciculus (SLF_L). **a.** Raw data for each metric in function of age, stratified by sex and cohort. **b.** GAMLSS normative models with median and 5^th^/95^th^ percentiles. FA: Fractional Anisotropy. AD: Axial Diffusivity. RD: Radial Diffusivity. MD: Mean Diffusivity.

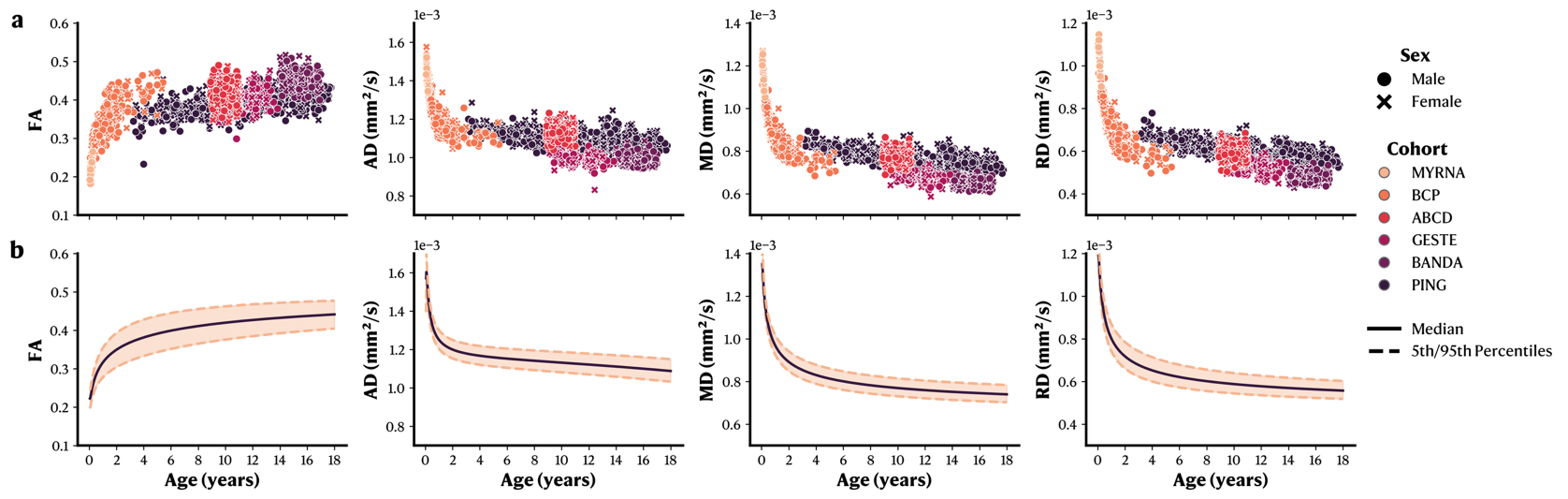

**Supplementary Figure 35.** WM microstructural changes across the developmental age range on the right superior longitudinal fasciculus (SLF_R). **a.** Raw data for each metric in function of age, stratified by sex and cohort. **b.** GAMLSS normative models with median and 5^th^/95^th^ percentiles. FA: Fractional Anisotropy. AD: Axial Diffusivity. RD: Radial Diffusivity. MD: Mean Diffusivity.

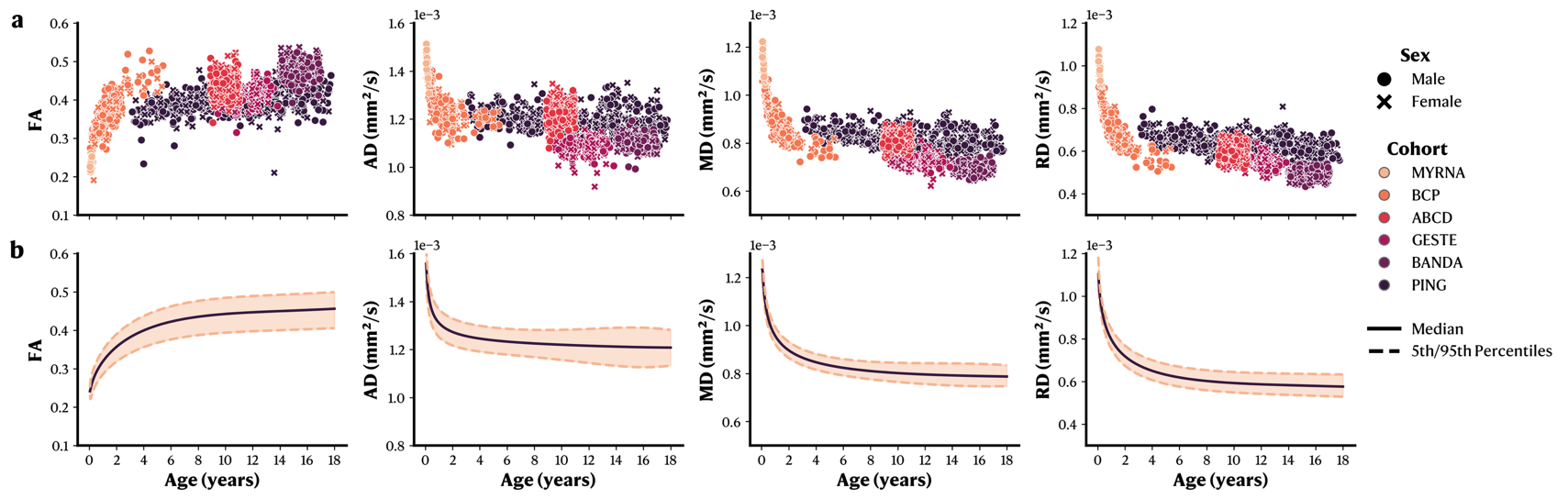

**Supplementary Figure 36.** WM microstructural changes across the developmental age range on the left uncinate fasciculus (UF_L). **a.** Raw data for each metric in function of age, stratified by sex and cohort. **b.** GAMLSS normative models with median and 5^th^/95^th^ percentiles. FA: Fractional Anisotropy. AD: Axial Diffusivity. RD: Radial Diffusivity. MD: Mean Diffusivity.

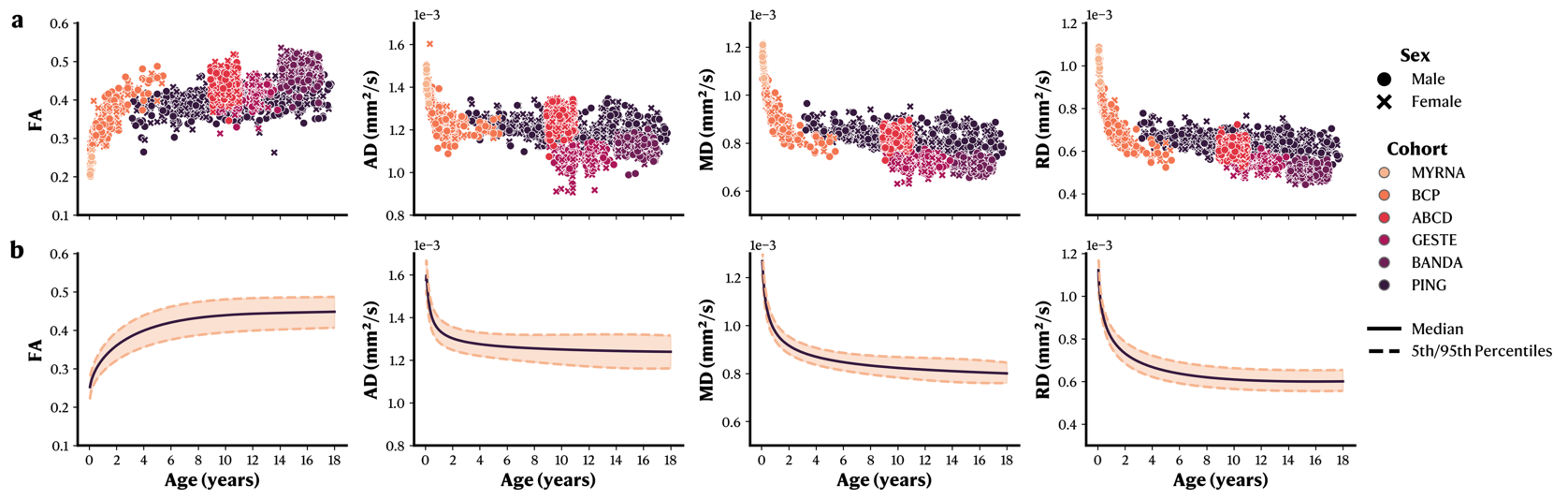

**Supplementary Figure 37.** WM microstructural changes across the developmental age range on the right uncinate fasciculus (UF_R). **a.** Raw data for each metric in function of age, stratified by sex and cohort. **b.** GAMLSS normative models with median and 5^th^/95^th^ percentiles. FA: Fractional Anisotropy. AD: Axial Diffusivity. RD: Radial Diffusivity. MD: Mean Diffusivity.

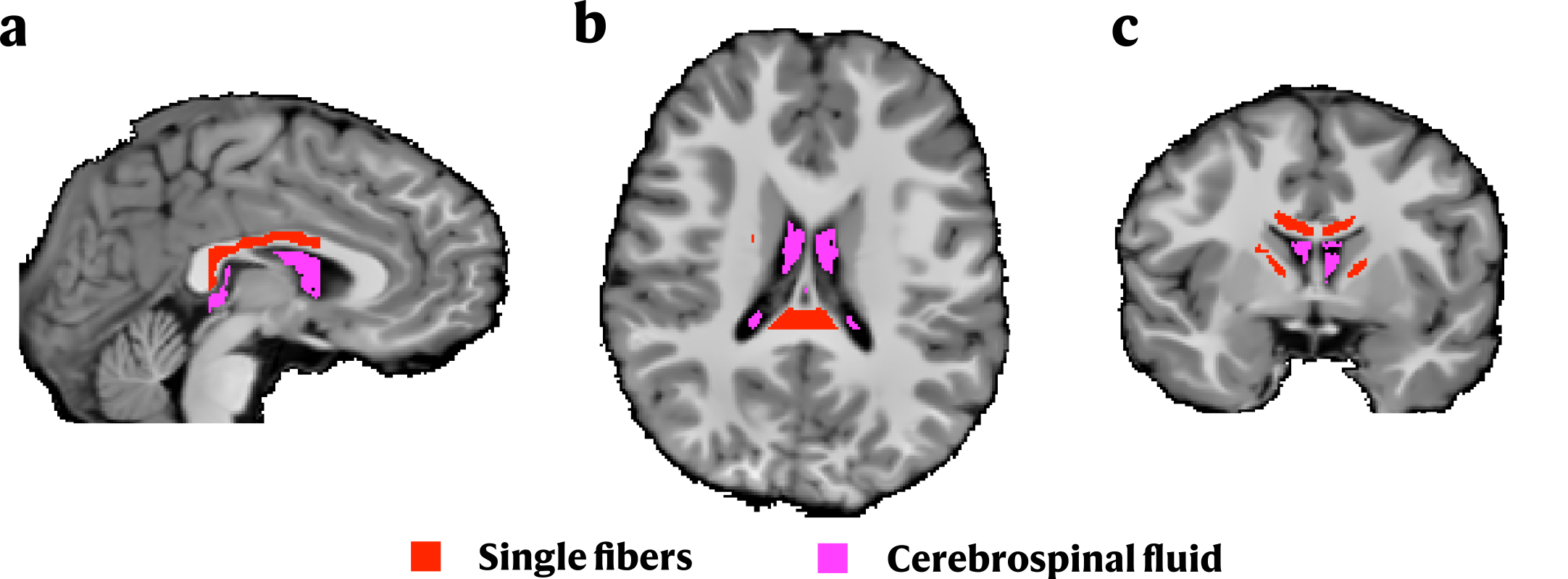

**Supplementary Figure 38.** Automatically extracted voxels representing single fibers and cerebrospinal fluid (CSF). CSF voxels are represented in pink, while single fibers voxels are represented in red. **a.** Sagittal slice. **b.** Axial slice. **c.** Coronal slice.
