## Supplementary for "sf-pediatric: A robust and age-adaptable end-to-end pipeline for pediatric diffusion MRI": SupplementaryFile1.docx

### sf-pediatric Methods Description

Suggested text and references to use when describing pipeline usage within the methods section of a publication. <https://github.com/scilus/sf-pediatric>

#### Methods

Data was processed using sf-pediatric v0.2.0 of the [Sherbrooke Connectivity Imaging Lab (SCIL)](https://github.com/scilus) , utilizing reproducible modules and subworkflows developed by the [nf-neuro](https://github.com/scilus/nf-neuro) Team.

The pipeline was executed with Nextflow v25.04.6 ([Di Tommaso et al., 2017](https://doi.org/10.1038/nbt.3820)) using apptainer with the following command:

nextflow run scilus/sf-pediatric -r dev --input /home/agagnon/projects/def-larissa1/datasets/testData/ --outdir /home/agagnon/scratch/testNODDI/nf-pediatric-0.2.0-featNODDI/ --run_freewater true --run_noddi true -profile tracking,segmentation,connectomics,bundling,apptainer,slurm --fs_license /home/agagnon/license.txt -resume

##### Processing Steps

###### DWI preprocessing

Diffusion weighting imaging (DWI) files were extracted from the input BIDS folder and associated with their corresponding reverse phase-encoded images when available. DWI volumes were denoised using the MP-PCA algorithm (Veraart et al., 2016) implemented in the MRtrix3 toolbox (Tournier et al., 2019). Susceptibility-induced distortions were corrected using FSL's TOPUP (Andersson et al., 2003; Jenkinson et al., 2012) when reverse phase-encoded images were available. Eddy current and motion correction were performed using FSL's EDDY (Andersson & Sotiropoulos, 2016; Jenkinson et al., 2012); maximum framewise displacement was recorded for quality control purposes. Brain extraction was performed by applying the deep learning model SynthStrip (Hoopes et al., 2022) on powdered average images. Pediatric-tailored weights were used for very young subjects where applicable (Kelley et al., 2024). The resulting mask was applied to the DWI volumes. Bias field correction was applied using the N4 algorithm (Tustison et al., 2010) from the ANTs toolbox (Tustison et al., 2021) using a b-spline knot per voxel of 8 and a shrink factor of 4. DWI volumes were normalized using the mean B0 intensity within white matter (FA > 0.4) using MRtrix3 (Tournier et al., 2019). Preprocessed DWI volumes were resampled to an isotropic voxel size of 1 mm.

###### Anatomical preprocessing

Anatomical T1w and/or T2w images were denoised using the Non-Local Means algorithm (Coupe et al., 2008) as implemented in the DIPY toolbox (Garyfallidis et al., 2014). Bias field correction was applied using the N4 algorithm (Tustison et al., 2010) from the ANTs toolbox (Tustison et al., 2021) using a b-spline knot per voxel of 8 and a shrink factor of 4 for the T1w and a b-spline knot per voxel of 8 and a shrink factor of 4 for the T2w image. Anatomical images were resampled to an isotropic voxel size of 1 mm for the T1w and 1 mm for the T2w image. Brain extraction was performed using SynthStrip (Hoopes et al., 2022); pediatric-tailored weights were used for very young subjects where applicable (Kelley et al., 2024). If both T1w and T2w images were available, they were registered using ANTs (Tustison et al., 2021) using an affine transform.

###### Diffusion Tensor Imaging (DTI)

Diffusion tensor imaging (DTI) models were fitted on the processed volume using the scilpy toolbox (Renauld et al., 2026); fractional anisotropy (FA), axial diffusivity (AD), radial diffusivity (RD), mean diffusivity (MD), mode of anisotropy, and color-coded FA maps were generated. DTI fitting used all available shells under the maximum b-value of 1500 s/mm².

###### Fiber Orientation Distribution Function (fODF)

Fiber orientation distribution functions (fODF) were computed using the scilpy toolbox (Renauld et al., 2026) using the single-shell single-tissue method on the all available shells over the minimum b-value of 700 s/mm². fODF were computed using a maximum spherical harmonic order of 8 in basis descoteaux07. Fiber response functions were estimated based on normative curves of the brain's diffusivities through the developmental age-range as described in Gagnon et al. 2026.

###### Registration to DWI space

Anatomical images were registered to the preprocessed DWI space using ANTs (Tustison et al., 2021). For younger participants (< 2.5 years old), the T2w image, if available, was preferred for registration due to better tissue contrast. If used, the T2w image was registered using non-linear methods using the mean diffusivity map and B0 image as targets. For older participants or if only T1w images were available, the T1w image was registered using non-linear methods with the FA map and B0 image as targets.

###### Tissue segmentation

Tissue segmentation into white matter, grey matter, and cerebrospinal fluid was performed on the anatomical images registered to DWI space. For younger participants (< 2.5 years old), segmentation was performed by registering age-matched templates from the UNC/UMN Baby Connectome Project (Chen et al., 2022). Briefly, templates closest to the participant's age were non-linearly registered to the participant's anatomical images using ANTs (Tustison et al., 2021), and the resulting transforms were applied to the corresponding tissue probability maps. The resulting maps were then thresholded to generate binary masks for each tissue type. For older participants, tissue segmentation was performed using the FAST algorithm from FSL (Zhang et al., 2001; Jenkinson et al., 2012). Similarly to younger participants, resulting probability maps were thresholded to obtain binary masks.

###### Tractography

Whole-brain tractography was performed using the scilpy toolbox (Renauld et al., 2026). Particle Filter Tracking (PFT) was used to leverage anatomical priors from the tissue segmentation to improve streamline generation (Girard et al., 2014). Tracking seeds were randomly placed within the white matter mask with a density of 10 seeds per voxel. Streamlines were propagated using a probabilistic algorithm with a step size of 0.5 mm, a maximum angle between steps of 20°, a minimum length of 20 mm, and a maximum length of 200 mm. Local tracking was performed using a probabilistic algorithm using 10 seeds per voxel. The seeding mask was defined as the white matter mask. Similarly, the tracking mask, in which tracking is allowed, was defined as the white matter mask. Streamlines were propagated with a step size of 0.5 mm, a maximum angle between steps of 20°, a minimum length of 20 mm, and a maximum length of 200 mm. The resulting two tractograms from both methods were then concatenated to form the final whole-brain tractogram.

###### Neurite Orientation Dispersion and Density Imaging (NODDI)

Neurite Orientation Dispersion and Density Imaging (NODDI) models were fitted on the processed DWI volume using the AMICO implementation (Daducci et al., 2015; Zhang et al., 2012); intra-cellular volume fraction (ICVF), orientation dispersion index (ODI), extrac-cellular volume fraction (ECVF), and isotropic volume fraction (ISOVF) maps were generated. The regularization parameters for the NODDI model fitting were: λ_1_ of 0.5 and λ_2_ of 0.001. Diffusivity priors were automatically derived based on the participant's age using normative growth curves as described in Gagnon et al. 2026.

###### Freewater-corrected DTI

Freewater compartment was removed from diffusion tensor imaging (DTI) maps using the freewater model implemented in the AMICO toolbox (Daducci et al., 2015; Pasternak et al., 2009); freewater-corrected diffusion volume, fibervolume, freewater, freewater-corrected fractional anisotropy (FA), axial diffusivity (AD), radial diffusivity (RD), mean diffusivity (MD), mode of anisotropy, and color-coded FA maps were generated. Regularization parameters for model fitting were set as: λ_1_ of 0 and λ_2_ of 0.25. Diffusivity priors were automatically derived based on the participant's age using normative growth curves as described in Gagnon et al. 2026.

###### Bundle segmentation

The closest age-matched white matter atlas (neonates, 3 months, 6 months, 12 months, 24 months or children) was registered into subject-space using an affine transformation. Whole-brain tractograms were segmented using BundleSeg from the scilpy toolbox (St-Onge et al., 2023; Renauld et al., 2026) with a minimal vote ratio of 0.5, an outlier threshold of 0.6, and the euclidean distance. Extracted bundles were then filtered to remove invalid streamlines, single point streamlines, and overlapping points. Then, fixel-based apparent fiber density was computed for each bundle (Raffelt et al., 2017).

###### Tractometry

Atlas' centroids were registered into subject-space using an affine transformation. The centroids were then resampled to 5 points, enabling the derivation of per point metrics. Metric derived per bundle or per point were weighted based on the number of streamline passing through the voxel. This reduces the impact of spurious streamlines on final metric value. For each bundle, multiple metrics were extracted: length, statistic for each endpoint, mean (standard deviation), volume, and streamline count. For each point per bundle (5 points), the following metric were extracted: volume, and mean (standard deviation). Final segmented bundles were colored per point using the jet colormap (affects only the visualisation).

###### Cortical and sub-cortical segmentation

Cortical and subcortical segmentation was performed using recon-all-clinical from Freesurfer (Fischl, 2012; Billot et al., 2023; Iglesias et al., 2023) on the T1w anatomical images. Following segmentation, the Brainnetome Child Atlas (Li et al., 2023) was mapped in subject-space using surface-based registration methods from FreeSurfer (Fischl, 2012) and then converted into voxel labels. For each parcels, volume, surface area, and cortical thickness were measured and outputted in tab-separated value files. For younger participants (< 3 months old), cortical and sub-cortical segmentation was performed using the M-CRIB-S pipeline (Adamson et al., 2020). Younger participants were segmented the Desikan-Killiany (Desikan et al., 2006, Adamson et al., 2020). Following segmentation, volume, surface area, and cortical thickness were measured for each parcel and outputted in tab-separated value files.

###### Connectomics

Structural connectivity matrices were generated using the scilpy toolbox (Renauld et al., 2026) based on the Brainnetome Child Atlas (Li et al., 2023) or the Desikan-Killiany atlas (Desikan et al., 2006) depending on the participant's age. For each participant, labels in anatomical space were first registered in diffusion space using the already computed transformations with a nearest neighbor interpolation method. Then, the final tractogram was decomposed into individual connections by extracting each streamline connecting a pair of parcels. Streamlines shorter than 20 mm or longer than 200 mm were discarded. Loops were removed. Hierarchical QuickBundles was used to remove outliers using a threshold of 0.6. Curvature-based filtering was applied to remove streamlines with sharp curves using a maximum angle of 330.0° over 10.0 mm. To mitigate the risk of false-positive connections, COMMIT (Daducci et al., 2015) was applied to the tractogram using the stick, zeppelin, and ball model to optimize the fit between the tractogram and the diffusion data. Diffusivity parameters for COMMIT were set based on age-specific normative values as described in Gagnon et al. 2026. Using COMMIT2 (Schiavi et al., 2020) with a clustering prior strength of 0.001, the contribution of each streamline to the diffusion signal was evaluated and streamlines with zero contribution were removed from the tractogram to further reduce false-positive connections. To obtain the fODF amplitude specific to each connection, fixel-based apparent fiber density was computed for each extracted connection (Raffelt et al., 2017). Finally, structural connectivity matrices were generated by computing, for each pair of parcels, the number of streamlines, the mean streamline length, and the mean FA, AD, RD, MD, total apparent fiber density, number of fiber orientation, and fixel-based apparent fiber density.
