## Supplementary for "sf-pediatric: A robust and age-adaptable end-to-end pipeline for pediatric diffusion MRI": SupplementaryFile2.html

sf-pediatric MultiQC Global Report: MultiQC Report


Loading report..

v1.32


Theme

- Light
- Dark
- Auto

### sf-pediatric MultiQC Global Report

Highlight

 Rename

 Show / Hide


AI Analysis

 Export

 Settings

 Citations

 About

- General Stats
- Framewise Displacement
- Streamline Count
- Coverage
- Tractometry
  - Fractional Anisotropy (FA)
  - Bundle Volume
  - Streamline Count
- Cortical Regions
  - Left Hemisphere Volume Distribution
  - Right Hemisphere Volume Distribution
- Subcortical Regions
- scilus/sf-pediatric Workflow Summary
- Software Versions
- sf-pediatric Methods Description

Toolbox

###### MultiQC Toolbox

###### Apply Highlight Samples


Regex mode

regex help

 Clear all

###### Rename Samples Bulk input Apply


Paste two columns of a tab-delimited table here (eg. from Excel). First column should be the old name, second
column the new name.

Add

Regex mode

regex help

 Clear all

###### Apply Show / Hide Samples

Hide matching samples

Show only matching samples

Regex mode

regex help

 Clear all

###### Explain with AI

Configure AI settings to get explanations of plots and data in this report.

AI Provider

Endpoint


Use the OpenAI API-style requests with a custom endpoint.

Model

API Key


Keys entered here will be stored in your browser's local storage. See
the docs.

Additional Payload


Any additional options passed in API request payload. Enter as a JSON object.

Context Window


The maximum number of tokens that can be processed in a single request

Anonymize samples

Switch out sample names with random identifiers

###### Export Plots

- Images
- Data

Width

px

Height

px

Maintain aspect ratio on resize

Plot format

PNG
SVG

Plot scaling

X

File format:

Tab-separated
Comma-separated
JSON

Note: Additional data was saved in
`multiqc_data` when this report was generated.

###### Choose Plots All None

   Download Plot Images

If you use plots from MultiQC in a publication or presentation, please cite:

> **MultiQC: Summarize analysis results for multiple tools and samples in a single report**  
> *Philip Ewels, Måns Magnusson, Sverker Lundin and Max Käller*  
> Bioinformatics (2016)  
> doi:
> 10.1093/bioinformatics/btw354  
> PMID: 27312411

###### Save Settings

Report settings are automatically saved in your browser as you use the
toolbox. You can also save *named* configurations below.

Save to Browser

 Save to File

###### Load Settings

Choose a saved report profile from the browser or load from a file:

[ select named settings from browser ]

Load

 Delete

Set as default for all reports

 Clear default

Load from file

###### Tool Citations

Please remember to cite *all of the tools* that you use in your analysis.

List of DOIs

 BibTeX file

###### About MultiQC

This report was generated using MultiQC, version 1.32

Video: Using MultiQC Reports

 MultiQC homepage

 MultiQC documentation

 Source code

 Issue tracker

MultiQC is published in Bioinformatics:

> **MultiQC: Summarize analysis results for multiple tools and samples in a single report**  
> *Philip Ewels, Måns Magnusson, Sverker Lundin and Max Käller*  
> Bioinformatics (2016)  
> doi:
> 10.1093/bioinformatics/btw354  
> PMID: 27312411

MultiQC is developed by Seqera.

Scroll to top

### sf-pediatric MultiQC Global Report

A modular tool to aggregate results from bioinformatics analyses across many samples into a single report.

> This report has been generated by the sf-pediatric analysis pipeline.

###### JavaScript Disabled

MultiQC reports use JavaScript for plots and toolbox functions. It looks like you have JavaScript disabled in your
web browser. Please note that many of the report functions will not work as intended.

Loading report..

Report
generated on 2025-12-19, 17:20 EST
based on data in:
`/tmp/nxf.UUSuzs9Ta3`

Summarize report

Copy report prompt

**Welcome!** Not sure where to start?
Watch a tutorial video
*(6:06)*

don't show again

###### Report AI Summary

More details…

Provider: , model:

Chat with Seqera AI

#### General Statistics

###### AI Summary

Provider: , model:

Chat with Seqera AI

Table
 Export...

Copy prompt


Summarize plot

Created with MultiQC

Copy table

 Configure columns

 Sort by highlight

 Scatter plot

 Violin plot
Export as CSV...
Showing ^10^/_10_ rows and ^6^/_6_ columns.

Copy Prompt


Summarize table

| Sample Name | Max FD | SC | Dice Coef. | % Bundles | % Outliers | % Outliers |
| --- | --- | --- | --- | --- | --- | --- |
| sub-01\_ses-baseline | 0.25mm | 393996 | 0.880 | 100.0% | 0.5% | 0.0% |
| sub-02\_ses-baseline | 0.18mm | 574856 | 0.898 | 100.0% | 0.5% | 0.0% |
| sub-03\_ses-baseline | 0.86mm | 512868 | 0.883 | 98.0% | 0.5% | 0.0% |
| sub-04\_ses-baseline | 1.84mm | 418947 | 0.880 | 100.0% | 0.0% | 0.0% |
| sub-05\_ses-baseline | 0.17mm | 512012 | 0.894 | 100.0% | 0.5% | 0.0% |
| sub-06\_ses-baseline | 3.09mm | 495825 | 0.899 | 100.0% | 0.0% | 0.0% |
| sub-07\_ses-baseline | 1.12mm | 503293 | 0.890 | 98.0% | 2.1% | 0.0% |
| sub-08\_ses-baseline | 1.58mm | 552596 | 0.889 | 100.0% | 0.5% | 0.0% |
| sub-09\_ses-baseline | 3.36mm | 506468 | 0.900 | 100.0% | 0.5% | 0.0% |
| sub-11\_ses-baseline | 1.27mm | 535355 | 0.908 | 100.0% | 0.5% | 0.0% |

###### General Statistics: Columns

Uncheck the tick box to hide columns. Click and drag the handle on the left to change order. Table ID: `general_stats_table_table`

Show All
Show None

| Sort | Visible | Group | Column | Description | ID | Scale |
| --- | --- | --- | --- | --- | --- | --- |
| \|\| |  | Framewise Displacement | Max FD | Maximum framewise displacement | `framewise_displacement-max_fd` |  |
| \|\| |  | Streamline Count | SC | Streamline count | `streamline_count-streamline_count` |  |
| \|\| |  | Coverage | Dice Coef. | Dice coefficient for coverage | `coverage-dice_coefficient` |  |
| \|\| |  | Tractometry | % Bundles | Percentage of bundles detected | `tractometry-bundle_percentage` |  |
| \|\| |  | Cortical Regions | % Outliers | Percentage of cortical regions with volumes outside 3\*IQR range | `cortical_regions-region_pct` |  |
| \|\| |  | Subcortical Regions | % Outliers | Percentage of subcortical regions with volumes outside 3\*IQR range | `subcortical_regions-region_pct` |  |

Close

#### Framewise Displacement

Assessment of subject motion during acquisition using the framewise displacement (FD) metric calculated by FSL's `eddy` tool for quality control.https://github.com/nf-neuro/MultiQC\_neuroimaging

##### Framewise Displacement 3 5 2

Framewise displacement (FD) across volumes for all subjects. Each line represents a subject and its relative movement over time. For each volume, the FDrepresents the amount of movement from the previous volume. Subjects with big spikes in FD may indicate excessive motion during scanning. While `eddy` attempts to correct for motion, users should be cautious when interpreting data from subjects with high FD values. Lines are colored based on maximum FD. Green: <0.8mm (pass), Yellow: 0.8-2.0mm (warn), Red: >2.0mm (fail)

###### AI Summary

Provider: , model:

Chat with Seqera AI

Export...

Copy prompt


Summarize plot

Created with MultiQC

#### Streamline Count

Visualization of the number of streamlines in a filtered/unfiltered tractogram using IQR-based outlier detection for quality control.https://github.com/nf-neuro/MultiQC\_neuroimaging

##### Streamline Count Quality 9 1

Tractogram streamline count quality control with outliers detected using the IQR method.
Using an acceptable range defined as IQR \* 3, subjects with streamline counts
falling outside this range will be flagged, and might indicate potential issues with tractography,
tissue segmentation, or fODF reconstruction. Often times, extremely high streamline counts can be
attributed to a lot of small streamlines generated from noisy fODF peaks. Extremely low streamline counts
may indicate poor white matter segmentation or insufficient seeding. While this might not be sufficient to
exclude a subject, users should investigate such outliers further to ensure data quality.
Pass: within Q1 - 3*IQR to Q3 + 3*IQR range
[401568 - 625857], Fail: outside range

###### AI Summary

Provider: , model:

Chat with Seqera AI

Table
 Export...

Copy prompt


Summarize plot

Created with MultiQC

Copy table

 Sort by highlight

 Violin plot
Export as CSV...
Showing ^10^/_10_ rows.

Copy Prompt


Summarize table

| Sample Name | Streamline Count |
| --- | --- |
| sub-01\_ses-baseline | 393996 |
| sub-02\_ses-baseline | 574856 |
| sub-03\_ses-baseline | 512868 |
| sub-04\_ses-baseline | 418947 |
| sub-05\_ses-baseline | 512012 |
| sub-06\_ses-baseline | 495825 |
| sub-07\_ses-baseline | 503293 |
| sub-08\_ses-baseline | 552596 |
| sub-09\_ses-baseline | 506468 |
| sub-11\_ses-baseline | 535355 |

###### Streamline Count: Columns

Uncheck the tick box to hide columns. Click and drag the handle on the left to change order. Table ID: `streamline_count_violin_table`

Show All
Show None

| Sort | Visible | Group | Column | Description | ID | Scale |
| --- | --- | --- | --- | --- | --- | --- |
| \|\| |  |  | Streamline Count | Total number of streamlines | `Streamline_Count` |  |

Close

#### Coverage

Assessment of the white matter coverage of tractograms using the DICE coefficient between the tract density map and white matter mask for quality control.https://github.com/nf-neuro/MultiQC\_neuroimaging

##### Coverage Quality 2 8

White matter coverage by the tractogram measured by computing the DICE coefficient
between the subject's tract density map and its corresponding white matter mask.
Higher DICE coefficients indicate better coverage of the white matter. Users should expect
values over 0.9 for unfiltered tractograms and over 0.8 for filtered tractograms.
Pass: >0.9, Warn: 0.8-0.9, Fail: <0.8

###### AI Summary

Provider: , model:

Chat with Seqera AI

Table
 Export...

Copy prompt


Summarize plot

Created with MultiQC

Copy table

 Sort by highlight

 Violin plot
Export as CSV...
Showing ^10^/_10_ rows.

Copy Prompt


Summarize table

| Sample Name | Dice Coefficient |
| --- | --- |
| sub-01\_ses-baseline | 0.880 |
| sub-02\_ses-baseline | 0.898 |
| sub-03\_ses-baseline | 0.883 |
| sub-04\_ses-baseline | 0.880 |
| sub-05\_ses-baseline | 0.894 |
| sub-06\_ses-baseline | 0.899 |
| sub-07\_ses-baseline | 0.890 |
| sub-08\_ses-baseline | 0.889 |
| sub-09\_ses-baseline | 0.900 |
| sub-11\_ses-baseline | 0.908 |

###### Coverage: Dice Coefficients: Columns

Uncheck the tick box to hide columns. Click and drag the handle on the left to change order. Table ID: `coverage_dice_violin_table`

Show All
Show None

| Sort | Visible | Group | Column | Description | ID | Scale |
| --- | --- | --- | --- | --- | --- | --- |
| \|\| |  |  | Dice Coefficient | Dice coefficient for coverage | `Dice` |  |

Close

#### Tractometry

This section contains QC metrics from tractometry analysis. For QC purposes, they only include fractional anisotropy (FA), volume, and streamline for the whole bundles. Additional metrics can be found in the statistics table exported by the pipeline. The status bars indicate flagged subjects based on the percentage of bundles detected, with thresholds configurable in the MultiQC configuration file.https://github.com/scilus/nf-pediatric

##### Fractional Anisotropy (FA) 10

Distribution of FA values per bundle. You should look for extreme outliers, as it can be indicative of issues in bundle extraction.

###### AI Summary

Provider: , model:

Chat with Seqera AI

Table
 Export...

Copy prompt


Summarize plot

Created with MultiQC

Copy table

 Configure columns

 Sort by highlight

 Scatter plot

 Violin plot
Export as CSV...
Showing ^10^/_10_ rows and ^50^/_50_ columns.

Copy Prompt


Summarize table

| Sample Name | AC | AF\_L | AF\_R | CC\_Fr\_1 | CC\_Fr\_2 | CC\_Oc | CC\_Pa | CC\_Pr\_Po | CC\_Te | CG\_L | CG\_L\_An | CG\_L\_Po | CG\_L\_curve | CG\_R | CG\_R\_An | CG\_R\_Po | CG\_R\_curve | FAT\_L | FAT\_R | FPT\_L | FPT\_L\_Brainstem | FPT\_R | FPT\_R\_Brainstem | FX\_L | FX\_R | ICP\_L | ICP\_R | IFOF\_L | IFOF\_R | ILF\_L | ILF\_R | MCP | MdLF\_L | MdLF\_R | OR\_ML\_L | OR\_ML\_R | POPT\_L | POPT\_L\_Brainstem | POPT\_R | POPT\_R\_Brainstem | PYT\_L | PYT\_L\_Brainstem | PYT\_R | PYT\_R\_Brainstem | SCP\_L | SCP\_R | SLF\_L | SLF\_R | UF\_L | UF\_R |
| --- | --- | --- | --- | --- | --- | --- | --- | --- | --- | --- | --- | --- | --- | --- | --- | --- | --- | --- | --- | --- | --- | --- | --- | --- | --- | --- | --- | --- | --- | --- | --- | --- | --- | --- | --- | --- | --- | --- | --- | --- | --- | --- | --- | --- | --- | --- | --- | --- | --- | --- |
| sub-01\_ses-baseline | 0.4 | 0.4 | 0.4 | 0.5 | 0.5 | 0.6 | 0.5 | 0.5 | 0.5 | 0.5 | 0.5 | 0.4 | 0.5 | 0.5 | 0.5 | 0.4 | 0.5 | 0.4 | 0.4 | 0.5 | 0.5 | 0.5 | 0.5 | 0.4 | 0.3 | 0.4 | 0.5 | 0.5 | 0.5 | 0.5 | 0.5 | 0.6 | 0.4 | 0.4 | 0.5 | 0.5 | 0.5 | 0.5 | 0.5 | 0.5 | 0.5 | 0.5 | 0.5 | 0.5 | 0.5 | 0.5 | 0.4 | 0.4 | 0.4 | 0.4 |
| sub-02\_ses-baseline | 0.4 | 0.4 | 0.4 | 0.5 | 0.5 | 0.6 | 0.5 | 0.5 | 0.6 | 0.5 | 0.5 | 0.4 | 0.5 | 0.5 | 0.5 | 0.4 | 0.4 | 0.4 | 0.4 | 0.5 | 0.5 | 0.5 | 0.5 | 0.5 | 0.4 | 0.4 | 0.4 | 0.5 | 0.5 | 0.5 | 0.5 | 0.5 | 0.4 | 0.4 | 0.5 | 0.5 | 0.5 | 0.5 | 0.5 | 0.5 | 0.5 | 0.5 | 0.5 | 0.5 | 0.4 | 0.4 | 0.4 | 0.4 | 0.4 | 0.4 |
| sub-03\_ses-baseline | 0.4 | 0.4 | 0.4 | 0.5 | 0.5 | 0.6 | 0.5 | 0.5 | 0.6 | 0.4 | 0.4 | 0.4 | 0.4 | 0.4 | 0.4 | 0.4 | 0.4 | 0.4 | 0.4 | 0.5 | 0.5 | 0.5 | 0.5 | 0.5 |  | 0.4 | 0.4 | 0.5 | 0.5 | 0.5 | 0.5 | 0.5 | 0.4 | 0.4 | 0.5 | 0.5 | 0.5 | 0.5 | 0.5 | 0.5 | 0.5 | 0.5 | 0.5 | 0.5 | 0.4 | 0.4 | 0.4 | 0.4 | 0.4 | 0.4 |
| sub-04\_ses-baseline | 0.4 | 0.4 | 0.4 | 0.5 | 0.5 | 0.6 | 0.6 | 0.5 | 0.6 | 0.5 | 0.5 | 0.4 | 0.5 | 0.5 | 0.5 | 0.4 | 0.5 | 0.4 | 0.4 | 0.5 | 0.5 | 0.5 | 0.4 | 0.4 | 0.4 | 0.4 | 0.4 | 0.5 | 0.5 | 0.5 | 0.5 | 0.5 | 0.5 | 0.4 | 0.5 | 0.5 | 0.5 | 0.5 | 0.5 | 0.5 | 0.5 | 0.5 | 0.5 | 0.5 | 0.5 | 0.4 | 0.4 | 0.4 | 0.4 | 0.4 |
| sub-05\_ses-baseline | 0.3 | 0.4 | 0.4 | 0.5 | 0.5 | 0.6 | 0.5 | 0.5 | 0.5 | 0.5 | 0.5 | 0.4 | 0.4 | 0.4 | 0.4 | 0.4 | 0.4 | 0.4 | 0.4 | 0.5 | 0.4 | 0.5 | 0.5 | 0.3 | 0.3 | 0.4 | 0.4 | 0.4 | 0.4 | 0.4 | 0.4 | 0.5 | 0.4 | 0.4 | 0.5 | 0.5 | 0.5 | 0.4 | 0.5 | 0.5 | 0.5 | 0.5 | 0.5 | 0.5 | 0.4 | 0.4 | 0.4 | 0.4 | 0.4 | 0.4 |
| sub-06\_ses-baseline | 0.3 | 0.4 | 0.4 | 0.4 | 0.5 | 0.6 | 0.5 | 0.5 | 0.6 | 0.4 | 0.4 | 0.4 | 0.4 | 0.4 | 0.4 | 0.4 | 0.4 | 0.4 | 0.4 | 0.4 | 0.4 | 0.4 | 0.4 | 0.3 | 0.3 | 0.4 | 0.4 | 0.4 | 0.4 | 0.5 | 0.4 | 0.5 | 0.4 | 0.4 | 0.5 | 0.5 | 0.5 | 0.5 | 0.5 | 0.5 | 0.5 | 0.5 | 0.5 | 0.5 | 0.4 | 0.4 | 0.3 | 0.4 | 0.4 | 0.4 |
| sub-07\_ses-baseline | 0.3 | 0.5 | 0.5 | 0.6 | 0.6 | 0.6 | 0.6 | 0.6 | 0.6 | 0.5 | 0.5 | 0.4 | 0.5 | 0.5 | 0.5 | 0.4 |  | 0.5 | 0.5 | 0.5 | 0.5 | 0.5 | 0.5 | 0.4 | 0.4 | 0.4 | 0.4 | 0.5 | 0.5 | 0.5 | 0.5 | 0.4 | 0.4 | 0.5 | 0.5 | 0.5 | 0.5 | 0.5 | 0.5 | 0.4 | 0.5 | 0.5 | 0.5 | 0.5 | 0.4 | 0.4 | 0.5 | 0.5 | 0.4 | 0.4 |
| sub-08\_ses-baseline | 0.4 | 0.4 | 0.4 | 0.5 | 0.5 | 0.6 | 0.5 | 0.5 | 0.5 | 0.5 | 0.5 | 0.3 | 0.4 | 0.5 | 0.5 | 0.3 | 0.4 | 0.4 | 0.4 | 0.5 | 0.4 | 0.5 | 0.4 | 0.4 | 0.3 | 0.4 | 0.4 | 0.5 | 0.5 | 0.4 | 0.5 | 0.5 | 0.4 | 0.4 | 0.4 | 0.5 | 0.5 | 0.4 | 0.4 | 0.4 | 0.5 | 0.4 | 0.5 | 0.4 | 0.4 | 0.4 | 0.4 | 0.4 | 0.4 | 0.4 |
| sub-09\_ses-baseline | 0.3 | 0.4 | 0.4 | 0.5 | 0.5 | 0.6 | 0.5 | 0.5 | 0.5 | 0.4 | 0.4 | 0.3 | 0.4 | 0.4 | 0.4 | 0.3 | 0.4 | 0.4 | 0.4 | 0.4 | 0.4 | 0.5 | 0.5 | 0.4 | 0.4 | 0.4 | 0.4 | 0.4 | 0.4 | 0.5 | 0.5 | 0.4 | 0.4 | 0.4 | 0.5 | 0.5 | 0.5 | 0.5 | 0.5 | 0.5 | 0.5 | 0.4 | 0.5 | 0.5 | 0.4 | 0.4 | 0.4 | 0.4 | 0.4 | 0.4 |
| sub-11\_ses-baseline | 0.4 | 0.4 | 0.4 | 0.5 | 0.5 | 0.6 | 0.5 | 0.5 | 0.6 | 0.5 | 0.5 | 0.4 | 0.5 | 0.5 | 0.5 | 0.4 | 0.4 | 0.4 | 0.4 | 0.5 | 0.4 | 0.5 | 0.4 | 0.4 | 0.4 | 0.5 | 0.4 | 0.5 | 0.5 | 0.5 | 0.5 | 0.5 | 0.4 | 0.4 | 0.5 | 0.5 | 0.5 | 0.5 | 0.5 | 0.5 | 0.5 | 0.5 | 0.5 | 0.5 | 0.4 | 0.4 | 0.4 | 0.4 | 0.4 | 0.4 |

###### Tractometry: Fractional Anisotropy (FA): Columns

Uncheck the tick box to hide columns. Click and drag the handle on the left to change order. Table ID: `tractometry_fa_violin_table`

Show All
Show None

| Sort | Visible | Group | Column | Description | ID | Scale |
| --- | --- | --- | --- | --- | --- | --- |
| \|\| |  |  | AC | Fractional Anisotropy (FA) for AC | `AC` |  |
| \|\| |  |  | AF\_L | Fractional Anisotropy (FA) for AF\_L | `AF_L` |  |
| \|\| |  |  | AF\_R | Fractional Anisotropy (FA) for AF\_R | `AF_R` |  |
| \|\| |  |  | CC\_Fr\_1 | Fractional Anisotropy (FA) for CC\_Fr\_1 | `CC_Fr_1` |  |
| \|\| |  |  | CC\_Fr\_2 | Fractional Anisotropy (FA) for CC\_Fr\_2 | `CC_Fr_2` |  |
| \|\| |  |  | CC\_Oc | Fractional Anisotropy (FA) for CC\_Oc | `CC_Oc` |  |
| \|\| |  |  | CC\_Pa | Fractional Anisotropy (FA) for CC\_Pa | `CC_Pa` |  |
| \|\| |  |  | CC\_Pr\_Po | Fractional Anisotropy (FA) for CC\_Pr\_Po | `CC_Pr_Po` |  |
| \|\| |  |  | CC\_Te | Fractional Anisotropy (FA) for CC\_Te | `CC_Te` |  |
| \|\| |  |  | CG\_L | Fractional Anisotropy (FA) for CG\_L | `CG_L` |  |
| \|\| |  |  | CG\_L\_An | Fractional Anisotropy (FA) for CG\_L\_An | `CG_L_An` |  |
| \|\| |  |  | CG\_L\_Po | Fractional Anisotropy (FA) for CG\_L\_Po | `CG_L_Po` |  |
| \|\| |  |  | CG\_L\_curve | Fractional Anisotropy (FA) for CG\_L\_curve | `CG_L_curve` |  |
| \|\| |  |  | CG\_R | Fractional Anisotropy (FA) for CG\_R | `CG_R` |  |
| \|\| |  |  | CG\_R\_An | Fractional Anisotropy (FA) for CG\_R\_An | `CG_R_An` |  |
| \|\| |  |  | CG\_R\_Po | Fractional Anisotropy (FA) for CG\_R\_Po | `CG_R_Po` |  |
| \|\| |  |  | CG\_R\_curve | Fractional Anisotropy (FA) for CG\_R\_curve | `CG_R_curve` |  |
| \|\| |  |  | FAT\_L | Fractional Anisotropy (FA) for FAT\_L | `FAT_L` |  |
| \|\| |  |  | FAT\_R | Fractional Anisotropy (FA) for FAT\_R | `FAT_R` |  |
| \|\| |  |  | FPT\_L | Fractional Anisotropy (FA) for FPT\_L | `FPT_L` |  |
| \|\| |  |  | FPT\_L\_Brainstem | Fractional Anisotropy (FA) for FPT\_L\_Brainstem | `FPT_L_Brainstem` |  |
| \|\| |  |  | FPT\_R | Fractional Anisotropy (FA) for FPT\_R | `FPT_R` |  |
| \|\| |  |  | FPT\_R\_Brainstem | Fractional Anisotropy (FA) for FPT\_R\_Brainstem | `FPT_R_Brainstem` |  |
| \|\| |  |  | FX\_L | Fractional Anisotropy (FA) for FX\_L | `FX_L` |  |
| \|\| |  |  | FX\_R | Fractional Anisotropy (FA) for FX\_R | `FX_R` |  |
| \|\| |  |  | ICP\_L | Fractional Anisotropy (FA) for ICP\_L | `ICP_L` |  |
| \|\| |  |  | ICP\_R | Fractional Anisotropy (FA) for ICP\_R | `ICP_R` |  |
| \|\| |  |  | IFOF\_L | Fractional Anisotropy (FA) for IFOF\_L | `IFOF_L` |  |
| \|\| |  |  | IFOF\_R | Fractional Anisotropy (FA) for IFOF\_R | `IFOF_R` |  |
| \|\| |  |  | ILF\_L | Fractional Anisotropy (FA) for ILF\_L | `ILF_L` |  |
| \|\| |  |  | ILF\_R | Fractional Anisotropy (FA) for ILF\_R | `ILF_R` |  |
| \|\| |  |  | MCP | Fractional Anisotropy (FA) for MCP | `MCP` |  |
| \|\| |  |  | MdLF\_L | Fractional Anisotropy (FA) for MdLF\_L | `MdLF_L` |  |
| \|\| |  |  | MdLF\_R | Fractional Anisotropy (FA) for MdLF\_R | `MdLF_R` |  |
| \|\| |  |  | OR\_ML\_L | Fractional Anisotropy (FA) for OR\_ML\_L | `OR_ML_L` |  |
| \|\| |  |  | OR\_ML\_R | Fractional Anisotropy (FA) for OR\_ML\_R | `OR_ML_R` |  |
| \|\| |  |  | POPT\_L | Fractional Anisotropy (FA) for POPT\_L | `POPT_L` |  |
| \|\| |  |  | POPT\_L\_Brainstem | Fractional Anisotropy (FA) for POPT\_L\_Brainstem | `POPT_L_Brainstem` |  |
| \|\| |  |  | POPT\_R | Fractional Anisotropy (FA) for POPT\_R | `POPT_R` |  |
| \|\| |  |  | POPT\_R\_Brainstem | Fractional Anisotropy (FA) for POPT\_R\_Brainstem | `POPT_R_Brainstem` |  |
| \|\| |  |  | PYT\_L | Fractional Anisotropy (FA) for PYT\_L | `PYT_L` |  |
| \|\| |  |  | PYT\_L\_Brainstem | Fractional Anisotropy (FA) for PYT\_L\_Brainstem | `PYT_L_Brainstem` |  |
| \|\| |  |  | PYT\_R | Fractional Anisotropy (FA) for PYT\_R | `PYT_R` |  |
| \|\| |  |  | PYT\_R\_Brainstem | Fractional Anisotropy (FA) for PYT\_R\_Brainstem | `PYT_R_Brainstem` |  |
| \|\| |  |  | SCP\_L | Fractional Anisotropy (FA) for SCP\_L | `SCP_L` |  |
| \|\| |  |  | SCP\_R | Fractional Anisotropy (FA) for SCP\_R | `SCP_R` |  |
| \|\| |  |  | SLF\_L | Fractional Anisotropy (FA) for SLF\_L | `SLF_L` |  |
| \|\| |  |  | SLF\_R | Fractional Anisotropy (FA) for SLF\_R | `SLF_R` |  |
| \|\| |  |  | UF\_L | Fractional Anisotropy (FA) for UF\_L | `UF_L` |  |
| \|\| |  |  | UF\_R | Fractional Anisotropy (FA) for UF\_R | `UF_R` |  |

Close

##### Bundle Volume 10

Distribution of volume per bundle. You should look for extreme outliers, as it can indicate issues in bundle extraction. Bundles with very low volume may be incomplete while bundles with high volume might include spurious streamlines or non-desirable structures.

###### AI Summary

Provider: , model:

Chat with Seqera AI

Table
 Export...

Copy prompt


Summarize plot

Created with MultiQC

Copy table

 Configure columns

 Sort by highlight

 Scatter plot

 Violin plot
Export as CSV...
Showing ^10^/_10_ rows and ^50^/_50_ columns.

Copy Prompt


Summarize table

| Sample Name | AC | AF\_L | AF\_R | CC\_Fr\_1 | CC\_Fr\_2 | CC\_Oc | CC\_Pa | CC\_Pr\_Po | CC\_Te | CG\_L | CG\_L\_An | CG\_L\_Po | CG\_L\_curve | CG\_R | CG\_R\_An | CG\_R\_Po | CG\_R\_curve | FAT\_L | FAT\_R | FPT\_L | FPT\_L\_Brainstem | FPT\_R | FPT\_R\_Brainstem | FX\_L | FX\_R | ICP\_L | ICP\_R | IFOF\_L | IFOF\_R | ILF\_L | ILF\_R | MCP | MdLF\_L | MdLF\_R | OR\_ML\_L | OR\_ML\_R | POPT\_L | POPT\_L\_Brainstem | POPT\_R | POPT\_R\_Brainstem | PYT\_L | PYT\_L\_Brainstem | PYT\_R | PYT\_R\_Brainstem | SCP\_L | SCP\_R | SLF\_L | SLF\_R | UF\_L | UF\_R |
| --- | --- | --- | --- | --- | --- | --- | --- | --- | --- | --- | --- | --- | --- | --- | --- | --- | --- | --- | --- | --- | --- | --- | --- | --- | --- | --- | --- | --- | --- | --- | --- | --- | --- | --- | --- | --- | --- | --- | --- | --- | --- | --- | --- | --- | --- | --- | --- | --- | --- | --- |
| sub-01\_ses-baseline | 5090.0 | 40022.0 | 39859.0 | 53246.0 | 93379.0 | 51263.0 | 63164.0 | 83937.0 | 1847.0 | 10464.0 | 8174.0 | 1542.0 | 2686.0 | 11078.0 | 9569.0 | 1531.0 | 2650.0 | 37264.0 | 32083.0 | 58081.0 | 55022.0 | 58734.0 | 60912.0 | 2242.0 | 883.0 | 6384.0 | 9726.0 | 53601.0 | 56971.0 | 44834.0 | 40797.0 | 23123.0 | 20668.0 | 27812.0 | 18247.0 | 13931.0 | 58570.0 | 49942.0 | 61475.0 | 55003.0 | 55629.0 | 49569.0 | 59410.0 | 60058.0 | 11012.0 | 13412.0 | 38570.0 | 50787.0 | 23472.0 | 11189.0 |
| sub-02\_ses-baseline | 946.0 | 44934.0 | 49578.0 | 62660.0 | 90724.0 | 55768.0 | 54429.0 | 69210.0 | 2480.0 | 20647.0 | 16390.0 | 2356.0 | 2010.0 | 16711.0 | 11084.0 | 2210.0 | 2165.0 | 38570.0 | 39074.0 | 73426.0 | 55655.0 | 71530.0 | 70166.0 | 557.0 | 913.0 | 11577.0 | 2992.0 | 66832.0 | 57243.0 | 50142.0 | 46516.0 | 46240.0 | 30576.0 | 33010.0 | 23700.0 | 20115.0 | 65176.0 | 50546.0 | 57940.0 | 52007.0 | 61176.0 | 55223.0 | 61144.0 | 59351.0 | 16908.0 | 17970.0 | 58534.0 | 81003.0 | 21004.0 | 18880.0 |
| sub-03\_ses-baseline | 3425.0 | 37944.0 | 25460.0 | 55498.0 | 81842.0 | 46808.0 | 84114.0 | 79182.0 | 11204.0 | 2759.0 | 3396.0 | 2580.0 | 1115.0 | 14158.0 | 12697.0 | 2391.0 | 1930.0 | 38184.0 | 35449.0 | 65678.0 | 55777.0 | 75770.0 | 70383.0 | 1728.0 |  | 13966.0 | 7389.0 | 53451.0 | 56390.0 | 47083.0 | 39331.0 | 41884.0 | 16815.0 | 23646.0 | 18701.0 | 22489.0 | 57084.0 | 32293.0 | 52249.0 | 36085.0 | 57025.0 | 46234.0 | 58276.0 | 52284.0 | 25180.0 | 13786.0 | 41990.0 | 45064.0 | 6243.0 | 17799.0 |
| sub-04\_ses-baseline | 2604.0 | 40457.0 | 51287.0 | 50578.0 | 91353.0 | 44478.0 | 49018.0 | 65263.0 | 16023.0 | 9705.0 | 6679.0 | 2376.0 | 1631.0 | 21879.0 | 16672.0 | 3584.0 | 2825.0 | 40093.0 | 37800.0 | 77282.0 | 53219.0 | 69909.0 | 63854.0 | 831.0 | 3866.0 | 9881.0 | 9525.0 | 25748.0 | 53388.0 | 50749.0 | 47345.0 | 22351.0 | 29047.0 | 24351.0 | 10899.0 | 13967.0 | 60372.0 | 47000.0 | 56804.0 | 41818.0 | 61161.0 | 55109.0 | 62020.0 | 56041.0 | 16858.0 | 17789.0 | 50540.0 | 42912.0 | 20408.0 | 19238.0 |
| sub-05\_ses-baseline | 165.0 | 56342.0 | 65135.0 | 49097.0 | 79646.0 | 50902.0 | 66143.0 | 53455.0 | 709.0 | 7585.0 | 8129.0 | 973.0 | 1900.0 | 20647.0 | 22433.0 | 1114.0 | 767.0 | 39326.0 | 45373.0 | 78758.0 | 55343.0 | 81827.0 | 74973.0 | 7153.0 | 3849.0 | 3842.0 | 6535.0 | 47783.0 | 40881.0 | 41596.0 | 49537.0 | 38918.0 | 13276.0 | 21765.0 | 18864.0 | 22060.0 | 64792.0 | 49515.0 | 67484.0 | 57103.0 | 67210.0 | 51184.0 | 59638.0 | 56760.0 | 22245.0 | 16842.0 | 60952.0 | 65591.0 | 18386.0 | 17575.0 |
| sub-06\_ses-baseline | 1036.0 | 64740.0 | 33674.0 | 60108.0 | 87288.0 | 38230.0 | 59470.0 | 60927.0 | 4099.0 | 10170.0 | 8151.0 | 2697.0 | 1969.0 | 9736.0 | 8553.0 | 2127.0 | 2405.0 | 40617.0 | 45664.0 | 66726.0 | 73266.0 | 76380.0 | 76381.0 | 4798.0 | 3480.0 | 8128.0 | 7310.0 | 47722.0 | 42485.0 | 49560.0 | 37299.0 | 40564.0 | 12887.0 | 17539.0 | 20771.0 | 17581.0 | 67213.0 | 63041.0 | 56162.0 | 51907.0 | 67820.0 | 68395.0 | 65121.0 | 60575.0 | 14677.0 | 16678.0 | 51732.0 | 59486.0 | 25416.0 | 15417.0 |
| sub-07\_ses-baseline | 412.0 | 47862.0 | 51649.0 | 55831.0 | 95601.0 | 51583.0 | 61622.0 | 47209.0 | 1047.0 | 12390.0 | 9978.0 | 1362.0 | 2069.0 | 20406.0 | 19370.0 | 1309.0 |  | 40757.0 | 42919.0 | 80443.0 | 84741.0 | 82248.0 | 81093.0 | 13300.0 | 6812.0 | 11570.0 | 7477.0 | 73620.0 | 74272.0 | 42744.0 | 45494.0 | 55073.0 | 12971.0 | 21098.0 | 22940.0 | 16030.0 | 54911.0 | 50258.0 | 50841.0 | 42363.0 | 72939.0 | 68085.0 | 68705.0 | 62618.0 | 36165.0 | 22151.0 | 47827.0 | 65552.0 | 27592.0 | 22730.0 |
| sub-08\_ses-baseline | 2110.0 | 52949.0 | 47765.0 | 54732.0 | 89616.0 | 53099.0 | 59832.0 | 50625.0 | 12387.0 | 13116.0 | 10539.0 | 2774.0 | 3061.0 | 12134.0 | 12684.0 | 1643.0 | 2279.0 | 41813.0 | 36793.0 | 75121.0 | 68885.0 | 74323.0 | 67403.0 | 7286.0 | 6248.0 | 3174.0 | 8532.0 | 66692.0 | 66201.0 | 48594.0 | 47698.0 | 32233.0 | 29389.0 | 27424.0 | 23357.0 | 17994.0 | 54546.0 | 34863.0 | 55505.0 | 42592.0 | 58503.0 | 54587.0 | 58810.0 | 50950.0 | 23419.0 | 15516.0 | 42457.0 | 60206.0 | 26907.0 | 19590.0 |
| sub-09\_ses-baseline | 4122.0 | 60343.0 | 63050.0 | 51336.0 | 79296.0 | 51039.0 | 62547.0 | 64936.0 | 8894.0 | 13678.0 | 17706.0 | 2348.0 | 8841.0 | 11413.0 | 17431.0 | 1041.0 | 4256.0 | 42551.0 | 39881.0 | 61205.0 | 55001.0 | 68333.0 | 64071.0 | 2092.0 | 620.0 | 7084.0 | 7115.0 | 45284.0 | 40865.0 | 45651.0 | 43334.0 | 57405.0 | 29813.0 | 26850.0 | 15888.0 | 18243.0 | 61410.0 | 61265.0 | 61966.0 | 69586.0 | 62889.0 | 61142.0 | 61759.0 | 59944.0 | 22654.0 | 20122.0 | 60263.0 | 69747.0 | 12900.0 | 15065.0 |
| sub-11\_ses-baseline | 959.0 | 31655.0 | 47946.0 | 48395.0 | 109202.0 | 55278.0 | 65233.0 | 82284.0 | 13109.0 | 10081.0 | 8449.0 | 2941.0 | 2887.0 | 12967.0 | 10736.0 | 2036.0 | 2066.0 | 39577.0 | 42317.0 | 67900.0 | 55137.0 | 75495.0 | 61559.0 | 2314.0 | 4409.0 | 7000.0 | 5713.0 | 54552.0 | 59862.0 | 47671.0 | 40713.0 | 28645.0 | 34496.0 | 26359.0 | 19698.0 | 20973.0 | 63221.0 | 42092.0 | 65499.0 | 39329.0 | 65168.0 | 49797.0 | 64510.0 | 54330.0 | 14001.0 | 19295.0 | 51697.0 | 42455.0 | 14881.0 | 18603.0 |

###### Tractometry: Bundle Volume: Columns

Uncheck the tick box to hide columns. Click and drag the handle on the left to change order. Table ID: `tractometry_volume_violin_table`

Show All
Show None

| Sort | Visible | Group | Column | Description | ID | Scale |
| --- | --- | --- | --- | --- | --- | --- |
| \|\| |  |  | AC | Bundle Volume for AC | `AC` |  |
| \|\| |  |  | AF\_L | Bundle Volume for AF\_L | `AF_L` |  |
| \|\| |  |  | AF\_R | Bundle Volume for AF\_R | `AF_R` |  |
| \|\| |  |  | CC\_Fr\_1 | Bundle Volume for CC\_Fr\_1 | `CC_Fr_1` |  |
| \|\| |  |  | CC\_Fr\_2 | Bundle Volume for CC\_Fr\_2 | `CC_Fr_2` |  |
| \|\| |  |  | CC\_Oc | Bundle Volume for CC\_Oc | `CC_Oc` |  |
| \|\| |  |  | CC\_Pa | Bundle Volume for CC\_Pa | `CC_Pa` |  |
| \|\| |  |  | CC\_Pr\_Po | Bundle Volume for CC\_Pr\_Po | `CC_Pr_Po` |  |
| \|\| |  |  | CC\_Te | Bundle Volume for CC\_Te | `CC_Te` |  |
| \|\| |  |  | CG\_L | Bundle Volume for CG\_L | `CG_L` |  |
| \|\| |  |  | CG\_L\_An | Bundle Volume for CG\_L\_An | `CG_L_An` |  |
| \|\| |  |  | CG\_L\_Po | Bundle Volume for CG\_L\_Po | `CG_L_Po` |  |
| \|\| |  |  | CG\_L\_curve | Bundle Volume for CG\_L\_curve | `CG_L_curve` |  |
| \|\| |  |  | CG\_R | Bundle Volume for CG\_R | `CG_R` |  |
| \|\| |  |  | CG\_R\_An | Bundle Volume for CG\_R\_An | `CG_R_An` |  |
| \|\| |  |  | CG\_R\_Po | Bundle Volume for CG\_R\_Po | `CG_R_Po` |  |
| \|\| |  |  | CG\_R\_curve | Bundle Volume for CG\_R\_curve | `CG_R_curve` |  |
| \|\| |  |  | FAT\_L | Bundle Volume for FAT\_L | `FAT_L` |  |
| \|\| |  |  | FAT\_R | Bundle Volume for FAT\_R | `FAT_R` |  |
| \|\| |  |  | FPT\_L | Bundle Volume for FPT\_L | `FPT_L` |  |
| \|\| |  |  | FPT\_L\_Brainstem | Bundle Volume for FPT\_L\_Brainstem | `FPT_L_Brainstem` |  |
| \|\| |  |  | FPT\_R | Bundle Volume for FPT\_R | `FPT_R` |  |
| \|\| |  |  | FPT\_R\_Brainstem | Bundle Volume for FPT\_R\_Brainstem | `FPT_R_Brainstem` |  |
| \|\| |  |  | FX\_L | Bundle Volume for FX\_L | `FX_L` |  |
| \|\| |  |  | FX\_R | Bundle Volume for FX\_R | `FX_R` |  |
| \|\| |  |  | ICP\_L | Bundle Volume for ICP\_L | `ICP_L` |  |
| \|\| |  |  | ICP\_R | Bundle Volume for ICP\_R | `ICP_R` |  |
| \|\| |  |  | IFOF\_L | Bundle Volume for IFOF\_L | `IFOF_L` |  |
| \|\| |  |  | IFOF\_R | Bundle Volume for IFOF\_R | `IFOF_R` |  |
| \|\| |  |  | ILF\_L | Bundle Volume for ILF\_L | `ILF_L` |  |
| \|\| |  |  | ILF\_R | Bundle Volume for ILF\_R | `ILF_R` |  |
| \|\| |  |  | MCP | Bundle Volume for MCP | `MCP` |  |
| \|\| |  |  | MdLF\_L | Bundle Volume for MdLF\_L | `MdLF_L` |  |
| \|\| |  |  | MdLF\_R | Bundle Volume for MdLF\_R | `MdLF_R` |  |
| \|\| |  |  | OR\_ML\_L | Bundle Volume for OR\_ML\_L | `OR_ML_L` |  |
| \|\| |  |  | OR\_ML\_R | Bundle Volume for OR\_ML\_R | `OR_ML_R` |  |
| \|\| |  |  | POPT\_L | Bundle Volume for POPT\_L | `POPT_L` |  |
| \|\| |  |  | POPT\_L\_Brainstem | Bundle Volume for POPT\_L\_Brainstem | `POPT_L_Brainstem` |  |
| \|\| |  |  | POPT\_R | Bundle Volume for POPT\_R | `POPT_R` |  |
| \|\| |  |  | POPT\_R\_Brainstem | Bundle Volume for POPT\_R\_Brainstem | `POPT_R_Brainstem` |  |
| \|\| |  |  | PYT\_L | Bundle Volume for PYT\_L | `PYT_L` |  |
| \|\| |  |  | PYT\_L\_Brainstem | Bundle Volume for PYT\_L\_Brainstem | `PYT_L_Brainstem` |  |
| \|\| |  |  | PYT\_R | Bundle Volume for PYT\_R | `PYT_R` |  |
| \|\| |  |  | PYT\_R\_Brainstem | Bundle Volume for PYT\_R\_Brainstem | `PYT_R_Brainstem` |  |
| \|\| |  |  | SCP\_L | Bundle Volume for SCP\_L | `SCP_L` |  |
| \|\| |  |  | SCP\_R | Bundle Volume for SCP\_R | `SCP_R` |  |
| \|\| |  |  | SLF\_L | Bundle Volume for SLF\_L | `SLF_L` |  |
| \|\| |  |  | SLF\_R | Bundle Volume for SLF\_R | `SLF_R` |  |
| \|\| |  |  | UF\_L | Bundle Volume for UF\_L | `UF_L` |  |
| \|\| |  |  | UF\_R | Bundle Volume for UF\_R | `UF_R` |  |

Close

##### Streamline Count 10

Distribution of streamline counts per bundle. You should look for extreme outliers, as it can indicate issues in bundle extraction. Too high streamline counts may indicate inclusion of spurious streamlines, while too low counts may indicate incomplete bundles.

###### AI Summary

Provider: , model:

Chat with Seqera AI

Table
 Export...

Copy prompt


Summarize plot

Created with MultiQC

Copy table

 Configure columns

 Sort by highlight

 Scatter plot

 Violin plot
Export as CSV...
Showing ^10^/_10_ rows and ^50^/_50_ columns.

Copy Prompt


Summarize table

| Sample Name | AC | AF\_L | AF\_R | CC\_Fr\_1 | CC\_Fr\_2 | CC\_Oc | CC\_Pa | CC\_Pr\_Po | CC\_Te | CG\_L | CG\_L\_An | CG\_L\_Po | CG\_L\_curve | CG\_R | CG\_R\_An | CG\_R\_Po | CG\_R\_curve | FAT\_L | FAT\_R | FPT\_L | FPT\_L\_Brainstem | FPT\_R | FPT\_R\_Brainstem | FX\_L | FX\_R | ICP\_L | ICP\_R | IFOF\_L | IFOF\_R | ILF\_L | ILF\_R | MCP | MdLF\_L | MdLF\_R | OR\_ML\_L | OR\_ML\_R | POPT\_L | POPT\_L\_Brainstem | POPT\_R | POPT\_R\_Brainstem | PYT\_L | PYT\_L\_Brainstem | PYT\_R | PYT\_R\_Brainstem | SCP\_L | SCP\_R | SLF\_L | SLF\_R | UF\_L | UF\_R |
| --- | --- | --- | --- | --- | --- | --- | --- | --- | --- | --- | --- | --- | --- | --- | --- | --- | --- | --- | --- | --- | --- | --- | --- | --- | --- | --- | --- | --- | --- | --- | --- | --- | --- | --- | --- | --- | --- | --- | --- | --- | --- | --- | --- | --- | --- | --- | --- | --- | --- | --- |
| sub-01\_ses-baseline | 273.0 | 12083.0 | 9781.0 | 21849.0 | 88306.0 | 13914.0 | 35236.0 | 78094.0 | 17.0 | 8751.0 | 6270.0 | 229.0 | 232.0 | 7382.0 | 6180.0 | 98.0 | 189.0 | 38215.0 | 13882.0 | 27481.0 | 10288.0 | 39201.0 | 22530.0 | 41.0 | 17.0 | 1682.0 | 6220.0 | 10660.0 | 24684.0 | 35095.0 | 23082.0 | 15246.0 | 1727.0 | 6048.0 | 8952.0 | 4658.0 | 18555.0 | 7945.0 | 27485.0 | 14413.0 | 36888.0 | 18944.0 | 48729.0 | 23297.0 | 703.0 | 926.0 | 5023.0 | 14313.0 | 4952.0 | 2745.0 |
| sub-02\_ses-baseline | 10.0 | 15286.0 | 9962.0 | 58212.0 | 45461.0 | 13576.0 | 21147.0 | 27215.0 | 20.0 | 11115.0 | 8527.0 | 301.0 | 41.0 | 9575.0 | 6790.0 | 517.0 | 65.0 | 62264.0 | 32987.0 | 64721.0 | 13103.0 | 62516.0 | 23347.0 | 5.0 | 19.0 | 1576.0 | 198.0 | 13452.0 | 13976.0 | 15568.0 | 14199.0 | 34668.0 | 5103.0 | 5013.0 | 9804.0 | 5484.0 | 34657.0 | 9886.0 | 14716.0 | 6615.0 | 77781.0 | 20190.0 | 62253.0 | 18832.0 | 2277.0 | 2092.0 | 22266.0 | 54890.0 | 1496.0 | 2415.0 |
| sub-03\_ses-baseline | 181.0 | 14503.0 | 1824.0 | 24597.0 | 34895.0 | 7292.0 | 55717.0 | 48200.0 | 397.0 | 440.0 | 82.0 | 861.0 | 11.0 | 5136.0 | 4863.0 | 292.0 | 33.0 | 49014.0 | 39237.0 | 37001.0 | 9158.0 | 67808.0 | 18933.0 | 57.0 |  | 2386.0 | 1359.0 | 27266.0 | 21529.0 | 30269.0 | 17586.0 | 13298.0 | 1269.0 | 2163.0 | 10194.0 | 10428.0 | 37304.0 | 3244.0 | 12848.0 | 3806.0 | 63637.0 | 10484.0 | 62068.0 | 15387.0 | 3787.0 | 1380.0 | 4904.0 | 6708.0 | 229.0 | 2719.0 |
| sub-04\_ses-baseline | 137.0 | 11780.0 | 27701.0 | 39683.0 | 54257.0 | 14428.0 | 26948.0 | 27027.0 | 1391.0 | 7779.0 | 6191.0 | 624.0 | 73.0 | 8790.0 | 6247.0 | 1190.0 | 94.0 | 42234.0 | 35950.0 | 47675.0 | 7582.0 | 32095.0 | 11842.0 | 18.0 | 91.0 | 6386.0 | 3502.0 | 3040.0 | 10771.0 | 44912.0 | 37482.0 | 12681.0 | 15746.0 | 9921.0 | 1347.0 | 2938.0 | 30675.0 | 9418.0 | 35321.0 | 10105.0 | 41504.0 | 12643.0 | 49637.0 | 15025.0 | 1135.0 | 3577.0 | 9634.0 | 9052.0 | 2022.0 | 4324.0 |
| sub-05\_ses-baseline | 3.0 | 16743.0 | 23871.0 | 16403.0 | 23304.0 | 14141.0 | 37054.0 | 23153.0 | 4.0 | 2877.0 | 1855.0 | 95.0 | 45.0 | 5451.0 | 5999.0 | 80.0 | 16.0 | 40244.0 | 58680.0 | 38921.0 | 5616.0 | 42045.0 | 15586.0 | 160.0 | 168.0 | 201.0 | 225.0 | 6426.0 | 7185.0 | 14628.0 | 28022.0 | 13939.0 | 268.0 | 1183.0 | 5029.0 | 5752.0 | 32106.0 | 8717.0 | 30459.0 | 10794.0 | 56248.0 | 16399.0 | 49946.0 | 24914.0 | 2854.0 | 1804.0 | 12685.0 | 13173.0 | 1362.0 | 1363.0 |
| sub-06\_ses-baseline | 14.0 | 17619.0 | 3029.0 | 32988.0 | 34569.0 | 9613.0 | 33415.0 | 30817.0 | 51.0 | 6513.0 | 5678.0 | 1131.0 | 35.0 | 5058.0 | 4331.0 | 409.0 | 72.0 | 31372.0 | 68715.0 | 18675.0 | 12890.0 | 27389.0 | 15245.0 | 177.0 | 65.0 | 3333.0 | 538.0 | 7550.0 | 4265.0 | 22542.0 | 8865.0 | 17922.0 | 453.0 | 742.0 | 7425.0 | 4738.0 | 36274.0 | 16074.0 | 30314.0 | 15833.0 | 50601.0 | 23290.0 | 55524.0 | 28494.0 | 1355.0 | 1146.0 | 6822.0 | 10783.0 | 4550.0 | 1841.0 |
| sub-07\_ses-baseline | 12.0 | 12227.0 | 11779.0 | 42733.0 | 53755.0 | 13748.0 | 37635.0 | 19411.0 | 7.0 | 426.0 | 374.0 | 46.0 | 73.0 | 7264.0 | 8466.0 | 203.0 |  | 25110.0 | 43410.0 | 31012.0 | 24267.0 | 41166.0 | 19961.0 | 1243.0 | 149.0 | 1296.0 | 938.0 | 14968.0 | 17506.0 | 12378.0 | 12505.0 | 12600.0 | 410.0 | 1826.0 | 9155.0 | 3767.0 | 13598.0 | 7694.0 | 7362.0 | 2949.0 | 60469.0 | 34297.0 | 52466.0 | 24121.0 | 4404.0 | 1473.0 | 7281.0 | 21082.0 | 3112.0 | 2905.0 |
| sub-08\_ses-baseline | 67.0 | 22212.0 | 10228.0 | 30870.0 | 38202.0 | 16346.0 | 24459.0 | 15798.0 | 246.0 | 10324.0 | 9528.0 | 525.0 | 230.0 | 5911.0 | 5474.0 | 428.0 | 33.0 | 32744.0 | 48068.0 | 40272.0 | 14111.0 | 26806.0 | 7740.0 | 337.0 | 563.0 | 535.0 | 1723.0 | 21078.0 | 21815.0 | 19999.0 | 14876.0 | 24231.0 | 5185.0 | 4036.0 | 7650.0 | 2968.0 | 19546.0 | 4214.0 | 11707.0 | 3155.0 | 44524.0 | 12599.0 | 29044.0 | 7719.0 | 3189.0 | 1518.0 | 8686.0 | 29413.0 | 4877.0 | 1203.0 |
| sub-09\_ses-baseline | 92.0 | 35339.0 | 35732.0 | 38517.0 | 41115.0 | 16348.0 | 29918.0 | 38492.0 | 380.0 | 6531.0 | 5914.0 | 276.0 | 225.0 | 7103.0 | 2613.0 | 119.0 | 86.0 | 64040.0 | 84808.0 | 21385.0 | 11940.0 | 50380.0 | 23269.0 | 30.0 | 7.0 | 1417.0 | 1805.0 | 9718.0 | 10733.0 | 29177.0 | 22339.0 | 27022.0 | 7405.0 | 5266.0 | 2787.0 | 5812.0 | 31333.0 | 21216.0 | 27609.0 | 29704.0 | 52690.0 | 33198.0 | 55908.0 | 34164.0 | 2983.0 | 4444.0 | 18366.0 | 37847.0 | 1284.0 | 1384.0 |
| sub-11\_ses-baseline | 19.0 | 4762.0 | 7285.0 | 22658.0 | 73437.0 | 13622.0 | 29674.0 | 42652.0 | 486.0 | 6609.0 | 5622.0 | 211.0 | 44.0 | 5486.0 | 5243.0 | 182.0 | 36.0 | 26097.0 | 36697.0 | 20740.0 | 3784.0 | 26960.0 | 7121.0 | 25.0 | 239.0 | 3723.0 | 1689.0 | 11830.0 | 18281.0 | 15601.0 | 9801.0 | 9825.0 | 8091.0 | 6503.0 | 3462.0 | 6345.0 | 23797.0 | 3876.0 | 21705.0 | 2539.0 | 47826.0 | 7803.0 | 48194.0 | 21114.0 | 1401.0 | 1922.0 | 4627.0 | 7220.0 | 1225.0 | 2059.0 |

###### Tractometry: Streamline Count: Columns

Uncheck the tick box to hide columns. Click and drag the handle on the left to change order. Table ID: `tractometry_streamlines_count_violin_table`

Show All
Show None

| Sort | Visible | Group | Column | Description | ID | Scale |
| --- | --- | --- | --- | --- | --- | --- |
| \|\| |  |  | AC | Streamline Count for AC | `AC` |  |
| \|\| |  |  | AF\_L | Streamline Count for AF\_L | `AF_L` |  |
| \|\| |  |  | AF\_R | Streamline Count for AF\_R | `AF_R` |  |
| \|\| |  |  | CC\_Fr\_1 | Streamline Count for CC\_Fr\_1 | `CC_Fr_1` |  |
| \|\| |  |  | CC\_Fr\_2 | Streamline Count for CC\_Fr\_2 | `CC_Fr_2` |  |
| \|\| |  |  | CC\_Oc | Streamline Count for CC\_Oc | `CC_Oc` |  |
| \|\| |  |  | CC\_Pa | Streamline Count for CC\_Pa | `CC_Pa` |  |
| \|\| |  |  | CC\_Pr\_Po | Streamline Count for CC\_Pr\_Po | `CC_Pr_Po` |  |
| \|\| |  |  | CC\_Te | Streamline Count for CC\_Te | `CC_Te` |  |
| \|\| |  |  | CG\_L | Streamline Count for CG\_L | `CG_L` |  |
| \|\| |  |  | CG\_L\_An | Streamline Count for CG\_L\_An | `CG_L_An` |  |
| \|\| |  |  | CG\_L\_Po | Streamline Count for CG\_L\_Po | `CG_L_Po` |  |
| \|\| |  |  | CG\_L\_curve | Streamline Count for CG\_L\_curve | `CG_L_curve` |  |
| \|\| |  |  | CG\_R | Streamline Count for CG\_R | `CG_R` |  |
| \|\| |  |  | CG\_R\_An | Streamline Count for CG\_R\_An | `CG_R_An` |  |
| \|\| |  |  | CG\_R\_Po | Streamline Count for CG\_R\_Po | `CG_R_Po` |  |
| \|\| |  |  | CG\_R\_curve | Streamline Count for CG\_R\_curve | `CG_R_curve` |  |
| \|\| |  |  | FAT\_L | Streamline Count for FAT\_L | `FAT_L` |  |
| \|\| |  |  | FAT\_R | Streamline Count for FAT\_R | `FAT_R` |  |
| \|\| |  |  | FPT\_L | Streamline Count for FPT\_L | `FPT_L` |  |
| \|\| |  |  | FPT\_L\_Brainstem | Streamline Count for FPT\_L\_Brainstem | `FPT_L_Brainstem` |  |
| \|\| |  |  | FPT\_R | Streamline Count for FPT\_R | `FPT_R` |  |
| \|\| |  |  | FPT\_R\_Brainstem | Streamline Count for FPT\_R\_Brainstem | `FPT_R_Brainstem` |  |
| \|\| |  |  | FX\_L | Streamline Count for FX\_L | `FX_L` |  |
| \|\| |  |  | FX\_R | Streamline Count for FX\_R | `FX_R` |  |
| \|\| |  |  | ICP\_L | Streamline Count for ICP\_L | `ICP_L` |  |
| \|\| |  |  | ICP\_R | Streamline Count for ICP\_R | `ICP_R` |  |
| \|\| |  |  | IFOF\_L | Streamline Count for IFOF\_L | `IFOF_L` |  |
| \|\| |  |  | IFOF\_R | Streamline Count for IFOF\_R | `IFOF_R` |  |
| \|\| |  |  | ILF\_L | Streamline Count for ILF\_L | `ILF_L` |  |
| \|\| |  |  | ILF\_R | Streamline Count for ILF\_R | `ILF_R` |  |
| \|\| |  |  | MCP | Streamline Count for MCP | `MCP` |  |
| \|\| |  |  | MdLF\_L | Streamline Count for MdLF\_L | `MdLF_L` |  |
| \|\| |  |  | MdLF\_R | Streamline Count for MdLF\_R | `MdLF_R` |  |
| \|\| |  |  | OR\_ML\_L | Streamline Count for OR\_ML\_L | `OR_ML_L` |  |
| \|\| |  |  | OR\_ML\_R | Streamline Count for OR\_ML\_R | `OR_ML_R` |  |
| \|\| |  |  | POPT\_L | Streamline Count for POPT\_L | `POPT_L` |  |
| \|\| |  |  | POPT\_L\_Brainstem | Streamline Count for POPT\_L\_Brainstem | `POPT_L_Brainstem` |  |
| \|\| |  |  | POPT\_R | Streamline Count for POPT\_R | `POPT_R` |  |
| \|\| |  |  | POPT\_R\_Brainstem | Streamline Count for POPT\_R\_Brainstem | `POPT_R_Brainstem` |  |
| \|\| |  |  | PYT\_L | Streamline Count for PYT\_L | `PYT_L` |  |
| \|\| |  |  | PYT\_L\_Brainstem | Streamline Count for PYT\_L\_Brainstem | `PYT_L_Brainstem` |  |
| \|\| |  |  | PYT\_R | Streamline Count for PYT\_R | `PYT_R` |  |
| \|\| |  |  | PYT\_R\_Brainstem | Streamline Count for PYT\_R\_Brainstem | `PYT_R_Brainstem` |  |
| \|\| |  |  | SCP\_L | Streamline Count for SCP\_L | `SCP_L` |  |
| \|\| |  |  | SCP\_R | Streamline Count for SCP\_R | `SCP_R` |  |
| \|\| |  |  | SLF\_L | Streamline Count for SLF\_L | `SLF_L` |  |
| \|\| |  |  | SLF\_R | Streamline Count for SLF\_R | `SLF_R` |  |
| \|\| |  |  | UF\_L | Streamline Count for UF\_L | `UF_L` |  |
| \|\| |  |  | UF\_R | Streamline Count for UF\_R | `UF_R` |  |

Close

#### Cortical Regions

Assessment of cortical region volumes for quality control using IQR-based outlier detection. Each cortical region's volume is evaluated across subjects, and regions with volumes falling outside the range are considered outliers. The percentage of outlier regions per subject is reported in the general statistics, with thresholds for pass/warn/fail configurable in the MultiQC configuration file.https://github.com/nf-neuro/MultiQC\_neuroimaging

##### Left Hemisphere Volume Distribution 10

Distribution of cortical region volumes in the left hemisphere across all samples. You may look for extreme outliers, which could indicate segmentation issues or data quality problems. Automatic detection of these outliers is based on the interquartile range (IQR) method, where values falling outside the range defined by Q1 - 3*IQR to Q3 + 3*IQR are considered outliers. Combined with other indicators, these outliers may help identify subjects that require further investigation or exclusion.

###### AI Summary

Provider: , model:

Chat with Seqera AI

Table
 Export...

Copy prompt


Summarize plot

Created with MultiQC

Copy table

 Configure columns

 Sort by highlight

 Scatter plot

 Violin plot
Export as CSV...
Showing ^10^/_10_ rows and ^94^/_94_ columns.

Copy Prompt


Summarize table

| Sample Name | lh\_SFG\_L\_6\_1 | lh\_SFG\_L\_6\_2 | lh\_SFG\_L\_6\_3 | lh\_SFG\_L\_6\_4 | lh\_SFG\_L\_6\_5 | lh\_SFG\_L\_6\_6 | lh\_MFG\_L\_7\_1 | lh\_MFG\_L\_7\_2 | lh\_MFG\_L\_7\_3 | lh\_MFG\_L\_7\_4 | lh\_MFG\_L\_7\_5 | lh\_MFG\_L\_7\_6 | lh\_MFG\_L\_7\_7 | lh\_IFG\_L\_6\_1 | lh\_IFG\_L\_6\_2 | lh\_IFG\_L\_6\_3 | lh\_IFG\_L\_6\_4 | lh\_IFG\_L\_6\_5 | lh\_IFG\_L\_6\_6 | lh\_OrG\_L\_6\_1 | lh\_OrG\_L\_6\_2 | lh\_OrG\_L\_6\_3 | lh\_OrG\_L\_6\_4 | lh\_OrG\_L\_6\_5 | lh\_OrG\_L\_6\_6 | lh\_PrG\_L\_6\_1 | lh\_PrG\_L\_6\_2 | lh\_PrG\_L\_6\_3 | lh\_PrG\_L\_6\_4 | lh\_PrG\_L\_6\_5 | lh\_PrG\_L\_6\_6 | lh\_PCL\_L\_2\_1 | lh\_PCL\_L\_2\_2 | lh\_STG\_L\_6\_1 | lh\_STG\_L\_6\_2 | lh\_STG\_L\_6\_3 | lh\_STG\_L\_6\_4 | lh\_STG\_L\_6\_5 | lh\_STG\_L\_6\_6 | lh\_MTG\_L\_4\_1 | lh\_MTG\_L\_4\_2 | lh\_MTG\_L\_4\_3 | lh\_MTG\_L\_4\_4 | lh\_ITG\_L\_5\_1 | lh\_ITG\_L\_5\_2 | lh\_ITG\_L\_5\_3 | lh\_ITG\_L\_5\_4 | lh\_ITG\_L\_5\_5 | lh\_FuG\_L\_3\_1 | lh\_FuG\_L\_3\_2 | lh\_FuG\_L\_3\_3 | lh\_PhG\_L\_6\_1 | lh\_PhG\_L\_6\_2 | lh\_PhG\_L\_6\_3 | lh\_PhG\_L\_6\_4 | lh\_PhG\_L\_6\_5 | lh\_PhG\_L\_6\_6 | lh\_pSTS\_L\_2\_1 | lh\_pSTS\_L\_2\_2 | lh\_SPL\_L\_4\_1 | lh\_SPL\_L\_4\_2 | lh\_SPL\_L\_4\_3 | lh\_SPL\_L\_4\_4 | lh\_IPL\_L\_6\_1 | lh\_IPL\_L\_6\_2 | lh\_IPL\_L\_6\_3 | lh\_IPL\_L\_6\_4 | lh\_IPL\_L\_6\_5 | lh\_IPL\_L\_6\_6 | lh\_PCun\_L\_4\_1 | lh\_PCun\_L\_4\_2 | lh\_PCun\_L\_4\_3 | lh\_PCun\_L\_4\_4 | lh\_PoG\_L\_4\_1 | lh\_PoG\_L\_4\_2 | lh\_PoG\_L\_4\_3 | lh\_PoG\_L\_4\_4 | lh\_INS\_L\_6\_1 | lh\_INS\_L\_6\_2 | lh\_INS\_L\_6\_3 | lh\_CG\_L\_5\_1 | lh\_CG\_L\_5\_2 | lh\_CG\_L\_5\_3 | lh\_CG\_L\_5\_4 | lh\_CG\_L\_5\_5 | lh\_MVOcC\_L\_5\_1 | lh\_MVOcC\_L\_5\_2 | lh\_MVOcC\_L\_5\_3 | lh\_MVOcC\_L\_5\_4 | lh\_MVOcC\_L\_5\_5 | lh\_LOcC\_L\_4\_1 | lh\_LOcC\_L\_4\_2 | lh\_LOcC\_L\_4\_3 | lh\_LOcC\_L\_4\_4 |
| --- | --- | --- | --- | --- | --- | --- | --- | --- | --- | --- | --- | --- | --- | --- | --- | --- | --- | --- | --- | --- | --- | --- | --- | --- | --- | --- | --- | --- | --- | --- | --- | --- | --- | --- | --- | --- | --- | --- | --- | --- | --- | --- | --- | --- | --- | --- | --- | --- | --- | --- | --- | --- | --- | --- | --- | --- | --- | --- | --- | --- | --- | --- | --- | --- | --- | --- | --- | --- | --- | --- | --- | --- | --- | --- | --- | --- | --- | --- | --- | --- | --- | --- | --- | --- | --- | --- | --- | --- | --- | --- | --- | --- | --- | --- |
| sub-01\_ses-baseline | 1832.0 | 2045.0 | 5299.0 | 5297.0 | 2391.0 | 2096.0 | 3453.0 | 5782.0 | 1877.0 | 2565.0 | 5728.0 | 3975.0 | 850.0 | 1962.0 | 1061.0 | 1642.0 | 1280.0 | 2077.0 | 1924.0 | 936.0 | 2660.0 | 2205.0 | 3885.0 | 1868.0 | 858.0 | 2418.0 | 1345.0 | 2554.0 | 2064.0 | 2339.0 | 1731.0 | 2076.0 | 1535.0 | 1304.0 | 1678.0 | 1459.0 | 3013.0 | 2392.0 | 3900.0 | 2236.0 | 3057.0 | 2023.0 | 2116.0 | 4359.0 | 1316.0 | 1330.0 | 2808.0 | 1893.0 | 1900.0 | 2205.0 | 2529.0 | 2770.0 | 1251.0 | 115.0 | 844.0 | 21.0 | 183.0 | 344.0 | 1608.0 | 799.0 | 4830.0 | 2565.0 | 1610.0 | 2386.0 | 4603.0 | 5597.0 | 2136.0 | 5524.0 | 3762.0 | 3608.0 | 2416.0 | 1502.0 | 1955.0 | 2440.0 | 3246.0 | 2304.0 | 2546.0 | 1093.0 | 815.0 | 4125.0 | 2594.0 | 3082.0 | 1205.0 | 536.0 | 2096.0 | 1971.0 | 863.0 | 2016.0 | 712.0 | 1926.0 | 1790.0 | 2269.0 | 2118.0 | 3109.0 |
| sub-02\_ses-baseline | 1905.0 | 2151.0 | 5873.0 | 6457.0 | 2484.0 | 1904.0 | 4305.0 | 5865.0 | 1774.0 | 4405.0 | 5690.0 | 3820.0 | 551.0 | 2263.0 | 852.0 | 1224.0 | 1319.0 | 2537.0 | 2172.0 | 1172.0 | 3380.0 | 2828.0 | 5288.0 | 1867.0 | 987.0 | 2618.0 | 1594.0 | 2482.0 | 2165.0 | 2241.0 | 2010.0 | 2040.0 | 1883.0 | 1475.0 | 1789.0 | 1289.0 | 3409.0 | 2529.0 | 4583.0 | 2203.0 | 3801.0 | 1472.0 | 3547.0 | 3938.0 | 2056.0 | 1760.0 | 3211.0 | 2999.0 | 2080.0 | 2927.0 | 3231.0 | 2789.0 | 910.0 | 163.0 | 577.0 | 60.0 | 325.0 | 403.0 | 1707.0 | 1289.0 | 5310.0 | 3647.0 | 2208.0 | 2850.0 | 7443.0 | 3862.0 | 2217.0 | 4430.0 | 7675.0 | 2762.0 | 2265.0 | 1807.0 | 2752.0 | 2541.0 | 2660.0 | 2612.0 | 2081.0 | 1436.0 | 1139.0 | 4796.0 | 2926.0 | 3453.0 | 1294.0 | 635.0 | 2276.0 | 2531.0 | 959.0 | 1910.0 | 603.0 | 1817.0 | 1780.0 | 2550.0 | 2570.0 | 3511.0 |
| sub-03\_ses-baseline | 1453.0 | 2141.0 | 3792.0 | 4596.0 | 2471.0 | 2791.0 | 3338.0 | 4531.0 | 1371.0 | 3206.0 | 5047.0 | 2333.0 | 567.0 | 2022.0 | 1005.0 | 1884.0 | 1245.0 | 1951.0 | 1813.0 | 759.0 | 2744.0 | 2758.0 | 3822.0 | 2031.0 | 918.0 | 1842.0 | 1143.0 | 2779.0 | 2393.0 | 2404.0 | 1714.0 | 1812.0 | 2026.0 | 1902.0 | 2059.0 | 1603.0 | 3575.0 | 2683.0 | 4199.0 | 2012.0 | 2928.0 | 2076.0 | 1835.0 | 3940.0 | 1088.0 | 1841.0 | 2998.0 | 1898.0 | 1532.0 | 2816.0 | 2735.0 | 2622.0 | 381.0 | 112.0 | 390.0 | 38.0 | 274.0 | 296.0 | 2108.0 | 825.0 | 3974.0 | 2969.0 | 1808.0 | 2665.0 | 5633.0 | 4253.0 | 1937.0 | 5148.0 | 4760.0 | 2964.0 | 2178.0 | 1589.0 | 1750.0 | 2443.0 | 2396.0 | 1915.0 | 1861.0 | 1139.0 | 578.0 | 3738.0 | 2747.0 | 3126.0 | 1092.0 | 594.0 | 1927.0 | 1859.0 | 1159.0 | 2003.0 | 634.0 | 2543.0 | 1531.0 | 2382.0 | 2336.0 | 3131.0 |
| sub-04\_ses-baseline | 1739.0 | 2528.0 | 5688.0 | 5577.0 | 2955.0 | 2561.0 | 3775.0 | 4554.0 | 1541.0 | 3280.0 | 4726.0 | 3758.0 | 795.0 | 2194.0 | 827.0 | 1229.0 | 1074.0 | 1738.0 | 1728.0 | 805.0 | 2409.0 | 2342.0 | 3984.0 | 2049.0 | 861.0 | 2247.0 | 1250.0 | 3063.0 | 2097.0 | 2352.0 | 1637.0 | 1759.0 | 1541.0 | 1562.0 | 2013.0 | 1005.0 | 2672.0 | 1973.0 | 4453.0 | 1720.0 | 3103.0 | 2083.0 | 2477.0 | 3751.0 | 1674.0 | 1560.0 | 2772.0 | 2004.0 | 1782.0 | 2124.0 | 2148.0 | 1945.0 | 828.0 | 97.0 | 731.0 | 22.0 | 237.0 | 333.0 | 1579.0 | 1328.0 | 3750.0 | 3299.0 | 1760.0 | 2531.0 | 4117.0 | 5279.0 | 2668.0 | 5015.0 | 3883.0 | 2538.0 | 2581.0 | 1524.0 | 1757.0 | 2306.0 | 2534.0 | 1896.0 | 1916.0 | 1198.0 | 748.0 | 3820.0 | 2571.0 | 2985.0 | 1177.0 | 698.0 | 2059.0 | 2102.0 | 1208.0 | 2581.0 | 1014.0 | 2585.0 | 2294.0 | 2061.0 | 1934.0 | 3071.0 |
| sub-05\_ses-baseline | 2059.0 | 2158.0 | 5963.0 | 6544.0 | 2478.0 | 2497.0 | 4785.0 | 6112.0 | 2940.0 | 3165.0 | 4627.0 | 5296.0 | 1040.0 | 2751.0 | 781.0 | 1184.0 | 1221.0 | 1792.0 | 1944.0 | 1164.0 | 3194.0 | 2937.0 | 4122.0 | 2540.0 | 1093.0 | 2385.0 | 1957.0 | 2441.0 | 2407.0 | 2358.0 | 2028.0 | 2786.0 | 1946.0 | 1629.0 | 2188.0 | 1999.0 | 3805.0 | 3528.0 | 4833.0 | 1933.0 | 4904.0 | 3078.0 | 3062.0 | 6312.0 | 2255.0 | 1690.0 | 3919.0 | 2592.0 | 2234.0 | 3357.0 | 3267.0 | 3365.0 | 604.0 | 139.0 | 449.0 | 26.0 | 301.0 | 321.0 | 2437.0 | 1552.0 | 4561.0 | 2148.0 | 2081.0 | 2910.0 | 6075.0 | 7125.0 | 2236.0 | 6781.0 | 6627.0 | 3271.0 | 2723.0 | 1953.0 | 2512.0 | 3029.0 | 3116.0 | 2885.0 | 2115.0 | 1494.0 | 1075.0 | 4921.0 | 3035.0 | 3887.0 | 1345.0 | 852.0 | 2536.0 | 2212.0 | 1379.0 | 2338.0 | 759.0 | 2067.0 | 2234.0 | 2149.0 | 2436.0 | 3022.0 |
| sub-06\_ses-baseline | 1717.0 | 2076.0 | 7067.0 | 6095.0 | 3579.0 | 2804.0 | 4122.0 | 6121.0 | 1937.0 | 3817.0 | 6462.0 | 3460.0 | 679.0 | 1953.0 | 1082.0 | 1326.0 | 1282.0 | 2085.0 | 1916.0 | 1078.0 | 3202.0 | 2382.0 | 4766.0 | 2039.0 | 1009.0 | 1980.0 | 1219.0 | 2787.0 | 2629.0 | 2373.0 | 1909.0 | 2061.0 | 2088.0 | 1626.0 | 1841.0 | 1416.0 | 3384.0 | 2589.0 | 4855.0 | 2452.0 | 3281.0 | 2740.0 | 2765.0 | 4557.0 | 1728.0 | 2293.0 | 4044.0 | 2079.0 | 2103.0 | 2299.0 | 2944.0 | 2426.0 | 742.0 | 132.0 | 428.0 | 22.0 | 258.0 | 338.0 | 1640.0 | 1045.0 | 4083.0 | 3003.0 | 1857.0 | 2988.0 | 5446.0 | 4130.0 | 2369.0 | 5607.0 | 4579.0 | 3044.0 | 2302.0 | 2143.0 | 2673.0 | 2717.0 | 3146.0 | 2141.0 | 2608.0 | 1981.0 | 791.0 | 4557.0 | 2863.0 | 3852.0 | 1179.0 | 830.0 | 2672.0 | 2072.0 | 1275.0 | 2370.0 | 776.0 | 2629.0 | 2250.0 | 3642.0 | 3272.0 | 3949.0 |
| sub-07\_ses-baseline | 2806.0 | 3208.0 | 5620.0 | 9978.0 | 2428.0 | 2798.0 | 5132.0 | 8022.0 | 2185.0 | 4968.0 | 6072.0 | 3467.0 | 655.0 | 2777.0 | 1233.0 | 1279.0 | 1367.0 | 2352.0 | 2209.0 | 1615.0 | 3443.0 | 3036.0 | 5593.0 | 2481.0 | 1283.0 | 2797.0 | 1747.0 | 2675.0 | 2668.0 | 2662.0 | 2356.0 | 2278.0 | 2361.0 | 2196.0 | 2485.0 | 1918.0 | 4158.0 | 3325.0 | 5128.0 | 2277.0 | 4005.0 | 2971.0 | 2989.0 | 5611.0 | 1677.0 | 2384.0 | 4281.0 | 1953.0 | 2542.0 | 3290.0 | 3279.0 | 2810.0 | 1898.0 | 303.0 | 1301.0 | 72.0 | 422.0 | 599.0 | 2976.0 | 1509.0 | 6400.0 | 3448.0 | 2292.0 | 3430.0 | 8217.0 | 5786.0 | 2444.0 | 6431.0 | 6953.0 | 3965.0 | 2256.0 | 1550.0 | 3747.0 | 2851.0 | 3831.0 | 2942.0 | 3458.0 | 1663.0 | 1222.0 | 5007.0 | 3763.0 | 3923.0 | 1868.0 | 959.0 | 2750.0 | 3220.0 | 1451.0 | 2314.0 | 923.0 | 3186.0 | 2628.0 | 2007.0 | 2102.0 | 3241.0 |
| sub-08\_ses-baseline | 2633.0 | 2217.0 | 5569.0 | 7409.0 | 1891.0 | 2080.0 | 4224.0 | 8034.0 | 2310.0 | 3391.0 | 5594.0 | 5157.0 | 833.0 | 2342.0 | 754.0 | 1354.0 | 1094.0 | 1638.0 | 1763.0 | 902.0 | 3259.0 | 2710.0 | 5038.0 | 2065.0 | 1141.0 | 2499.0 | 1421.0 | 2484.0 | 2439.0 | 2112.0 | 1694.0 | 2452.0 | 1651.0 | 1450.0 | 1947.0 | 1895.0 | 3785.0 | 3057.0 | 5299.0 | 2214.0 | 3795.0 | 2562.0 | 2316.0 | 5063.0 | 1717.0 | 2092.0 | 3849.0 | 2369.0 | 1758.0 | 2728.0 | 3637.0 | 3431.0 | 1210.0 | 298.0 | 1157.0 | 37.0 | 310.0 | 480.0 | 2185.0 | 1355.0 | 4478.0 | 2945.0 | 2000.0 | 2462.0 | 5598.0 | 5212.0 | 2212.0 | 6339.0 | 5308.0 | 3623.0 | 2399.0 | 1847.0 | 2973.0 | 3255.0 | 3507.0 | 2462.0 | 3193.0 | 1823.0 | 1222.0 | 4246.0 | 2547.0 | 3950.0 | 1317.0 | 832.0 | 2431.0 | 2618.0 | 1309.0 | 2373.0 | 856.0 | 2409.0 | 2363.0 | 2647.0 | 3100.0 | 3353.0 |
| sub-09\_ses-baseline | 1281.0 | 1828.0 | 5346.0 | 5534.0 | 2013.0 | 1706.0 | 3236.0 | 4346.0 | 1593.0 | 3475.0 | 3083.0 | 2813.0 | 473.0 | 2319.0 | 511.0 | 723.0 | 933.0 | 1472.0 | 1573.0 | 718.0 | 2634.0 | 2303.0 | 3576.0 | 2058.0 | 816.0 | 2122.0 | 1141.0 | 2091.0 | 1990.0 | 2084.0 | 1552.0 | 1655.0 | 1367.0 | 1378.0 | 1513.0 | 1240.0 | 2464.0 | 1659.0 | 4268.0 | 1637.0 | 2338.0 | 1989.0 | 1769.0 | 3665.0 | 1721.0 | 1453.0 | 2842.0 | 1874.0 | 1488.0 | 2083.0 | 2215.0 | 2200.0 | 1089.0 | 140.0 | 800.0 | 24.0 | 201.0 | 377.0 | 1486.0 | 983.0 | 3908.0 | 2732.0 | 1499.0 | 2279.0 | 3714.0 | 4113.0 | 2017.0 | 3928.0 | 3770.0 | 2631.0 | 1380.0 | 1075.0 | 1990.0 | 1729.0 | 2457.0 | 1976.0 | 1700.0 | 1193.0 | 376.0 | 3177.0 | 2151.0 | 2458.0 | 1209.0 | 595.0 | 2132.0 | 1653.0 | 875.0 | 2098.0 | 787.0 | 2191.0 | 2170.0 | 2283.0 | 2352.0 | 2978.0 |
| sub-11\_ses-baseline | 2604.0 | 2640.0 | 5244.0 | 7586.0 | 2788.0 | 2498.0 | 5164.0 | 7452.0 | 2267.0 | 5020.0 | 6869.0 | 4510.0 | 955.0 | 2817.0 | 1188.0 | 1441.0 | 1579.0 | 2725.0 | 2284.0 | 1240.0 | 3641.0 | 3391.0 | 5000.0 | 2502.0 | 1064.0 | 2641.0 | 1487.0 | 3064.0 | 2889.0 | 2556.0 | 2149.0 | 2924.0 | 2027.0 | 1731.0 | 2018.0 | 1924.0 | 3544.0 | 3477.0 | 5340.0 | 1852.0 | 2507.0 | 2708.0 | 3028.0 | 4889.0 | 1680.0 | 1867.0 | 2731.0 | 1994.0 | 1609.0 | 2685.0 | 3139.0 | 2777.0 | 859.0 | 110.0 | 540.0 | 26.0 | 265.0 | 453.0 | 3027.0 | 1559.0 | 5681.0 | 4008.0 | 2181.0 | 3163.0 | 5695.0 | 4837.0 | 2545.0 | 5741.0 | 5502.0 | 4372.0 | 2029.0 | 1711.0 | 2473.0 | 3038.0 | 3195.0 | 3557.0 | 2641.0 | 1661.0 | 1124.0 | 5234.0 | 3300.0 | 3936.0 | 1661.0 | 887.0 | 2326.0 | 3114.0 | 1223.0 | 3136.0 | 1288.0 | 2845.0 | 3353.0 | 2880.0 | 2806.0 | 4234.0 |

###### Cortical Regions: Left Hemisphere Volume Distribution: Columns

Uncheck the tick box to hide columns. Click and drag the handle on the left to change order. Table ID: `cortical_lh_volume_plot_table`

Show All
Show None

| Sort | Visible | Group | Column | Description | ID | Scale |
| --- | --- | --- | --- | --- | --- | --- |
| \|\| |  |  | lh\_SFG\_L\_6\_1 | Volume for lh\_SFG\_L\_6\_1 | `lh_SFG_L_6_1` |  |
| \|\| |  |  | lh\_SFG\_L\_6\_2 | Volume for lh\_SFG\_L\_6\_2 | `lh_SFG_L_6_2` |  |
| \|\| |  |  | lh\_SFG\_L\_6\_3 | Volume for lh\_SFG\_L\_6\_3 | `lh_SFG_L_6_3` |  |
| \|\| |  |  | lh\_SFG\_L\_6\_4 | Volume for lh\_SFG\_L\_6\_4 | `lh_SFG_L_6_4` |  |
| \|\| |  |  | lh\_SFG\_L\_6\_5 | Volume for lh\_SFG\_L\_6\_5 | `lh_SFG_L_6_5` |  |
| \|\| |  |  | lh\_SFG\_L\_6\_6 | Volume for lh\_SFG\_L\_6\_6 | `lh_SFG_L_6_6` |  |
| \|\| |  |  | lh\_MFG\_L\_7\_1 | Volume for lh\_MFG\_L\_7\_1 | `lh_MFG_L_7_1` |  |
| \|\| |  |  | lh\_MFG\_L\_7\_2 | Volume for lh\_MFG\_L\_7\_2 | `lh_MFG_L_7_2` |  |
| \|\| |  |  | lh\_MFG\_L\_7\_3 | Volume for lh\_MFG\_L\_7\_3 | `lh_MFG_L_7_3` |  |
| \|\| |  |  | lh\_MFG\_L\_7\_4 | Volume for lh\_MFG\_L\_7\_4 | `lh_MFG_L_7_4` |  |
| \|\| |  |  | lh\_MFG\_L\_7\_5 | Volume for lh\_MFG\_L\_7\_5 | `lh_MFG_L_7_5` |  |
| \|\| |  |  | lh\_MFG\_L\_7\_6 | Volume for lh\_MFG\_L\_7\_6 | `lh_MFG_L_7_6` |  |
| \|\| |  |  | lh\_MFG\_L\_7\_7 | Volume for lh\_MFG\_L\_7\_7 | `lh_MFG_L_7_7` |  |
| \|\| |  |  | lh\_IFG\_L\_6\_1 | Volume for lh\_IFG\_L\_6\_1 | `lh_IFG_L_6_1` |  |
| \|\| |  |  | lh\_IFG\_L\_6\_2 | Volume for lh\_IFG\_L\_6\_2 | `lh_IFG_L_6_2` |  |
| \|\| |  |  | lh\_IFG\_L\_6\_3 | Volume for lh\_IFG\_L\_6\_3 | `lh_IFG_L_6_3` |  |
| \|\| |  |  | lh\_IFG\_L\_6\_4 | Volume for lh\_IFG\_L\_6\_4 | `lh_IFG_L_6_4` |  |
| \|\| |  |  | lh\_IFG\_L\_6\_5 | Volume for lh\_IFG\_L\_6\_5 | `lh_IFG_L_6_5` |  |
| \|\| |  |  | lh\_IFG\_L\_6\_6 | Volume for lh\_IFG\_L\_6\_6 | `lh_IFG_L_6_6` |  |
| \|\| |  |  | lh\_OrG\_L\_6\_1 | Volume for lh\_OrG\_L\_6\_1 | `lh_OrG_L_6_1` |  |
| \|\| |  |  | lh\_OrG\_L\_6\_2 | Volume for lh\_OrG\_L\_6\_2 | `lh_OrG_L_6_2` |  |
| \|\| |  |  | lh\_OrG\_L\_6\_3 | Volume for lh\_OrG\_L\_6\_3 | `lh_OrG_L_6_3` |  |
| \|\| |  |  | lh\_OrG\_L\_6\_4 | Volume for lh\_OrG\_L\_6\_4 | `lh_OrG_L_6_4` |  |
| \|\| |  |  | lh\_OrG\_L\_6\_5 | Volume for lh\_OrG\_L\_6\_5 | `lh_OrG_L_6_5` |  |
| \|\| |  |  | lh\_OrG\_L\_6\_6 | Volume for lh\_OrG\_L\_6\_6 | `lh_OrG_L_6_6` |  |
| \|\| |  |  | lh\_PrG\_L\_6\_1 | Volume for lh\_PrG\_L\_6\_1 | `lh_PrG_L_6_1` |  |
| \|\| |  |  | lh\_PrG\_L\_6\_2 | Volume for lh\_PrG\_L\_6\_2 | `lh_PrG_L_6_2` |  |
| \|\| |  |  | lh\_PrG\_L\_6\_3 | Volume for lh\_PrG\_L\_6\_3 | `lh_PrG_L_6_3` |  |
| \|\| |  |  | lh\_PrG\_L\_6\_4 | Volume for lh\_PrG\_L\_6\_4 | `lh_PrG_L_6_4` |  |
| \|\| |  |  | lh\_PrG\_L\_6\_5 | Volume for lh\_PrG\_L\_6\_5 | `lh_PrG_L_6_5` |  |
| \|\| |  |  | lh\_PrG\_L\_6\_6 | Volume for lh\_PrG\_L\_6\_6 | `lh_PrG_L_6_6` |  |
| \|\| |  |  | lh\_PCL\_L\_2\_1 | Volume for lh\_PCL\_L\_2\_1 | `lh_PCL_L_2_1` |  |
| \|\| |  |  | lh\_PCL\_L\_2\_2 | Volume for lh\_PCL\_L\_2\_2 | `lh_PCL_L_2_2` |  |
| \|\| |  |  | lh\_STG\_L\_6\_1 | Volume for lh\_STG\_L\_6\_1 | `lh_STG_L_6_1` |  |
| \|\| |  |  | lh\_STG\_L\_6\_2 | Volume for lh\_STG\_L\_6\_2 | `lh_STG_L_6_2` |  |
| \|\| |  |  | lh\_STG\_L\_6\_3 | Volume for lh\_STG\_L\_6\_3 | `lh_STG_L_6_3` |  |
| \|\| |  |  | lh\_STG\_L\_6\_4 | Volume for lh\_STG\_L\_6\_4 | `lh_STG_L_6_4` |  |
| \|\| |  |  | lh\_STG\_L\_6\_5 | Volume for lh\_STG\_L\_6\_5 | `lh_STG_L_6_5` |  |
| \|\| |  |  | lh\_STG\_L\_6\_6 | Volume for lh\_STG\_L\_6\_6 | `lh_STG_L_6_6` |  |
| \|\| |  |  | lh\_MTG\_L\_4\_1 | Volume for lh\_MTG\_L\_4\_1 | `lh_MTG_L_4_1` |  |
| \|\| |  |  | lh\_MTG\_L\_4\_2 | Volume for lh\_MTG\_L\_4\_2 | `lh_MTG_L_4_2` |  |
| \|\| |  |  | lh\_MTG\_L\_4\_3 | Volume for lh\_MTG\_L\_4\_3 | `lh_MTG_L_4_3` |  |
| \|\| |  |  | lh\_MTG\_L\_4\_4 | Volume for lh\_MTG\_L\_4\_4 | `lh_MTG_L_4_4` |  |
| \|\| |  |  | lh\_ITG\_L\_5\_1 | Volume for lh\_ITG\_L\_5\_1 | `lh_ITG_L_5_1` |  |
| \|\| |  |  | lh\_ITG\_L\_5\_2 | Volume for lh\_ITG\_L\_5\_2 | `lh_ITG_L_5_2` |  |
| \|\| |  |  | lh\_ITG\_L\_5\_3 | Volume for lh\_ITG\_L\_5\_3 | `lh_ITG_L_5_3` |  |
| \|\| |  |  | lh\_ITG\_L\_5\_4 | Volume for lh\_ITG\_L\_5\_4 | `lh_ITG_L_5_4` |  |
| \|\| |  |  | lh\_ITG\_L\_5\_5 | Volume for lh\_ITG\_L\_5\_5 | `lh_ITG_L_5_5` |  |
| \|\| |  |  | lh\_FuG\_L\_3\_1 | Volume for lh\_FuG\_L\_3\_1 | `lh_FuG_L_3_1` |  |
| \|\| |  |  | lh\_FuG\_L\_3\_2 | Volume for lh\_FuG\_L\_3\_2 | `lh_FuG_L_3_2` |  |
| \|\| |  |  | lh\_FuG\_L\_3\_3 | Volume for lh\_FuG\_L\_3\_3 | `lh_FuG_L_3_3` |  |
| \|\| |  |  | lh\_PhG\_L\_6\_1 | Volume for lh\_PhG\_L\_6\_1 | `lh_PhG_L_6_1` |  |
| \|\| |  |  | lh\_PhG\_L\_6\_2 | Volume for lh\_PhG\_L\_6\_2 | `lh_PhG_L_6_2` |  |
| \|\| |  |  | lh\_PhG\_L\_6\_3 | Volume for lh\_PhG\_L\_6\_3 | `lh_PhG_L_6_3` |  |
| \|\| |  |  | lh\_PhG\_L\_6\_4 | Volume for lh\_PhG\_L\_6\_4 | `lh_PhG_L_6_4` |  |
| \|\| |  |  | lh\_PhG\_L\_6\_5 | Volume for lh\_PhG\_L\_6\_5 | `lh_PhG_L_6_5` |  |
| \|\| |  |  | lh\_PhG\_L\_6\_6 | Volume for lh\_PhG\_L\_6\_6 | `lh_PhG_L_6_6` |  |
| \|\| |  |  | lh\_pSTS\_L\_2\_1 | Volume for lh\_pSTS\_L\_2\_1 | `lh_pSTS_L_2_1` |  |
| \|\| |  |  | lh\_pSTS\_L\_2\_2 | Volume for lh\_pSTS\_L\_2\_2 | `lh_pSTS_L_2_2` |  |
| \|\| |  |  | lh\_SPL\_L\_4\_1 | Volume for lh\_SPL\_L\_4\_1 | `lh_SPL_L_4_1` |  |
| \|\| |  |  | lh\_SPL\_L\_4\_2 | Volume for lh\_SPL\_L\_4\_2 | `lh_SPL_L_4_2` |  |
| \|\| |  |  | lh\_SPL\_L\_4\_3 | Volume for lh\_SPL\_L\_4\_3 | `lh_SPL_L_4_3` |  |
| \|\| |  |  | lh\_SPL\_L\_4\_4 | Volume for lh\_SPL\_L\_4\_4 | `lh_SPL_L_4_4` |  |
| \|\| |  |  | lh\_IPL\_L\_6\_1 | Volume for lh\_IPL\_L\_6\_1 | `lh_IPL_L_6_1` |  |
| \|\| |  |  | lh\_IPL\_L\_6\_2 | Volume for lh\_IPL\_L\_6\_2 | `lh_IPL_L_6_2` |  |
| \|\| |  |  | lh\_IPL\_L\_6\_3 | Volume for lh\_IPL\_L\_6\_3 | `lh_IPL_L_6_3` |  |
| \|\| |  |  | lh\_IPL\_L\_6\_4 | Volume for lh\_IPL\_L\_6\_4 | `lh_IPL_L_6_4` |  |
| \|\| |  |  | lh\_IPL\_L\_6\_5 | Volume for lh\_IPL\_L\_6\_5 | `lh_IPL_L_6_5` |  |
| \|\| |  |  | lh\_IPL\_L\_6\_6 | Volume for lh\_IPL\_L\_6\_6 | `lh_IPL_L_6_6` |  |
| \|\| |  |  | lh\_PCun\_L\_4\_1 | Volume for lh\_PCun\_L\_4\_1 | `lh_PCun_L_4_1` |  |
| \|\| |  |  | lh\_PCun\_L\_4\_2 | Volume for lh\_PCun\_L\_4\_2 | `lh_PCun_L_4_2` |  |
| \|\| |  |  | lh\_PCun\_L\_4\_3 | Volume for lh\_PCun\_L\_4\_3 | `lh_PCun_L_4_3` |  |
| \|\| |  |  | lh\_PCun\_L\_4\_4 | Volume for lh\_PCun\_L\_4\_4 | `lh_PCun_L_4_4` |  |
| \|\| |  |  | lh\_PoG\_L\_4\_1 | Volume for lh\_PoG\_L\_4\_1 | `lh_PoG_L_4_1` |  |
| \|\| |  |  | lh\_PoG\_L\_4\_2 | Volume for lh\_PoG\_L\_4\_2 | `lh_PoG_L_4_2` |  |
| \|\| |  |  | lh\_PoG\_L\_4\_3 | Volume for lh\_PoG\_L\_4\_3 | `lh_PoG_L_4_3` |  |
| \|\| |  |  | lh\_PoG\_L\_4\_4 | Volume for lh\_PoG\_L\_4\_4 | `lh_PoG_L_4_4` |  |
| \|\| |  |  | lh\_INS\_L\_6\_1 | Volume for lh\_INS\_L\_6\_1 | `lh_INS_L_6_1` |  |
| \|\| |  |  | lh\_INS\_L\_6\_2 | Volume for lh\_INS\_L\_6\_2 | `lh_INS_L_6_2` |  |
| \|\| |  |  | lh\_INS\_L\_6\_3 | Volume for lh\_INS\_L\_6\_3 | `lh_INS_L_6_3` |  |
| \|\| |  |  | lh\_CG\_L\_5\_1 | Volume for lh\_CG\_L\_5\_1 | `lh_CG_L_5_1` |  |
| \|\| |  |  | lh\_CG\_L\_5\_2 | Volume for lh\_CG\_L\_5\_2 | `lh_CG_L_5_2` |  |
| \|\| |  |  | lh\_CG\_L\_5\_3 | Volume for lh\_CG\_L\_5\_3 | `lh_CG_L_5_3` |  |
| \|\| |  |  | lh\_CG\_L\_5\_4 | Volume for lh\_CG\_L\_5\_4 | `lh_CG_L_5_4` |  |
| \|\| |  |  | lh\_CG\_L\_5\_5 | Volume for lh\_CG\_L\_5\_5 | `lh_CG_L_5_5` |  |
| \|\| |  |  | lh\_MVOcC\_L\_5\_1 | Volume for lh\_MVOcC\_L\_5\_1 | `lh_MVOcC_L_5_1` |  |
| \|\| |  |  | lh\_MVOcC\_L\_5\_2 | Volume for lh\_MVOcC\_L\_5\_2 | `lh_MVOcC_L_5_2` |  |
| \|\| |  |  | lh\_MVOcC\_L\_5\_3 | Volume for lh\_MVOcC\_L\_5\_3 | `lh_MVOcC_L_5_3` |  |
| \|\| |  |  | lh\_MVOcC\_L\_5\_4 | Volume for lh\_MVOcC\_L\_5\_4 | `lh_MVOcC_L_5_4` |  |
| \|\| |  |  | lh\_MVOcC\_L\_5\_5 | Volume for lh\_MVOcC\_L\_5\_5 | `lh_MVOcC_L_5_5` |  |
| \|\| |  |  | lh\_LOcC\_L\_4\_1 | Volume for lh\_LOcC\_L\_4\_1 | `lh_LOcC_L_4_1` |  |
| \|\| |  |  | lh\_LOcC\_L\_4\_2 | Volume for lh\_LOcC\_L\_4\_2 | `lh_LOcC_L_4_2` |  |
| \|\| |  |  | lh\_LOcC\_L\_4\_3 | Volume for lh\_LOcC\_L\_4\_3 | `lh_LOcC_L_4_3` |  |
| \|\| |  |  | lh\_LOcC\_L\_4\_4 | Volume for lh\_LOcC\_L\_4\_4 | `lh_LOcC_L_4_4` |  |

Close

##### Right Hemisphere Volume Distribution 10

Distribution of cortical region volumes in the right hemisphere across all samples. You may look for extreme outliers, which could indicate segmentation issues or data quality problems. Automatic detection of these outliers is based on the interquartile range (IQR) method, where values falling outside the range defined by Q1 - 3*IQR to Q3 + 3*IQR are considered outliers. Combined with other indicators, these outliers may help identify subjects that require further investigation or exclusion.

###### AI Summary

Provider: , model:

Chat with Seqera AI

Table
 Export...

Copy prompt


Summarize plot

Created with MultiQC

Copy table

 Configure columns

 Sort by highlight

 Scatter plot

 Violin plot
Export as CSV...
Showing ^10^/_10_ rows and ^94^/_94_ columns.

Copy Prompt


Summarize table

| Sample Name | rh\_SFG\_R\_6\_1 | rh\_SFG\_R\_6\_2 | rh\_SFG\_R\_6\_3 | rh\_SFG\_R\_6\_4 | rh\_SFG\_R\_6\_5 | rh\_SFG\_R\_6\_6 | rh\_MFG\_R\_7\_1 | rh\_MFG\_R\_7\_2 | rh\_MFG\_R\_7\_3 | rh\_MFG\_R\_7\_4 | rh\_MFG\_R\_7\_5 | rh\_MFG\_R\_7\_6 | rh\_MFG\_R\_7\_7 | rh\_IFG\_R\_6\_1 | rh\_IFG\_R\_6\_2 | rh\_IFG\_R\_6\_3 | rh\_IFG\_R\_6\_4 | rh\_IFG\_R\_6\_5 | rh\_IFG\_R\_6\_6 | rh\_OrG\_R\_6\_1 | rh\_OrG\_R\_6\_2 | rh\_OrG\_R\_6\_3 | rh\_OrG\_R\_6\_4 | rh\_OrG\_R\_6\_5 | rh\_OrG\_R\_6\_6 | rh\_PrG\_R\_6\_1 | rh\_PrG\_R\_6\_2 | rh\_PrG\_R\_6\_3 | rh\_PrG\_R\_6\_4 | rh\_PrG\_R\_6\_5 | rh\_PrG\_R\_6\_6 | rh\_PCL\_R\_2\_1 | rh\_PCL\_R\_2\_2 | rh\_STG\_R\_6\_1 | rh\_STG\_R\_6\_2 | rh\_STG\_R\_6\_3 | rh\_STG\_R\_6\_4 | rh\_STG\_R\_6\_5 | rh\_STG\_R\_6\_6 | rh\_MTG\_R\_4\_1 | rh\_MTG\_R\_4\_2 | rh\_MTG\_R\_4\_3 | rh\_MTG\_R\_4\_4 | rh\_ITG\_R\_5\_1 | rh\_ITG\_R\_5\_2 | rh\_ITG\_R\_5\_3 | rh\_ITG\_R\_5\_4 | rh\_ITG\_R\_5\_5 | rh\_FuG\_R\_3\_1 | rh\_FuG\_R\_3\_2 | rh\_FuG\_R\_3\_3 | rh\_PhG\_R\_6\_1 | rh\_PhG\_R\_6\_2 | rh\_PhG\_R\_6\_3 | rh\_PhG\_R\_6\_4 | rh\_PhG\_R\_6\_5 | rh\_PhG\_R\_6\_6 | rh\_pSTS\_R\_2\_1 | rh\_pSTS\_R\_2\_2 | rh\_SPL\_R\_4\_1 | rh\_SPL\_R\_4\_2 | rh\_SPL\_R\_4\_3 | rh\_SPL\_R\_4\_4 | rh\_IPL\_R\_6\_1 | rh\_IPL\_R\_6\_2 | rh\_IPL\_R\_6\_3 | rh\_IPL\_R\_6\_4 | rh\_IPL\_R\_6\_5 | rh\_IPL\_R\_6\_6 | rh\_PCun\_R\_4\_1 | rh\_PCun\_R\_4\_2 | rh\_PCun\_R\_4\_3 | rh\_PCun\_R\_4\_4 | rh\_PoG\_R\_4\_1 | rh\_PoG\_R\_4\_2 | rh\_PoG\_R\_4\_3 | rh\_PoG\_R\_4\_4 | rh\_INS\_R\_6\_1 | rh\_INS\_R\_6\_2 | rh\_INS\_R\_6\_3 | rh\_CG\_R\_5\_1 | rh\_CG\_R\_5\_2 | rh\_CG\_R\_5\_3 | rh\_CG\_R\_5\_4 | rh\_CG\_R\_5\_5 | rh\_MVOcC\_R\_5\_1 | rh\_MVOcC\_R\_5\_2 | rh\_MVOcC\_R\_5\_3 | rh\_MVOcC\_R\_5\_4 | rh\_MVOcC\_R\_5\_5 | rh\_LOcC\_R\_4\_1 | rh\_LOcC\_R\_4\_2 | rh\_LOcC\_R\_4\_3 | rh\_LOcC\_R\_4\_4 |
| --- | --- | --- | --- | --- | --- | --- | --- | --- | --- | --- | --- | --- | --- | --- | --- | --- | --- | --- | --- | --- | --- | --- | --- | --- | --- | --- | --- | --- | --- | --- | --- | --- | --- | --- | --- | --- | --- | --- | --- | --- | --- | --- | --- | --- | --- | --- | --- | --- | --- | --- | --- | --- | --- | --- | --- | --- | --- | --- | --- | --- | --- | --- | --- | --- | --- | --- | --- | --- | --- | --- | --- | --- | --- | --- | --- | --- | --- | --- | --- | --- | --- | --- | --- | --- | --- | --- | --- | --- | --- | --- | --- | --- | --- | --- |
| sub-01\_ses-baseline | 2231.0 | 2406.0 | 4099.0 | 5333.0 | 2374.0 | 1603.0 | 2981.0 | 4984.0 | 1873.0 | 4249.0 | 4687.0 | 4362.0 | 603.0 | 3437.0 | 1475.0 | 1457.0 | 1319.0 | 1278.0 | 1980.0 | 1448.0 | 1816.0 | 1727.0 | 5966.0 | 2613.0 | 1175.0 | 2316.0 | 3112.0 | 1737.0 | 1192.0 | 1965.0 | 1652.0 | 1588.0 | 2399.0 | 936.0 | 2731.0 | 1276.0 | 2891.0 | 2392.0 | 3745.0 | 2608.0 | 4226.0 | 2709.0 | 1363.0 | 4151.0 | 1207.0 | 1098.0 | 2586.0 | 1737.0 | 891.0 | 1037.0 | 3373.0 | 3690.0 | 538.0 | 139.0 | 384.0 | 255.0 | 578.0 | 72.0 | 1026.0 | 2339.0 | 3197.0 | 1463.0 | 1388.0 | 5088.0 | 4119.0 | 4300.0 | 2863.0 | 4373.0 | 3399.0 | 4767.0 | 2506.0 | 1477.0 | 2950.0 | 3534.0 | 3147.0 | 1784.0 | 1387.0 | 1311.0 | 883.0 | 3449.0 | 2879.0 | 3242.0 | 1501.0 | 687.0 | 1629.0 | 1845.0 | 900.0 | 1576.0 | 2136.0 | 1278.0 | 3042.0 | 4288.0 | 2337.0 | 1095.0 |
| sub-02\_ses-baseline | 2268.0 | 3320.0 | 3834.0 | 6823.0 | 2709.0 | 1782.0 | 3622.0 | 5408.0 | 1871.0 | 4672.0 | 6450.0 | 3698.0 | 575.0 | 3849.0 | 1580.0 | 1375.0 | 1548.0 | 1265.0 | 2252.0 | 1585.0 | 2335.0 | 2235.0 | 6766.0 | 2861.0 | 1326.0 | 2726.0 | 3104.0 | 1666.0 | 1237.0 | 1944.0 | 1578.0 | 2275.0 | 2468.0 | 974.0 | 2391.0 | 1788.0 | 3352.0 | 2930.0 | 4170.0 | 2507.0 | 5155.0 | 3112.0 | 1545.0 | 4945.0 | 1900.0 | 1515.0 | 3546.0 | 2539.0 | 1558.0 | 1219.0 | 4138.0 | 4076.0 | 392.0 | 178.0 | 528.0 | 267.0 | 549.0 | 83.0 | 1447.0 | 2242.0 | 3457.0 | 2147.0 | 1496.0 | 4893.0 | 4729.0 | 4301.0 | 2677.0 | 5525.0 | 4413.0 | 4896.0 | 2583.0 | 1845.0 | 2646.0 | 3631.0 | 2894.0 | 1822.0 | 1402.0 | 1231.0 | 780.0 | 4113.0 | 3560.0 | 3802.0 | 885.0 | 798.0 | 1868.0 | 1588.0 | 1613.0 | 1381.0 | 2268.0 | 1584.0 | 3238.0 | 5562.0 | 3032.0 | 1250.0 |
| sub-03\_ses-baseline | 1881.0 | 2287.0 | 4091.0 | 4477.0 | 2031.0 | 1382.0 | 3090.0 | 4662.0 | 1700.0 | 4394.0 | 5002.0 | 3576.0 | 553.0 | 3296.0 | 1213.0 | 1551.0 | 1480.0 | 1161.0 | 2263.0 | 1126.0 | 1341.0 | 1648.0 | 4976.0 | 2483.0 | 962.0 | 2092.0 | 3125.0 | 1711.0 | 1246.0 | 1806.0 | 1396.0 | 1691.0 | 2226.0 | 1147.0 | 1792.0 | 1431.0 | 2730.0 | 2881.0 | 4126.0 | 1796.0 | 4832.0 | 2315.0 | 955.0 | 4604.0 | 1422.0 | 1565.0 | 2321.0 | 1216.0 | 783.0 | 1319.0 | 3758.0 | 2889.0 | 185.0 | 148.0 | 416.0 | 191.0 | 316.0 | 61.0 | 1459.0 | 2587.0 | 3324.0 | 2500.0 | 1513.0 | 4660.0 | 3328.0 | 4948.0 | 3548.0 | 5314.0 | 2934.0 | 4326.0 | 2556.0 | 1676.0 | 2633.0 | 2701.0 | 3405.0 | 1511.0 | 1444.0 | 1292.0 | 489.0 | 3282.0 | 3085.0 | 3620.0 | 1440.0 | 696.0 | 1629.0 | 1903.0 | 1064.0 | 1945.0 | 2165.0 | 1962.0 | 3285.0 | 4152.0 | 2686.0 | 1077.0 |
| sub-04\_ses-baseline | 1546.0 | 2170.0 | 3749.0 | 5446.0 | 2392.0 | 2042.0 | 3573.0 | 4641.0 | 1676.0 | 4292.0 | 4289.0 | 5265.0 | 918.0 | 3629.0 | 1337.0 | 1431.0 | 1286.0 | 1427.0 | 1865.0 | 1251.0 | 1576.0 | 1711.0 | 5324.0 | 2904.0 | 1131.0 | 2139.0 | 3083.0 | 2390.0 | 1698.0 | 2180.0 | 1486.0 | 1931.0 | 2447.0 | 1198.0 | 2885.0 | 1387.0 | 2474.0 | 2479.0 | 4206.0 | 1696.0 | 3519.0 | 2774.0 | 1021.0 | 4119.0 | 1200.0 | 1266.0 | 2663.0 | 1316.0 | 1259.0 | 1752.0 | 2921.0 | 3201.0 | 365.0 | 141.0 | 525.0 | 124.0 | 306.0 | 74.0 | 1139.0 | 2232.0 | 3831.0 | 1660.0 | 1291.0 | 4623.0 | 2860.0 | 4722.0 | 3364.0 | 4853.0 | 3206.0 | 4083.0 | 1955.0 | 1598.0 | 2392.0 | 2964.0 | 3443.0 | 1621.0 | 1937.0 | 1213.0 | 660.0 | 3374.0 | 3090.0 | 2990.0 | 1646.0 | 770.0 | 1606.0 | 1476.0 | 1525.0 | 1509.0 | 2987.0 | 1847.0 | 4046.0 | 4008.0 | 2261.0 | 1233.0 |
| sub-05\_ses-baseline | 2229.0 | 3330.0 | 4430.0 | 6864.0 | 2574.0 | 1992.0 | 3922.0 | 5855.0 | 2211.0 | 4257.0 | 5417.0 | 4818.0 | 1056.0 | 3746.0 | 1470.0 | 1787.0 | 1730.0 | 1242.0 | 2531.0 | 1362.0 | 1801.0 | 2008.0 | 5407.0 | 3358.0 | 1260.0 | 2751.0 | 3436.0 | 2110.0 | 1434.0 | 2030.0 | 1856.0 | 1723.0 | 2210.0 | 1049.0 | 3739.0 | 1823.0 | 2975.0 | 3205.0 | 5336.0 | 2280.0 | 5708.0 | 3845.0 | 1976.0 | 5390.0 | 1924.0 | 1929.0 | 4169.0 | 1880.0 | 1899.0 | 1888.0 | 3959.0 | 4203.0 | 477.0 | 138.0 | 379.0 | 240.0 | 401.0 | 52.0 | 1502.0 | 2315.0 | 4330.0 | 1718.0 | 1403.0 | 4651.0 | 4100.0 | 5599.0 | 3903.0 | 6700.0 | 4399.0 | 6172.0 | 2594.0 | 2174.0 | 4067.0 | 2853.0 | 3858.0 | 2111.0 | 1901.0 | 1401.0 | 967.0 | 4193.0 | 3636.0 | 4535.0 | 1309.0 | 767.0 | 1843.0 | 1864.0 | 1618.0 | 1775.0 | 2777.0 | 2100.0 | 4608.0 | 5788.0 | 3187.0 | 1126.0 |
| sub-06\_ses-baseline | 2236.0 | 3366.0 | 5967.0 | 6269.0 | 3372.0 | 2895.0 | 2787.0 | 5781.0 | 1561.0 | 5126.0 | 5459.0 | 3812.0 | 693.0 | 3546.0 | 1453.0 | 1845.0 | 1250.0 | 1487.0 | 2246.0 | 1452.0 | 1612.0 | 1831.0 | 5681.0 | 2952.0 | 1438.0 | 2231.0 | 3054.0 | 2233.0 | 1576.0 | 2358.0 | 1702.0 | 1906.0 | 2868.0 | 1098.0 | 2949.0 | 1463.0 | 3281.0 | 2792.0 | 4813.0 | 2257.0 | 3507.0 | 3772.0 | 884.0 | 4186.0 | 1722.0 | 2165.0 | 4331.0 | 1589.0 | 1619.0 | 1475.0 | 4232.0 | 4516.0 | 550.0 | 186.0 | 566.0 | 212.0 | 368.0 | 101.0 | 1145.0 | 1983.0 | 4137.0 | 1460.0 | 1587.0 | 4597.0 | 3906.0 | 4161.0 | 3630.0 | 4390.0 | 2767.0 | 5039.0 | 2753.0 | 1800.0 | 3297.0 | 3550.0 | 4056.0 | 1838.0 | 1962.0 | 1442.0 | 852.0 | 4079.0 | 3665.0 | 3706.0 | 1187.0 | 869.0 | 2131.0 | 2345.0 | 1577.0 | 1793.0 | 2151.0 | 2303.0 | 3824.0 | 4839.0 | 3059.0 | 1167.0 |
| sub-07\_ses-baseline | 3134.0 | 3889.0 | 5657.0 | 7799.0 | 3364.0 | 2210.0 | 4247.0 | 6458.0 | 2595.0 | 5504.0 | 7520.0 | 4538.0 | 734.0 | 3789.0 | 1690.0 | 2452.0 | 2233.0 | 1456.0 | 3015.0 | 1481.0 | 2507.0 | 1811.0 | 7601.0 | 2952.0 | 1629.0 | 2689.0 | 3783.0 | 2221.0 | 1648.0 | 2482.0 | 1913.0 | 1793.0 | 2773.0 | 1435.0 | 3509.0 | 1549.0 | 3690.0 | 3161.0 | 5727.0 | 2451.0 | 5352.0 | 3954.0 | 1328.0 | 6202.0 | 1266.0 | 2389.0 | 5046.0 | 1866.0 | 2272.0 | 2333.0 | 3337.0 | 4093.0 | 1164.0 | 169.0 | 779.0 | 247.0 | 441.0 | 86.0 | 1827.0 | 2074.0 | 4198.0 | 1496.0 | 2226.0 | 6077.0 | 5137.0 | 5715.0 | 4258.0 | 7229.0 | 4100.0 | 6902.0 | 3226.0 | 2160.0 | 3925.0 | 3484.0 | 4896.0 | 2352.0 | 3636.0 | 1920.0 | 1157.0 | 4810.0 | 4453.0 | 3890.0 | 2207.0 | 855.0 | 2305.0 | 2720.0 | 976.0 | 1902.0 | 2508.0 | 2113.0 | 4698.0 | 5825.0 | 2340.0 | 1482.0 |
| sub-08\_ses-baseline | 2555.0 | 2961.0 | 5165.0 | 7272.0 | 2739.0 | 1820.0 | 2737.0 | 5654.0 | 1970.0 | 4557.0 | 5429.0 | 4689.0 | 716.0 | 3456.0 | 1955.0 | 1612.0 | 1372.0 | 1415.0 | 1865.0 | 1393.0 | 1765.0 | 1783.0 | 6713.0 | 2895.0 | 1460.0 | 2680.0 | 3811.0 | 1793.0 | 1406.0 | 2148.0 | 1562.0 | 1998.0 | 2721.0 | 1053.0 | 3294.0 | 1523.0 | 3060.0 | 2635.0 | 5158.0 | 2137.0 | 4684.0 | 3755.0 | 1562.0 | 5249.0 | 1743.0 | 1904.0 | 4545.0 | 2369.0 | 1641.0 | 1543.0 | 4270.0 | 4445.0 | 821.0 | 185.0 | 696.0 | 219.0 | 525.0 | 103.0 | 1403.0 | 2412.0 | 3666.0 | 1650.0 | 1763.0 | 5212.0 | 4378.0 | 4643.0 | 3568.0 | 4800.0 | 3816.0 | 5050.0 | 2759.0 | 1907.0 | 3394.0 | 3964.0 | 3977.0 | 1818.0 | 2414.0 | 1350.0 | 1004.0 | 4018.0 | 3140.0 | 3948.0 | 1396.0 | 939.0 | 1702.0 | 2377.0 | 1170.0 | 1779.0 | 2908.0 | 2272.0 | 3889.0 | 5315.0 | 2907.0 | 1179.0 |
| sub-09\_ses-baseline | 1549.0 | 2411.0 | 3531.0 | 4909.0 | 2434.0 | 1760.0 | 2749.0 | 5158.0 | 1521.0 | 4169.0 | 4435.0 | 3231.0 | 550.0 | 3275.0 | 1105.0 | 994.0 | 1150.0 | 966.0 | 1617.0 | 1220.0 | 1666.0 | 1836.0 | 5015.0 | 2337.0 | 1104.0 | 2120.0 | 2598.0 | 1327.0 | 999.0 | 1841.0 | 1357.0 | 1140.0 | 2126.0 | 945.0 | 2408.0 | 1406.0 | 2780.0 | 2311.0 | 4947.0 | 1768.0 | 3412.0 | 2573.0 | 692.0 | 3673.0 | 1359.0 | 1220.0 | 3042.0 | 1297.0 | 1357.0 | 1245.0 | 3302.0 | 3693.0 | 327.0 | 167.0 | 398.0 | 159.0 | 214.0 | 64.0 | 1003.0 | 2300.0 | 3334.0 | 1062.0 | 1247.0 | 3571.0 | 2742.0 | 3482.0 | 2391.0 | 3721.0 | 2694.0 | 4195.0 | 1807.0 | 1593.0 | 2104.0 | 2314.0 | 3199.0 | 1486.0 | 1660.0 | 1363.0 | 295.0 | 2968.0 | 2688.0 | 2481.0 | 822.0 | 656.0 | 1570.0 | 1401.0 | 994.0 | 1356.0 | 2240.0 | 1383.0 | 2970.0 | 4490.0 | 2834.0 | 1087.0 |
| sub-11\_ses-baseline | 3085.0 | 3527.0 | 5878.0 | 7633.0 | 3936.0 | 2380.0 | 5032.0 | 7224.0 | 1788.0 | 6052.0 | 6473.0 | 4628.0 | 622.0 | 4377.0 | 1311.0 | 2242.0 | 1436.0 | 2004.0 | 2418.0 | 1594.0 | 2453.0 | 2494.0 | 6520.0 | 3039.0 | 1477.0 | 2989.0 | 3116.0 | 2361.0 | 1538.0 | 2312.0 | 1696.0 | 1385.0 | 3210.0 | 1156.0 | 2847.0 | 1654.0 | 3030.0 | 3120.0 | 5237.0 | 2452.0 | 4914.0 | 3519.0 | 2064.0 | 4465.0 | 2477.0 | 1614.0 | 3647.0 | 2134.0 | 1493.0 | 1782.0 | 4426.0 | 3873.0 | 184.0 | 143.0 | 504.0 | 160.0 | 264.0 | 106.0 | 1415.0 | 1778.0 | 4659.0 | 2109.0 | 1785.0 | 5304.0 | 4859.0 | 4702.0 | 3399.0 | 4221.0 | 4139.0 | 4547.0 | 2341.0 | 1646.0 | 3179.0 | 3411.0 | 3808.0 | 1780.0 | 2021.0 | 1389.0 | 1266.0 | 4029.0 | 3982.0 | 4372.0 | 1735.0 | 1129.0 | 2239.0 | 2621.0 | 1458.0 | 2197.0 | 3911.0 | 2067.0 | 5337.0 | 5856.0 | 3422.0 | 1703.0 |

###### Cortical Regions: Right Hemisphere Volume Distribution: Columns

Uncheck the tick box to hide columns. Click and drag the handle on the left to change order. Table ID: `cortical_rh_volume_plot_table`

Show All
Show None

| Sort | Visible | Group | Column | Description | ID | Scale |
| --- | --- | --- | --- | --- | --- | --- |
| \|\| |  |  | rh\_SFG\_R\_6\_1 | Volume for rh\_SFG\_R\_6\_1 | `rh_SFG_R_6_1` |  |
| \|\| |  |  | rh\_SFG\_R\_6\_2 | Volume for rh\_SFG\_R\_6\_2 | `rh_SFG_R_6_2` |  |
| \|\| |  |  | rh\_SFG\_R\_6\_3 | Volume for rh\_SFG\_R\_6\_3 | `rh_SFG_R_6_3` |  |
| \|\| |  |  | rh\_SFG\_R\_6\_4 | Volume for rh\_SFG\_R\_6\_4 | `rh_SFG_R_6_4` |  |
| \|\| |  |  | rh\_SFG\_R\_6\_5 | Volume for rh\_SFG\_R\_6\_5 | `rh_SFG_R_6_5` |  |
| \|\| |  |  | rh\_SFG\_R\_6\_6 | Volume for rh\_SFG\_R\_6\_6 | `rh_SFG_R_6_6` |  |
| \|\| |  |  | rh\_MFG\_R\_7\_1 | Volume for rh\_MFG\_R\_7\_1 | `rh_MFG_R_7_1` |  |
| \|\| |  |  | rh\_MFG\_R\_7\_2 | Volume for rh\_MFG\_R\_7\_2 | `rh_MFG_R_7_2` |  |
| \|\| |  |  | rh\_MFG\_R\_7\_3 | Volume for rh\_MFG\_R\_7\_3 | `rh_MFG_R_7_3` |  |
| \|\| |  |  | rh\_MFG\_R\_7\_4 | Volume for rh\_MFG\_R\_7\_4 | `rh_MFG_R_7_4` |  |
| \|\| |  |  | rh\_MFG\_R\_7\_5 | Volume for rh\_MFG\_R\_7\_5 | `rh_MFG_R_7_5` |  |
| \|\| |  |  | rh\_MFG\_R\_7\_6 | Volume for rh\_MFG\_R\_7\_6 | `rh_MFG_R_7_6` |  |
| \|\| |  |  | rh\_MFG\_R\_7\_7 | Volume for rh\_MFG\_R\_7\_7 | `rh_MFG_R_7_7` |  |
| \|\| |  |  | rh\_IFG\_R\_6\_1 | Volume for rh\_IFG\_R\_6\_1 | `rh_IFG_R_6_1` |  |
| \|\| |  |  | rh\_IFG\_R\_6\_2 | Volume for rh\_IFG\_R\_6\_2 | `rh_IFG_R_6_2` |  |
| \|\| |  |  | rh\_IFG\_R\_6\_3 | Volume for rh\_IFG\_R\_6\_3 | `rh_IFG_R_6_3` |  |
| \|\| |  |  | rh\_IFG\_R\_6\_4 | Volume for rh\_IFG\_R\_6\_4 | `rh_IFG_R_6_4` |  |
| \|\| |  |  | rh\_IFG\_R\_6\_5 | Volume for rh\_IFG\_R\_6\_5 | `rh_IFG_R_6_5` |  |
| \|\| |  |  | rh\_IFG\_R\_6\_6 | Volume for rh\_IFG\_R\_6\_6 | `rh_IFG_R_6_6` |  |
| \|\| |  |  | rh\_OrG\_R\_6\_1 | Volume for rh\_OrG\_R\_6\_1 | `rh_OrG_R_6_1` |  |
| \|\| |  |  | rh\_OrG\_R\_6\_2 | Volume for rh\_OrG\_R\_6\_2 | `rh_OrG_R_6_2` |  |
| \|\| |  |  | rh\_OrG\_R\_6\_3 | Volume for rh\_OrG\_R\_6\_3 | `rh_OrG_R_6_3` |  |
| \|\| |  |  | rh\_OrG\_R\_6\_4 | Volume for rh\_OrG\_R\_6\_4 | `rh_OrG_R_6_4` |  |
| \|\| |  |  | rh\_OrG\_R\_6\_5 | Volume for rh\_OrG\_R\_6\_5 | `rh_OrG_R_6_5` |  |
| \|\| |  |  | rh\_OrG\_R\_6\_6 | Volume for rh\_OrG\_R\_6\_6 | `rh_OrG_R_6_6` |  |
| \|\| |  |  | rh\_PrG\_R\_6\_1 | Volume for rh\_PrG\_R\_6\_1 | `rh_PrG_R_6_1` |  |
| \|\| |  |  | rh\_PrG\_R\_6\_2 | Volume for rh\_PrG\_R\_6\_2 | `rh_PrG_R_6_2` |  |
| \|\| |  |  | rh\_PrG\_R\_6\_3 | Volume for rh\_PrG\_R\_6\_3 | `rh_PrG_R_6_3` |  |
| \|\| |  |  | rh\_PrG\_R\_6\_4 | Volume for rh\_PrG\_R\_6\_4 | `rh_PrG_R_6_4` |  |
| \|\| |  |  | rh\_PrG\_R\_6\_5 | Volume for rh\_PrG\_R\_6\_5 | `rh_PrG_R_6_5` |  |
| \|\| |  |  | rh\_PrG\_R\_6\_6 | Volume for rh\_PrG\_R\_6\_6 | `rh_PrG_R_6_6` |  |
| \|\| |  |  | rh\_PCL\_R\_2\_1 | Volume for rh\_PCL\_R\_2\_1 | `rh_PCL_R_2_1` |  |
| \|\| |  |  | rh\_PCL\_R\_2\_2 | Volume for rh\_PCL\_R\_2\_2 | `rh_PCL_R_2_2` |  |
| \|\| |  |  | rh\_STG\_R\_6\_1 | Volume for rh\_STG\_R\_6\_1 | `rh_STG_R_6_1` |  |
| \|\| |  |  | rh\_STG\_R\_6\_2 | Volume for rh\_STG\_R\_6\_2 | `rh_STG_R_6_2` |  |
| \|\| |  |  | rh\_STG\_R\_6\_3 | Volume for rh\_STG\_R\_6\_3 | `rh_STG_R_6_3` |  |
| \|\| |  |  | rh\_STG\_R\_6\_4 | Volume for rh\_STG\_R\_6\_4 | `rh_STG_R_6_4` |  |
| \|\| |  |  | rh\_STG\_R\_6\_5 | Volume for rh\_STG\_R\_6\_5 | `rh_STG_R_6_5` |  |
| \|\| |  |  | rh\_STG\_R\_6\_6 | Volume for rh\_STG\_R\_6\_6 | `rh_STG_R_6_6` |  |
| \|\| |  |  | rh\_MTG\_R\_4\_1 | Volume for rh\_MTG\_R\_4\_1 | `rh_MTG_R_4_1` |  |
| \|\| |  |  | rh\_MTG\_R\_4\_2 | Volume for rh\_MTG\_R\_4\_2 | `rh_MTG_R_4_2` |  |
| \|\| |  |  | rh\_MTG\_R\_4\_3 | Volume for rh\_MTG\_R\_4\_3 | `rh_MTG_R_4_3` |  |
| \|\| |  |  | rh\_MTG\_R\_4\_4 | Volume for rh\_MTG\_R\_4\_4 | `rh_MTG_R_4_4` |  |
| \|\| |  |  | rh\_ITG\_R\_5\_1 | Volume for rh\_ITG\_R\_5\_1 | `rh_ITG_R_5_1` |  |
| \|\| |  |  | rh\_ITG\_R\_5\_2 | Volume for rh\_ITG\_R\_5\_2 | `rh_ITG_R_5_2` |  |
| \|\| |  |  | rh\_ITG\_R\_5\_3 | Volume for rh\_ITG\_R\_5\_3 | `rh_ITG_R_5_3` |  |
| \|\| |  |  | rh\_ITG\_R\_5\_4 | Volume for rh\_ITG\_R\_5\_4 | `rh_ITG_R_5_4` |  |
| \|\| |  |  | rh\_ITG\_R\_5\_5 | Volume for rh\_ITG\_R\_5\_5 | `rh_ITG_R_5_5` |  |
| \|\| |  |  | rh\_FuG\_R\_3\_1 | Volume for rh\_FuG\_R\_3\_1 | `rh_FuG_R_3_1` |  |
| \|\| |  |  | rh\_FuG\_R\_3\_2 | Volume for rh\_FuG\_R\_3\_2 | `rh_FuG_R_3_2` |  |
| \|\| |  |  | rh\_FuG\_R\_3\_3 | Volume for rh\_FuG\_R\_3\_3 | `rh_FuG_R_3_3` |  |
| \|\| |  |  | rh\_PhG\_R\_6\_1 | Volume for rh\_PhG\_R\_6\_1 | `rh_PhG_R_6_1` |  |
| \|\| |  |  | rh\_PhG\_R\_6\_2 | Volume for rh\_PhG\_R\_6\_2 | `rh_PhG_R_6_2` |  |
| \|\| |  |  | rh\_PhG\_R\_6\_3 | Volume for rh\_PhG\_R\_6\_3 | `rh_PhG_R_6_3` |  |
| \|\| |  |  | rh\_PhG\_R\_6\_4 | Volume for rh\_PhG\_R\_6\_4 | `rh_PhG_R_6_4` |  |
| \|\| |  |  | rh\_PhG\_R\_6\_5 | Volume for rh\_PhG\_R\_6\_5 | `rh_PhG_R_6_5` |  |
| \|\| |  |  | rh\_PhG\_R\_6\_6 | Volume for rh\_PhG\_R\_6\_6 | `rh_PhG_R_6_6` |  |
| \|\| |  |  | rh\_pSTS\_R\_2\_1 | Volume for rh\_pSTS\_R\_2\_1 | `rh_pSTS_R_2_1` |  |
| \|\| |  |  | rh\_pSTS\_R\_2\_2 | Volume for rh\_pSTS\_R\_2\_2 | `rh_pSTS_R_2_2` |  |
| \|\| |  |  | rh\_SPL\_R\_4\_1 | Volume for rh\_SPL\_R\_4\_1 | `rh_SPL_R_4_1` |  |
| \|\| |  |  | rh\_SPL\_R\_4\_2 | Volume for rh\_SPL\_R\_4\_2 | `rh_SPL_R_4_2` |  |
| \|\| |  |  | rh\_SPL\_R\_4\_3 | Volume for rh\_SPL\_R\_4\_3 | `rh_SPL_R_4_3` |  |
| \|\| |  |  | rh\_SPL\_R\_4\_4 | Volume for rh\_SPL\_R\_4\_4 | `rh_SPL_R_4_4` |  |
| \|\| |  |  | rh\_IPL\_R\_6\_1 | Volume for rh\_IPL\_R\_6\_1 | `rh_IPL_R_6_1` |  |
| \|\| |  |  | rh\_IPL\_R\_6\_2 | Volume for rh\_IPL\_R\_6\_2 | `rh_IPL_R_6_2` |  |
| \|\| |  |  | rh\_IPL\_R\_6\_3 | Volume for rh\_IPL\_R\_6\_3 | `rh_IPL_R_6_3` |  |
| \|\| |  |  | rh\_IPL\_R\_6\_4 | Volume for rh\_IPL\_R\_6\_4 | `rh_IPL_R_6_4` |  |
| \|\| |  |  | rh\_IPL\_R\_6\_5 | Volume for rh\_IPL\_R\_6\_5 | `rh_IPL_R_6_5` |  |
| \|\| |  |  | rh\_IPL\_R\_6\_6 | Volume for rh\_IPL\_R\_6\_6 | `rh_IPL_R_6_6` |  |
| \|\| |  |  | rh\_PCun\_R\_4\_1 | Volume for rh\_PCun\_R\_4\_1 | `rh_PCun_R_4_1` |  |
| \|\| |  |  | rh\_PCun\_R\_4\_2 | Volume for rh\_PCun\_R\_4\_2 | `rh_PCun_R_4_2` |  |
| \|\| |  |  | rh\_PCun\_R\_4\_3 | Volume for rh\_PCun\_R\_4\_3 | `rh_PCun_R_4_3` |  |
| \|\| |  |  | rh\_PCun\_R\_4\_4 | Volume for rh\_PCun\_R\_4\_4 | `rh_PCun_R_4_4` |  |
| \|\| |  |  | rh\_PoG\_R\_4\_1 | Volume for rh\_PoG\_R\_4\_1 | `rh_PoG_R_4_1` |  |
| \|\| |  |  | rh\_PoG\_R\_4\_2 | Volume for rh\_PoG\_R\_4\_2 | `rh_PoG_R_4_2` |  |
| \|\| |  |  | rh\_PoG\_R\_4\_3 | Volume for rh\_PoG\_R\_4\_3 | `rh_PoG_R_4_3` |  |
| \|\| |  |  | rh\_PoG\_R\_4\_4 | Volume for rh\_PoG\_R\_4\_4 | `rh_PoG_R_4_4` |  |
| \|\| |  |  | rh\_INS\_R\_6\_1 | Volume for rh\_INS\_R\_6\_1 | `rh_INS_R_6_1` |  |
| \|\| |  |  | rh\_INS\_R\_6\_2 | Volume for rh\_INS\_R\_6\_2 | `rh_INS_R_6_2` |  |
| \|\| |  |  | rh\_INS\_R\_6\_3 | Volume for rh\_INS\_R\_6\_3 | `rh_INS_R_6_3` |  |
| \|\| |  |  | rh\_CG\_R\_5\_1 | Volume for rh\_CG\_R\_5\_1 | `rh_CG_R_5_1` |  |
| \|\| |  |  | rh\_CG\_R\_5\_2 | Volume for rh\_CG\_R\_5\_2 | `rh_CG_R_5_2` |  |
| \|\| |  |  | rh\_CG\_R\_5\_3 | Volume for rh\_CG\_R\_5\_3 | `rh_CG_R_5_3` |  |
| \|\| |  |  | rh\_CG\_R\_5\_4 | Volume for rh\_CG\_R\_5\_4 | `rh_CG_R_5_4` |  |
| \|\| |  |  | rh\_CG\_R\_5\_5 | Volume for rh\_CG\_R\_5\_5 | `rh_CG_R_5_5` |  |
| \|\| |  |  | rh\_MVOcC\_R\_5\_1 | Volume for rh\_MVOcC\_R\_5\_1 | `rh_MVOcC_R_5_1` |  |
| \|\| |  |  | rh\_MVOcC\_R\_5\_2 | Volume for rh\_MVOcC\_R\_5\_2 | `rh_MVOcC_R_5_2` |  |
| \|\| |  |  | rh\_MVOcC\_R\_5\_3 | Volume for rh\_MVOcC\_R\_5\_3 | `rh_MVOcC_R_5_3` |  |
| \|\| |  |  | rh\_MVOcC\_R\_5\_4 | Volume for rh\_MVOcC\_R\_5\_4 | `rh_MVOcC_R_5_4` |  |
| \|\| |  |  | rh\_MVOcC\_R\_5\_5 | Volume for rh\_MVOcC\_R\_5\_5 | `rh_MVOcC_R_5_5` |  |
| \|\| |  |  | rh\_LOcC\_R\_4\_1 | Volume for rh\_LOcC\_R\_4\_1 | `rh_LOcC_R_4_1` |  |
| \|\| |  |  | rh\_LOcC\_R\_4\_2 | Volume for rh\_LOcC\_R\_4\_2 | `rh_LOcC_R_4_2` |  |
| \|\| |  |  | rh\_LOcC\_R\_4\_3 | Volume for rh\_LOcC\_R\_4\_3 | `rh_LOcC_R_4_3` |  |
| \|\| |  |  | rh\_LOcC\_R\_4\_4 | Volume for rh\_LOcC\_R\_4\_4 | `rh_LOcC_R_4_4` |  |

Close

#### Subcortical Regions

Assessment of subcortical region volumes for quality control using IQR-based outlier detection. Each subcortical region's volume is evaluated across subjects, and regions with volumes falling outside the range are considered outliers. The percentage of outlier regions per subject is reported in the general statistics, with thresholds for pass/warn/fail configurable in the MultiQC configuration file.https://github.com/nf-neuro/MultiQC\_neuroimaging

##### Subcortical Volume Distribution 10

Distribution of subcortical region volumes across all samples. You may look for extreme outliers, which could indicate segmentation issues or data quality problems. Automatic outlier detection is based on volumes falling outside the range defined by Q1 - 3*IQR to Q3 + 3*IQR. Combined with other indicators, these outliers may help identify subjects that require further investigation or exclusion.

###### AI Summary

Provider: , model:

Chat with Seqera AI

Table
 Export...

Copy prompt


Summarize plot

Created with MultiQC

Copy table

 Configure columns

 Sort by highlight

 Scatter plot

 Violin plot
Export as CSV...
Showing ^10^/_10_ rows and ^39^/_39_ columns.

Copy Prompt


Summarize table

| Sample Name | mAmyg\_L | mAmyg\_R | lAmyg\_L | lAmyg\_R | rHipp\_L | rHipp\_R | cHipp\_L | cHipp\_R | vCa\_L | vCa\_R | GP\_L | GP\_R | NAC\_L | NAC\_R | vmPu\_L | vmPu\_R | dCa\_L | dCa\_R | dlPu\_L | dlPu\_R | mPFtha\_L | mPFtha\_R | mPMtha\_L | mPMtha\_R | Stha\_L | Stha\_R | rTtha\_L | rTtha\_R | PPtha\_L | PPtha\_R | Otha\_L | Otha\_R | cTtha\_L | cTtha\_R | lPFtha\_L | lPFtha\_R | brainstem | left\_cerebellum\_cortex | right\_cerebellum\_cortex |
| --- | --- | --- | --- | --- | --- | --- | --- | --- | --- | --- | --- | --- | --- | --- | --- | --- | --- | --- | --- | --- | --- | --- | --- | --- | --- | --- | --- | --- | --- | --- | --- | --- | --- | --- | --- | --- | --- | --- | --- |
| sub-01\_ses-baseline | 1033.2 | 1345.1 | 363.7 | 645.8 | 3200.4 | 2803.8 | 2972.9 | 3075.9 | 2246.3 | 1642.1 | 2107.0 | 2191.7 | 1368.6 | 1304.2 | 1891.2 | 1581.7 | 2755.7 | 3543.2 | 2874.5 | 2672.1 | 1266.4 | 1221.5 | 718.2 | 1645.5 | 918.3 | 1080.2 | 1196.0 | 873.2 | 1495.9 | 1158.1 | 1163.5 | 741.2 | 1070.5 | 814.5 | 2444.4 | 1398.9 | 22122.0 | 59639.4 | 60262.4 |
| sub-02\_ses-baseline | 1053.2 | 1415.0 | 439.1 | 588.3 | 3478.7 | 2849.7 | 3054.4 | 3151.1 | 2469.7 | 1897.6 | 2116.2 | 2271.3 | 1245.7 | 1575.0 | 1998.5 | 1697.8 | 3027.5 | 3633.9 | 3166.4 | 2788.4 | 1165.6 | 1259.1 | 772.9 | 1738.6 | 939.3 | 1025.3 | 1322.1 | 784.0 | 1443.8 | 1094.3 | 1160.2 | 677.4 | 1187.6 | 934.1 | 2469.5 | 1490.4 | 20281.6 | 65303.0 | 62762.5 |
| sub-03\_ses-baseline | 1014.1 | 1276.7 | 438.0 | 551.8 | 3329.0 | 2619.9 | 2672.7 | 2756.6 | 2168.2 | 1625.2 | 1843.2 | 2129.5 | 1267.5 | 1360.3 | 1789.2 | 1594.0 | 2907.1 | 3441.7 | 2859.3 | 2648.5 | 1015.2 | 1151.7 | 750.1 | 1658.6 | 813.4 | 1009.3 | 1207.6 | 798.0 | 1377.0 | 1009.7 | 963.2 | 605.8 | 1079.7 | 845.8 | 2164.1 | 1373.5 | 17841.9 | 59294.2 | 55635.8 |
| sub-04\_ses-baseline | 888.9 | 1190.5 | 437.8 | 630.0 | 3149.1 | 2733.0 | 2888.1 | 2765.8 | 1900.6 | 1496.4 | 2022.8 | 2160.0 | 1268.8 | 1068.0 | 1718.0 | 1418.0 | 2879.5 | 3275.7 | 2543.8 | 2356.9 | 1032.6 | 1162.0 | 636.3 | 1635.8 | 790.5 | 985.1 | 1115.8 | 784.5 | 1331.1 | 954.6 | 948.4 | 669.7 | 1084.7 | 815.7 | 2230.6 | 1326.9 | 17627.2 | 55672.4 | 53263.0 |
| sub-05\_ses-baseline | 996.7 | 1435.1 | 507.7 | 821.6 | 3708.4 | 3164.6 | 2949.0 | 3500.1 | 2453.8 | 1782.3 | 2397.2 | 2491.9 | 1391.5 | 1600.9 | 2123.4 | 1749.7 | 3454.5 | 4004.1 | 3100.4 | 3135.0 | 1274.9 | 1286.6 | 777.0 | 1805.7 | 936.2 | 1154.0 | 1218.8 | 867.3 | 1604.7 | 1176.9 | 1328.5 | 747.1 | 1229.1 | 946.3 | 2700.3 | 1622.2 | 22726.9 | 59247.9 | 59541.6 |
| sub-06\_ses-baseline | 1028.9 | 1481.3 | 463.1 | 639.9 | 3622.3 | 2971.0 | 3041.2 | 3230.6 | 2288.9 | 1930.6 | 2309.6 | 2333.2 | 1496.6 | 1496.6 | 2048.7 | 1801.0 | 3282.3 | 3614.8 | 2973.9 | 3082.8 | 1238.3 | 1326.3 | 804.2 | 1782.0 | 893.0 | 1263.4 | 1228.6 | 943.4 | 1487.6 | 1119.0 | 1104.1 | 812.8 | 1314.4 | 896.8 | 2482.1 | 1485.6 | 22835.2 | 64406.6 | 61946.1 |
| sub-07\_ses-baseline | 1091.2 | 1554.6 | 606.4 | 811.8 | 4384.8 | 3651.8 | 3308.7 | 3983.5 | 2622.5 | 2101.0 | 2433.2 | 2684.0 | 1780.0 | 1630.5 | 2256.2 | 1963.1 | 3646.0 | 4157.0 | 3393.6 | 3630.2 | 1604.6 | 1235.2 | 895.6 | 1866.7 | 1097.0 | 1199.5 | 1599.2 | 1122.1 | 1770.6 | 1249.2 | 1399.0 | 780.5 | 1293.4 | 945.5 | 3095.8 | 1771.7 | 24138.1 | 66847.1 | 65710.0 |
| sub-08\_ses-baseline | 1064.4 | 1599.1 | 595.7 | 687.4 | 3815.3 | 3276.8 | 3437.3 | 3545.0 | 2460.1 | 1966.8 | 2314.0 | 2454.7 | 1442.6 | 1408.5 | 2005.2 | 1661.8 | 3356.7 | 4178.9 | 3121.5 | 3180.5 | 1255.2 | 1302.9 | 841.0 | 1800.7 | 1007.3 | 1185.2 | 1450.4 | 940.3 | 1632.2 | 1221.4 | 1213.9 | 811.9 | 1204.2 | 943.9 | 2599.4 | 1558.3 | 22610.2 | 70204.1 | 70426.1 |
| sub-09\_ses-baseline | 861.2 | 1126.8 | 463.9 | 598.3 | 2657.2 | 2266.3 | 2754.3 | 2509.9 | 1761.3 | 1386.2 | 1844.4 | 1876.9 | 1124.9 | 1126.9 | 1613.6 | 1396.6 | 2528.8 | 2938.2 | 2534.0 | 2380.3 | 888.7 | 968.1 | 676.1 | 1304.4 | 678.6 | 750.5 | 1242.5 | 750.1 | 1208.6 | 937.6 | 1063.3 | 575.4 | 742.4 | 728.4 | 1785.5 | 1277.8 | 20057.1 | 57402.4 | 56781.3 |
| sub-11\_ses-baseline | 1090.3 | 1507.8 | 554.7 | 754.5 | 3778.9 | 3174.1 | 3171.0 | 3355.7 | 2446.4 | 1904.9 | 2201.3 | 2586.2 | 1528.6 | 1685.8 | 2047.3 | 1748.9 | 3188.4 | 3622.3 | 3296.5 | 2824.7 | 1269.2 | 1302.1 | 823.7 | 1860.2 | 931.4 | 1147.1 | 1510.2 | 908.2 | 1549.4 | 1249.2 | 1314.1 | 696.8 | 1245.0 | 1074.4 | 2592.0 | 1705.4 | 20732.8 | 63245.4 | 60278.8 |

###### Subcortical Regions: Volume Distribution: Columns

Uncheck the tick box to hide columns. Click and drag the handle on the left to change order. Table ID: `subcortical_volume_plot_table`

Show All
Show None

| Sort | Visible | Group | Column | Description | ID | Scale |
| --- | --- | --- | --- | --- | --- | --- |
| \|\| |  |  | mAmyg\_L | Volume for mAmyg\_L | `mAmyg_L` |  |
| \|\| |  |  | mAmyg\_R | Volume for mAmyg\_R | `mAmyg_R` |  |
| \|\| |  |  | lAmyg\_L | Volume for lAmyg\_L | `lAmyg_L` |  |
| \|\| |  |  | lAmyg\_R | Volume for lAmyg\_R | `lAmyg_R` |  |
| \|\| |  |  | rHipp\_L | Volume for rHipp\_L | `rHipp_L` |  |
| \|\| |  |  | rHipp\_R | Volume for rHipp\_R | `rHipp_R` |  |
| \|\| |  |  | cHipp\_L | Volume for cHipp\_L | `cHipp_L` |  |
| \|\| |  |  | cHipp\_R | Volume for cHipp\_R | `cHipp_R` |  |
| \|\| |  |  | vCa\_L | Volume for vCa\_L | `vCa_L` |  |
| \|\| |  |  | vCa\_R | Volume for vCa\_R | `vCa_R` |  |
| \|\| |  |  | GP\_L | Volume for GP\_L | `GP_L` |  |
| \|\| |  |  | GP\_R | Volume for GP\_R | `GP_R` |  |
| \|\| |  |  | NAC\_L | Volume for NAC\_L | `NAC_L` |  |
| \|\| |  |  | NAC\_R | Volume for NAC\_R | `NAC_R` |  |
| \|\| |  |  | vmPu\_L | Volume for vmPu\_L | `vmPu_L` |  |
| \|\| |  |  | vmPu\_R | Volume for vmPu\_R | `vmPu_R` |  |
| \|\| |  |  | dCa\_L | Volume for dCa\_L | `dCa_L` |  |
| \|\| |  |  | dCa\_R | Volume for dCa\_R | `dCa_R` |  |
| \|\| |  |  | dlPu\_L | Volume for dlPu\_L | `dlPu_L` |  |
| \|\| |  |  | dlPu\_R | Volume for dlPu\_R | `dlPu_R` |  |
| \|\| |  |  | mPFtha\_L | Volume for mPFtha\_L | `mPFtha_L` |  |
| \|\| |  |  | mPFtha\_R | Volume for mPFtha\_R | `mPFtha_R` |  |
| \|\| |  |  | mPMtha\_L | Volume for mPMtha\_L | `mPMtha_L` |  |
| \|\| |  |  | mPMtha\_R | Volume for mPMtha\_R | `mPMtha_R` |  |
| \|\| |  |  | Stha\_L | Volume for Stha\_L | `Stha_L` |  |
| \|\| |  |  | Stha\_R | Volume for Stha\_R | `Stha_R` |  |
| \|\| |  |  | rTtha\_L | Volume for rTtha\_L | `rTtha_L` |  |
| \|\| |  |  | rTtha\_R | Volume for rTtha\_R | `rTtha_R` |  |
| \|\| |  |  | PPtha\_L | Volume for PPtha\_L | `PPtha_L` |  |
| \|\| |  |  | PPtha\_R | Volume for PPtha\_R | `PPtha_R` |  |
| \|\| |  |  | Otha\_L | Volume for Otha\_L | `Otha_L` |  |
| \|\| |  |  | Otha\_R | Volume for Otha\_R | `Otha_R` |  |
| \|\| |  |  | cTtha\_L | Volume for cTtha\_L | `cTtha_L` |  |
| \|\| |  |  | cTtha\_R | Volume for cTtha\_R | `cTtha_R` |  |
| \|\| |  |  | lPFtha\_L | Volume for lPFtha\_L | `lPFtha_L` |  |
| \|\| |  |  | lPFtha\_R | Volume for lPFtha\_R | `lPFtha_R` |  |
| \|\| |  |  | brainstem | Volume for brainstem | `brainstem` |  |
| \|\| |  |  | left\_cerebellum\_cortex | Volume for left\_cerebellum\_cortex | `left_cerebellum_cortex` |  |
| \|\| |  |  | right\_cerebellum\_cortex | Volume for right\_cerebellum\_cortex | `right_cerebellum_cortex` |  |

Close

#### scilus/sf-pediatric Workflow Summary

- this information is collected when the pipeline is started.https://github.com/scilus/sf-pediatric

###### AI Summary

Provider: , model:

Chat with Seqera AI

**Input/output options**

bids\_script
:   `/home/agagnon/.nextflow/assets/scilus/nf-pediatric/bin/BIDSLayout.py`

input
:   `/home/agagnon/scratch/ExamplesMQC/ExampleData/`

multiqc\_title\_global
:   `sf-pediatric MultiQC Global Report`

multiqc\_title\_subject
:   `sf-pediatric MultiQC Subject Report`

outdir
:   `/home/agagnon/scratch/ExamplesMQC/nf-pediatric-0.2.0-Example/`

**Segmentation Options**

fs\_license
:   `/home/agagnon/license.txt`

**Atlases Options**

utils\_folder
:   `/home/agagnon/.nextflow/assets/scilus/nf-pediatric/assets/FS_BN_GL_SF_utils/`

**DWI Preprocessing Options**

dwi\_susceptibility\_readout
:   `0.040`

dwi\_synthstrip\_weights
:   `/home/agagnon/.nextflow/assets/scilus/nf-pediatric/assets/synthstrip.infant.1.pt`

**Pipeline profile**

bundling
:   `true`

connectomics
:   `true`

segmentation
:   `true`

tracking
:   `true`

**Generic options**

lean\_output
:   `true`

trace\_report\_suffix
:   `2025-12-18_19-36-39`

**Core Nextflow options**

configFiles
:   `/home/agagnon/.nextflow/assets/scilus/nf-pediatric/nextflow.config`

containerEngine
:   `apptainer`

launchDir
:   `/scratch/agagnon/ExamplesMQC`

profile
:   `tracking,bundling,segmentation,connectomics,apptainer,slurm`

projectDir
:   `/home/agagnon/.nextflow/assets/scilus/nf-pediatric`

revision
:   `dev`

runName
:   `fervent_noyce`

userName
:   `agagnon`

workDir
:   `/scratch/agagnon/ExamplesMQC/work`

#### Software Versions

Software Versions lists versions of software tools extracted from file contents.

###### AI Summary

Provider: , model:

Chat with Seqera AI

 Copy table

| Group | Software | Version |
| --- | --- | --- |
| ANATTODWI | ants | `2.4.3` |
|  | imagemagick | `6.9.11` |
|  | mrtrix | `3.0.4` |
| BETCROP\_SYNTHBET | synthstrip | `1.5` |
| BET\_DWI | mrtrix | `3.0.5` |
| BRAINNETOMECHILD | freesurfer | `7.4.1` |
|  | scilpy | `1.6` |
| BUNDLE\_FIXELAFD | scilpy | `2.2.1` |
| BUNDLE\_LABELMAP | scilpy | `2.2.1` |
| BUNDLE\_RECOGNIZE | scilpy | `2.2.1` |
| BUNDLE\_STATS | scilpy | `2.2.1` |
| BUNDLE\_UNIFORMIZE | scilpy | `2.2.1` |
| CONCATENATESTATS | pandas | `2.2.3` |
|  | python | `3.10.12` |
| CONNECTIVITY\_AFDFIXEL | scilpy | `1.6` |
| CONNECTIVITY\_DECOMPOSE | scilpy | `2.0.2` |
| CONNECTIVITY\_METRICS | scilpy | `2.0.2` |
| CONNECTIVITY\_VISUALIZE | scilpy | `2.0.2` |
| CONVERT | mrconvert | `3.0.5` |
| CROPB0 | scilpy | `2.0.2` |
| CROPDWI | scilpy | `2.0.2` |
| CROPMASK | scilpy | `2.0.2` |
| DENOISE\_DWI | mrtrix | `3.0.5` |
| DENOISING\_NLMEANS | scilpy | `2.0.2` |
| EXTRACTB0\_RESAMPLE | mrtrix | `3.0.4` |
|  | scilpy | `2.0.2` |
| FASTSEG | fsl | `6.0` |
|  | scilpy | `2.0.2` |
| FILTERING\_COMMIT | scilpy | `1.6` |
| IMAGE\_POWDERAVERAGE | scilpy | `2.0.2` |
| N4\_DWI | ants | `null` |
|  | mrtrix | `3.0.5` |
| NORMALIZE\_DWI | mrtrix | `3.0.4` |
|  | scilpy | `2.0.2` |
| PREPROC\_TOPUP | ants | `2.4.3` |
|  | fsl | `6.0` |
|  | imagemagick | `6.9.11-60` |
|  | mrtrix | `3.0.4` |
|  | scilpy | `2.1.0` |
| RECONALLCLINICAL | freesurfer | `8.0.0` |
| RECONST\_DTIMETRICS | imagemagick | `6.9.11` |
|  | mrtrix | `3.0.4` |
|  | scilpy | `2.1.0` |
| RECONST\_FODF | scilpy | `2.2.0` |
| RECONST\_FRF | scilpy | `2.1.0` |
| REGISTRATION\_ANTS | ants | `2.4.3` |
|  | imagemagick | `null` |
|  | mrtrix | `3.0.4` |
| RESAMPLE\_DWI | scilpy | `2.2.0` |
| RESAMPLE\_MASK | scilpy | `2.2.0` |
| TRACKING\_LOCALTRACKING | scilpy | `2.0.2` |
| TRACKING\_PFTTRACKING | scilpy | `2.0.2` |
| TRACTOGRAM\_MATH | scilpy | `2.2.0` |
| TRACTOGRAM\_REMOVEINVALID | scilpy | `2.2.1` |
| TRACTOGRAM\_RESAMPLE | scilpy | `2.2.1` |
| TRANSFORM\_CENTROIDS | ants | `null` |
|  | scilpy | `2.2.1` |
| TRANSFORM\_LABELS | ants | `2.4.3` |
|  | imagemagick | `null` |
|  | mrtrix | `3.0.4` |
| Workflow | Nextflow | `25.10.2` |
|  | scilus/sf-pediatric | `v0.1.0-gd2c2700` |

#### sf-pediatric Methods Description

Suggested text and references to use when describing pipeline usage within the methods section of a publication.https://github.com/scilus/sf-pediatric

###### AI Summary

Provider: , model:

Chat with Seqera AI

##### Methods

Data was processed using sf-pediatric v0.1.0 of the Sherbrooke Connectivity Imaging Lab (SCIL) ,
utilizing reproducible modules and subworkflows developed by the nf-neuro Team.

The pipeline was executed with Nextflow v25.10.2 (Di Tommaso *et al.*, 2017) using apptainer with the following command:

```
nextflow run scilus/nf-pediatric -r dev --input /home/agagnon/scratch/ExamplesMQC/ExampleData/ --outdir /home/agagnon/scratch/ExamplesMQC/nf-pediatric-0.2.0-Example/ -profile tracking,bundling,segmentation,connectomics,apptainer,slurm --fs_license /home/agagnon/license.txt -resume
```

###### Processing Steps

###### DWI preprocessing

Diffusion weighting imaging (DWI) files were extracted from the input BIDS folder and associated with their corresponding reverse phase-encoded images when available. DWI volumes were denoised using the MP-PCA algorithm (Veraart et al., 2016) implemented in the MRtrix3 toolbox (Tournier et al., 2019). Susceptibility-induced distortions were corrected using FSL's TOPUP (Andersson et al., 2003; Jenkinson et al., 2012) when reverse phase-encoded images were available. Eddy current and motion correction were performed using FSL's EDDY (Andersson & Sotiropoulos, 2016; Jenkinson et al., 2012); maximum framewise displacement was recorded for quality control purposes. Brain extraction was performed by applying the deep learning model SynthStrip (Hoopes et al., 2022) on powdered average images. Pediatric-tailored weights were used for very young subjects where applicable (Kelley et al., 2024). The resulting mask was applied to the DWI volumes. Bias field correction was applied using the N4 algorithm (Tustison et al., 2010) from the ANTs toolbox (Tustison et al., 2021) using a b-spline knot per voxel of 8 and a shrink factor of 4. DWI volumes were normalized using the mean B0 intensity within white matter (FA > 0.4) using MRtrix3 (Tournier et al., 2019). Preprocessed DWI volumes were resampled to an isotropic voxel size of 1 mm.

###### Anatomical preprocessing

Anatomical T1w and/or T2w images were denoised using the Non-Local Means algorithm (Coupe et al., 2008) as implemented in the DIPY toolbox (Garyfallidis et al., 2014). Bias field correction was applied using the N4 algorithm (Tustison et al., 2010) from the ANTs toolbox (Tustison et al., 2021) using a b-spline knot per voxel of 8 and a shrink factor of 4 for the T1w and a b-spline knot per voxel of 8 and a shrink factor of 4 for the T2w image. Anatomical images were resampled to an isotropic voxel size of 1 mm for the T1w and 1 mm for the T2w image. Brain extraction was performed using SynthStrip (Hoopes et al., 2022); pediatric-tailored weights were used for very young subjects where applicable (Kelley et al., 2024). If both T1w and T2w images were available, they were registered using ANTs (Tustison et al., 2021) using an affine transform.

###### Diffusion Tensor Imaging (DTI)

Diffusion tensor imaging (DTI) models were fitted on the processed volume using the scilpy toolbox (Renauld et al., 2025); fractional anisotropy (FA), axial diffusivity (AD), radial diffusivity (RD), mean diffusivity (MD), mode of anisotropy, and color-coded FA maps were generated. DTI fitting used all available shells under the maximum b-value of 1600 s/mm².

###### Fiber Orientation Distribution Function (fODF)

Fiber orientation distribution functions (fODF) were computed using the scilpy toolbox (Renauld et al., 2025) using the single-shell single-tissue method on the all available shells over the minimum b-value of 700 s/mm². fODF were computed using a maximum spherical harmonic order of 8 in basis descoteaux07. Fiber response functions were estimated based on normative curves of the brain's diffusivities through the developmental age-range as described in Gagnon et al. 2025.

###### Registration to DWI space

Anatomical images were registered to the preprocessed DWI space using ANTs (Tustison et al., 2021). For younger participants (< 2.5 years old), the T2w image, if available, was preferred for registration due to better tissue contrast. If used, the T2w image was registered using non-linear methods using the mean diffusivity map and B0 image as targets. For older participants or if only T1w images were available, the T1w image was registered using non-linear methods with the FA map and B0 image as targets.

###### Tissue segmentation

Tissue segmentation into white matter, grey matter, and cerebrospinal fluid was performed on the anatomical images registered to DWI space. For younger participants (< 2.5 years old), segmentation was performed by registering age-matched templates from the UNC/UMN Baby Connectome Project (Chen et al., 2022). Briefly, templates closest to the participant's age were non-linearly registered to the participant's anatomical images using ANTs (Tustison et al., 2021), and the resulting transforms were applied to the corresponding tissue probability maps. The resulting maps were then thresholded to generate binary masks for each tissue type. For older participants, tissue segmentation was performed using the FAST algorithm from FSL (Zhang et al., 2001; Jenkinson et al., 2012). Similarly to younger participants, resulting probability maps were thresholded to obtain binary masks.

###### Tractography

Whole-brain tractography was performed using the scilpy toolbox (Renauld et al., 2025). Particle Filter Tracking (PFT) was used to leverage anatomical priors from the tissue segmentation to improve streamline generation (Girard et al., 2014). Tracking seeds were randomly placed within the white matter mask with a density of 10 seeds per voxel. Streamlines were propagated using a probabilistic algorithm with a step size of 0.5 mm, a maximum angle between steps of 20°, a minimum length of 20 mm, and a maximum length of 200 mm. Local tracking was performed using a probabilistic algorithm using 10 seeds per voxel. The seeding mask was defined as the white matter mask. Similarly, the tracking mask, in which tracking is allowed, was defined as the white matter mask. Streamlines were propagated with a step size of 0.5 mm, a maximum angle between steps of 20°, a minimum length of 20 mm, and a maximum length of 200 mm. The resulting two tractograms from both methods were then concatenated to form the final whole-brain tractogram.

###### Bundle segmentation

The closest age-matched white matter atlas (neonates, 3 months, 6 months, 12 months, 24 months or children) was registered into subject-space using an affine transformation. Whole-brain tractograms were segmented using BundleSeg from the scilpy toolbox (St-Onge et al., 2023; Renauld et al., 2025) with a minimal vote ratio of 0.5, an outlier threshold of 0.6, and the euclidean distance. Extracted bundles were then filtered to remove invalid streamlines, single point streamlines, and overlapping points. Then, fixel-based apparent fiber density was computed for each bundle (Raffelt et al., 2017).

###### Tractometry

Atlas' centroids were registered into subject-space using an affine transformation. The centroids were then resampled to 5 points, enabling the derivation of per point metrics. Metric derived per bundle or per point were weighted based on the number of streamline passing through the voxel. This reduces the impact of spurious streamlines on final metric value. For each bundle, multiple metrics were extracted: length, statistic for each endpoint, mean (standard deviation), volume, and streamline count. For each point per bundle (5 points), the following metric were extracted: volume, and mean (standard deviation). Final segmented bundles were colored per point using the jet colormap (affects only the visualisation).

###### Cortical and sub-cortical segmentation

Cortical and subcortical segmentation was performed using recon-all-clinical from Freesurfer (Fischl, 2012; Billot et al., 2023; Iglesias et al., 2023) on the T1w anatomical images. Following segmentation, the Brainnetome Child Atlas (Li et al., 2023) was mapped in subject-space using surface-based registration methods from FreeSurfer (Fischl, 2012) and then converted into voxel labels. For each parcels, volume, surface area, and cortical thickness were measured and outputted in tab-separated value files. For younger participants (< 3 months old), cortical and sub-cortical segmentation was performed using the M-CRIB-S pipeline (Adamson et al., 2020). Younger participants were segmented the Desikan-Killiany (Desikan et al., 2006, Adamson et al., 2020). Following segmentation, volume, surface area, and cortical thickness were measured for each parcel and outputted in tab-separated value files.

###### Connectomics

Structural connectivity matrices were generated using the scilpy toolbox (Renauld et al., 2025) based on the Brainnetome Child Atlas (Li et al., 2023) or the Desikan-Killiany atlas (Desikan et al., 2006) depending on the participant's age. For each participant, labels in anatomical space were first registered in diffusion space using the already computed transformations with a nearest neighbor interpolation method. Then, the final tractogram was decomposed into individual connections by extracting each streamline connecting a pair of parcels. Streamlines shorter than 20 mm or longer than 200 mm were discarded. Loops were removed. Hierarchical QuickBundles was used to remove outliers using a threshold of 0.6. Curvature-based filtering was applied to remove streamlines with sharp curves using a maximum angle of 330.0° over 10.0 mm. To mitigate the risk of false-positive connections, COMMIT (Daducci et al., 2015) was applied to the tractogram using the stick, zeppelin, and ball model to optimize the fit between the tractogram and the diffusion data. Diffusivity parameters for COMMIT were set based on age-specific normative values as described in Gagnon et al. 2025. Using COMMIT2 (Schiavi et al., 2020) with a clustering prior strength of 0.001, the contribution of each streamline to the diffusion signal was evaluated and streamlines with zero contribution were removed from the tractogram to further reduce false-positive connections. To obtain the fODF amplitude specific to each connection, fixel-based apparent fiber density was computed for each extracted connection (Raffelt et al., 2017). Finally, structural connectivity matrices were generated by computing, for each pair of parcels, the number of streamlines, the mean streamline length, and the mean FA, AD, RD, MD, total apparent fiber density, number of fiber orientation, and fixel-based apparent fiber density.

###### Notes:

- If available, make sure to update the text to include the Zenodo DOI of version of the pipeline used.
- The command above does not include parameters contained in any configs or profiles that may have been used. Ensure the config file is also uploaded with your publication!
- You should also cite all software used within this run. Check the "Software Versions" of this report to get version information.

**MultiQC v1.32** 
- Written by Phil Ewels, available on
GitHub.

This report uses Plotly,
jQuery,
jQuery UI,
Bootstrap and
FileSaver.js.

##### Plot Table Data

Select Column

Select Column

Please select two table columns.

Close

##### Regex Help

Toolbox search strings can behave as regular expressions (regexes). Click a button below to see an example of
it in action. Try modifying them yourself in the text box.

`^` (start of string)

`$` (end of string)

`[]` (character choice)

`\d` (shorthand for `[0-9]`)

`\w` (shorthand for `[0-9a-zA-Z_]`)
`.` (any character)

`\.` (literal full stop)

`()` `|` (group / separator)

`*` (prev char 0 or more)

`+` (prev char 1 or more)

`?` (prev char 0 or 1)

`{}` (char num times)

`{,}` (count range)

```
samp_1
samp_1_edited
samp_2
samp_2_edited
samp_3
samp_3_edited
prepended_samp_1
tmp_samp_1_edited
tmpp_samp_1_edited
tmppp_samp_1_edited
#samp_1_edited.tmp
samp_11
samp_11111
```

See regex101.com for a more heavy duty testing suite.

Close
