## Supplementary for "sf-pediatric: A robust and age-adaptable end-to-end pipeline for pediatric diffusion MRI": SupplementaryFile3.html

sf-pediatric MultiQC Subject Report: MultiQC Report


Loading report..

v1.32


Theme

- Light
- Dark
- Auto

### sf-pediatric MultiQC Subject Report

Highlight

 Rename

 Show / Hide


AI Analysis

 Export

 Settings

 Citations

 About

- Shell QC
- Topup QC
- Framewise Displacement
- Eddy QC
- Metrics QC
- Registration of anatomical image in DWI space QC
- Tissue Segmentation QC
- Tracking Coverage
- Labels QC
- scilus/sf-pediatric Workflow Summary
- Software Versions
- sf-pediatric Methods Description

MultiQC is developed by Seqera.

Scroll to top

### sf-pediatric MultiQC Subject Report

A modular tool to aggregate results from bioinformatics analyses across many samples into a single report.

> This report has been generated by the scilus/sf-pediatric analysis pipeline.

###### JavaScript Disabled

MultiQC reports use JavaScript for plots and toolbox functions. It looks like you have JavaScript disabled in your
web browser. Please note that many of the report functions will not work as intended.

Loading report..

Report
generated on 2025-12-19, 17:20 EST
based on data in:
`/tmp/nxf.ASWOO6LtLE`

Summarize report

Copy report prompt

**Welcome!** Not sure where to start?
Watch a tutorial video
*(6:06)*

don't show again

###### Report AI Summary

More details…

Provider: , model:

Chat with Seqera AI

#### Shell QC

This section contains QC images for the diffusion shell, displaying
the distribution of b-vectors on the sphere. This visualization ensures
that the diffusion-weighted gradient directions are correctly assigned
and evenly distributed, which is critical for accurate modeling of
diffusion properties. To assess the quality of the b-vectors, check that
the gradient directions form a uniform and symmetric distribution on
the sphere, corresponding to the expected acquisition scheme (e.g.,
single-shell, multi-shell). Look for missing or misaligned directions,
which may indicate acquisition or preprocessing errors.

###### AI Summary

Provider: , model:

Chat with Seqera AI

#### Topup QC

This section contains QC images for TOPUP correction, which is used to correct
susceptibility-induced distortions by leveraging pairs of images acquired with
opposite phase-encoding directions. The animation shows a before-and-after
comparison of the correction. To assess the quality of the correction, focus
on regions most affected by susceptibility distortions, such as the frontal lobe
and the cerebellum/spinal cord area. Look for improved alignment of anatomical
structures, reduced stretching or compression, and better symmetry between the
corrected images. Additionally, ensure that the correction does not introduce
new artifacts or excessive blurring.

###### AI Summary

Provider: , model:

Chat with Seqera AI

#### Framewise Displacement

Assessment of subject motion during acquisition using the framewise displacement (FD) metric calculated by FSL's `eddy` tool for quality control.https://github.com/nf-neuro/MultiQC\_neuroimaging

##### Framewise Displacement

Framewise displacement across volumes. Each point represents the FD for a volume compared to the previous volume. High spikes in FD may indicate excessive motion during scanning. While `eddy` attempts to correct for motion, users should be cautious when interpreting data from subjects with high FD values. Green: <0.8mm, Yellow: 0.8-2.0mm, Red: >2.0mm

###### AI Summary

Provider: , model:

Chat with Seqera AI

Export...

Copy prompt


Summarize plot

Created with MultiQC

#### Eddy QC

This section contains QC images for EDDY correction, which is used to
correct distortions and motion artifacts. The animation shows a
before-and-after comparison of the correction. To assess the quality of
the correction, focus on areas prone to distortions, such as the frontal
lobe and the cerebellum/spinal cord region. Look for reduced geometric warping,
improved alignment with anatomical boundaries, and correction of
eddy-current-induced displacements. Additionally, ensure that no new
artifacts or excessive blurring are introduced in the corrected images.

###### AI Summary

Provider: , model:

Chat with Seqera AI

#### Metrics QC

This section contains QC images for diffusion tensor imaging
(DTI) metric maps and FODF metric maps, including fractional anisotropy
(FA), mean diffusivity (MD), RGB FA, and number of fiber orientation (NuFO).
To assess the quality of the DTI metrics, ensure that FA highlights major
white matter tracts with expected high values (e.g., corpus callosum,
corticospinal tract) and that MD, RGB FA, and NuFO values follow known
anatomical distributions. Look for smooth, artifact-free maps, avoiding
unexpected signal dropout, distortions, or excessive noise. Pay special
attention to regions prone to artifacts, such as near the ventricles or
areas affected by eddy currents and susceptibility distortions.

###### AI Summary

Provider: , model:

Chat with Seqera AI

#### Registration of anatomical image in DWI space QC

This section contains QC images for the registration of the T1-weighted
structural image to diffusion space, using the b0 image as a reference.
This step ensures accurate alignment between structural and diffusion
data. To assess the quality of the registration, check for good alignment
of anatomical structures. Specifically, look for proper overlay of cortical
and subcortical structures without excessive warping, stretching,
or misalignment. Pay particular attention to boundaries such as the
ventricles and major white matter tracts, ensuring they match well
between the T1-weighted and b0 images. If misalignment is present,
consider flagging this specific subject for further review.

###### AI Summary

Provider: , model:

Chat with Seqera AI

#### Tissue Segmentation QC

This section contains images for visual quality control of tissue segmentation,
where different tissue classes are color-coded: red for white matter (WM),
green for gray matter (GM), and blue for cerebrospinal fluid (CSF).
This segmentation is typically derived from a T1-weighted structural image
(if pediatric) or T2-weighted structural image (infant) and is crucial for
downstream analyses such as tractography. To assess the quality of the
segmentation, ensure that each tissue class is well-defined and corresponds
accurately to expected anatomical regions. White matter should be
predominantly within deep brain structures and major tracts,
gray matter should outline the cortex and subcortical nuclei, and
CSF should primarily appear in the ventricles and surrounding sulci.
Check for misclassifications, such as CSF incorrectly assigned within
the brain or gray/white matter boundaries appearing blurred or inconsistent.
If artifacts or misclassifications are present, consider flagging this specific
for further review.

###### AI Summary

Provider: , model:

Chat with Seqera AI

#### Tracking Coverage

This section contains QC images of tractography coverage, shown as an
overlay of the tract density image (pink) with the WM mask (green).
This visualization assesses how well
the reconstructed streamlines cover the white matter mask used for
tracking, with higher coverage generally indicating better results.
To evaluate tracking quality, check that the tract density image spans
the full extent of the white matter mask, ensuring that major pathways
(e.g., corpus callosum, corticospinal tract) are well-represented.
Ideally, the coverage should be uniform across the mask, with minimal
gaps or missing regions. Areas of low coverage may indicate poor
tracking due to insufficient diffusion signal, masking errors, or
overly restrictive tracking parameters. Conversely, excessive streamlines
outside the mask may suggest tracking leakage into non-white matter
regions, requiring further filtering.

###### AI Summary

Provider: , model:

Chat with Seqera AI

#### Labels QC

This section contains QC images for the segmentation of
cortical and subcortical structures, displayed as an overlay of
anatomical labels. These labels are derived from structural MRI
and serve as key regions of interest for connectivity analyses and
volumetric measurements. To assess segmentation accuracy, verify
that each label correctly corresponds to its respective anatomical
structure. Cortical labels should align with gyri and follow natural
sulcal boundaries, while subcortical labels should fit well within deep
gray matter structures such as the thalamus and basal ganglia. Check for
misalignments, incorrect label assignments, or excessive partial
volume effects. If discrepancies are found, consider refining
registration or segmentation parameters.
